# Accurate Multiple Sequence Alignment of Ultramassive Genome Sets

**DOI:** 10.1101/2024.09.22.613454

**Authors:** Xiao Lai, Haixin Luan, Pu Tian

## Abstract

With ever increasing sequencing efficiency, there is a pressing need to tackle a presently intractable task of accurate multiple sequence alignment (MSA) for ultramassive genome sets of millions and beyond. Additionally, efficient graph and probabilistic representations for downstream analysis are in dire lack. In light of these challenges, we develop a set of essentially linearly scalable algorithms, including that for constructing directed acyclic graphs, for training tiled profile hidden Markov models and for conducting alignment on such graphs. The power of these algorithms is demonstrated by both significantly improved accuracy and tremendous acceleration of SAR-CoV-2 MSA by *three observed to five projected orders of magnitude* for genome set sizes ranging from 40,000 to 4 million when compared with widely utilized MAFFT. Future application to other viral species and extension to more complex genomes will prove this algorithm set as a cornerstone for the coming era of ultramassive genome sets.

## INTRODUCTION

The SARS-CoV-2 pandemic brought us a genome dataset amount to more than 20 million (https://gisaid.org)^1^.This provides a potential opportunity for investigating evolution of a virus with unprecedented resolution^2^. With ever increasing efficiency and improving accuracy of sequencing technology, ultramassive genome sets of more species are likely to be generated. For example, the number of publicly available human genomes is approaching 1 million (https://www.ukbiobank.ac.uk,https://allofus.nih.gov). and is projected to reach half a in 2030 (https://www.ga4gh.org/news_item/ga4gh-releases-2018-strategic-roadmap/). Somatic mutation studies are expected to generate large sets of single cell genomes for many individuals^3^, and extension of such sequencing investigations to other important species is warranted. Therefore we are inevitably entering an upcoming era of ultramassive genome sets. To reliably extract valuable information on evolution and molecular interactions buried in genome diversity and correlations among different segments there in, it is essential to perform accurate multiple sequence alignment (MSA), which is unfortunately computationally intractable for ultra-massive genome sets with presently available algorithms.

The goal of MSA computation is to align residues of the same evolutional and/or (functional, structural) origin from different sequences to the same column as illustrated in Figure 1(a,b). As real evolutional paths of biological sequences are unknowable, the most widely utilized strategy is to find the maximum likelihood solution as an effective approximation, rigorous calculation of which is still intractable for hundreds of or more sequences as its computational cost scales exponentially with the number of sequences (*n*). There has been a long history of approximate MSA algorithm development^4–10^, which are based on dynamic programming^5,9^, hidden Markov models (HMMs)^4,11–13^, or some form of their mixtures^14,15^. The grand challenge of MSA algorithmic complexity was well documented theoretically^16,17^. Presently, consistency based algorithms^18^ are considered to be sufficiently accurate for practical purposes, it has a computational complexity of ∼ 𝒪 (*n*^3^*L*^2^) (Simultaneous alignment of *p* sequences results in ∼ *n*^*p*^ complexity), which amount to at least 3.8 × 10^30^ float-point operations for the particular case of more than 4 million high quality SARS-CoV-2 genomes, and takes more than 1, 000 years to complete on a exa-scale supercomputer. MAFFT^19^ is widely utilized^20–25^ for its much higher efficiency and acceptable accuracy. Regressive alignment^26^ utilizes present mainstream methods (e.g. MAFFT) with a dividing and conquer strategy to speed up MSA calculation. Recent FM-index alignment^27,28^ made some progress by removing some of tremendous repetitive computation, which is ubiquitous for all present algorithms when applied on similar sequences (e.g. SARS-CoV-2 or human genome sets, see Supplemental Information (SI) S 1.2 and Figure S1, Figure S2 for details). Unfortunately, performance of all above algorithms are far from practically feasible for such ultramassive genome sets. Halign4^29^ is an extremely efficient algorithm for millions of sequences, but is not sufficiently accurate when high level of accuracy is essential. Besides difficulty in obtaining accurate MSA, we are in dire lack of efficient and succinct graph and probabilistic representations for downstream tasks of viral genome analysis. These tasks include but are not limited to comparative genomics, rapid (joint) variant calling for large data sets, and monitoring and accurate phylogenetic characterization of continuously arising new genomes in real time.

**Figure 1:**
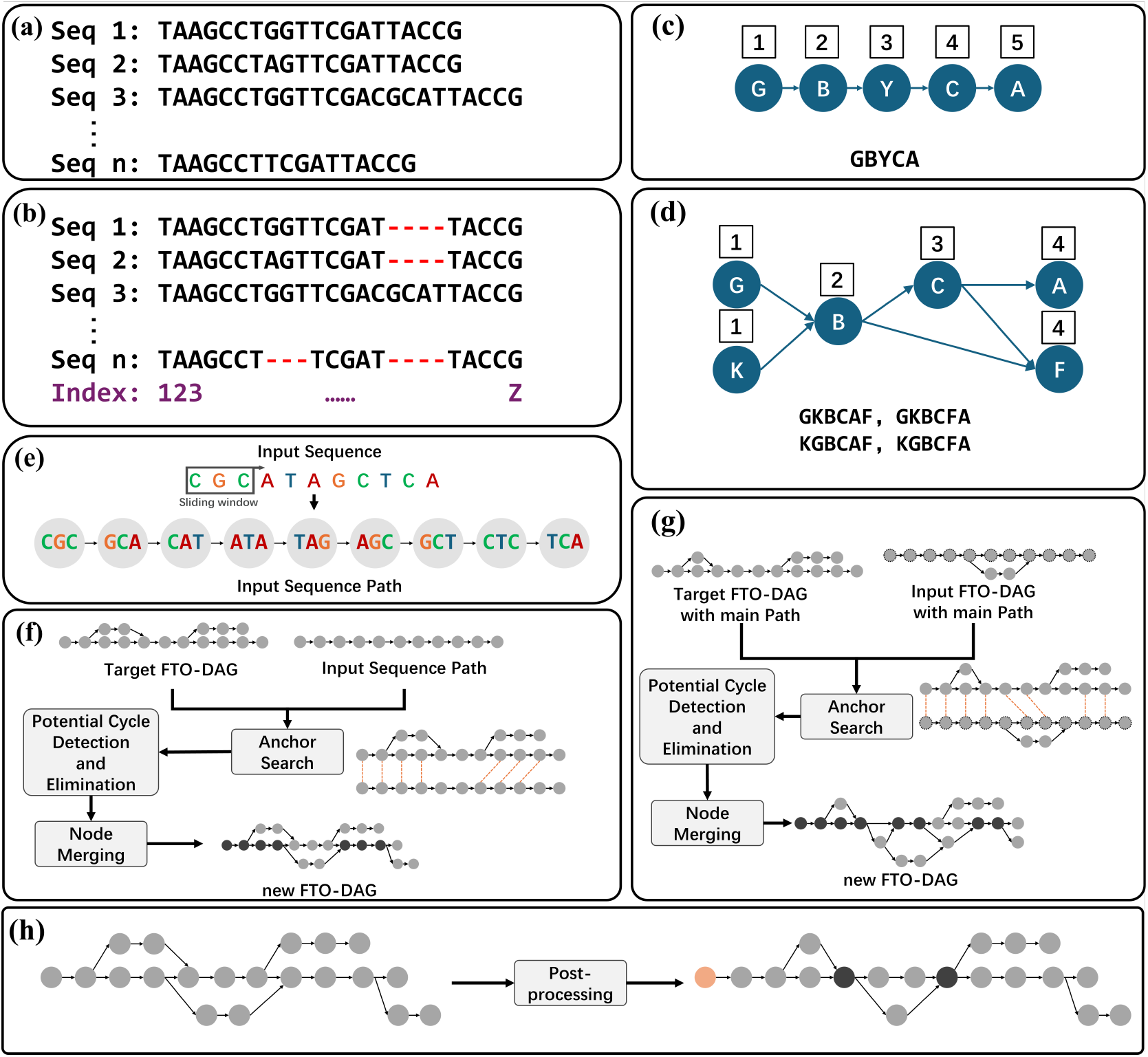
Schematic illustration of MSA, topological order (TO) and the FTO-DAG Construction Workflow. (a) A set of *n* unaligned sequences. (b) The sequences after alignment, arranged into *Z* columns. The process aims to maximize a score of a given metric by optimally placing gaps (red hyphens). (c) The topological order (TO) of a linear path of nodes determines a unique order of it comprising nodes. The numbers in squares above the nodes represent their topological coordinates (TC). (d) For a DAG, TO only requires that parent nodes precede child nodes, making the order non-unique. This example shows all 12 possible orderings of nodes conform to TO. Again, the numbers in squares above the nodes represent their TC. TCs on the direction of a DAG path have the following property: i) Start nodes (with no parent) have 1 as their TC. ii) A node with parent nodes have the maximum value of TCs from its parent nodes plus 1. The reverse TC is defined accordingly as: i)End nodes (with no child) have 1 as their reverse TC, ii) A node with child(ren) node(s) have the maximum value of TCs from its child(ren) node(s) plus one. So proper (reverse) TCs are always monotonically increase on the (reverse) direction of all DAG paths, and violation of such property is called TC conflict. (e) An input sequence is transformed into an Input Sequence Path via a sliding window. (f) Incremental integration of an input sequence path into a target FTO-DAG. (g) Merging of an input FTO-DAG into a target FTO-DAG. As described in the text and illustrated here, three steps: i) Anchor search, ii) Potential cycle detection and elimination, and iii) Node merging are shared for both incremental integration and graph merging. (h) Schematic of post-processing graph optimization, showing the initial graph (left) and the more compact final graph (right) after optimization.

We set to tackle these challenges by developing a large set of new algorithms, including that for Fragment Topological Order (FTO) guided construction of Directed Acyclic Graphs (DAGs) (with topological order (TO) illustrated in Figure 1(c,d), and graph construction in Figure 1(e,f,g)), for training of Tiled Profile Hidden Markov Models (TPHMM) on DAGs and aligning with graph adapted Viterbi algorithm (see Figure 3 and Figure S7, Figure S8, Figure S9). Since these algorithms hinge on the DAG data structure, the graphs are termed FTO-DAG, the probabilistic models DAG-TPHMM, the modified Viterbi algorithm DAG-Viterbi, and the whole algorithm package DAG-align. The DAG-align dramatically accelerates MSA by a combinative effect of: removing essentially all repetition of computation engendered by many present alignment algorithms on similar sequences (see Figure S2) with FTO-DAG; ii) removing highly unlikely state position computation in conventional profile HMM (Figure S5) with tiling (Figure S7).

## RESULTS

### The DAG-align algorithm set

#### The FTO-DAG graph construction algorithm set

Computation repetition takes vast majority of resources in conventional MSA of similar sequences (see Figure S2 and accompanying text in Supplemental Information (SI)). Brute force elimination of such repetition may result in excessive caching or losing contextual dependency. While De Bruijn Graph^30^ may effectively compress identical sequence fragments without above mentioned issues, its inherent cyclic structure is not readily compatible with MSA. To address these challenges, we develop a new set of algorithms to accomplish efficient construction of FTO-DAG, which simultaneously realizes effective removal of computation repetition and compatibility with alignment. The overall pipeline is schematically illustrated in Figure 1(e,f,g) and briefed below (see details in STAR Methods, FTO-DAG Construction Algorithm Set.). To realize efficient parallelization for ultra-massive genome sets (see Figure S6 for illustration), FTO-DAG construction is divided into subgraph construction and iterative merging stages. The subgraph construction handles raw biological sequences from each sequence subset. Firstly, a sliding window is utilized to slice a sequence into ordered fragments (Figure 1e), which are then organized into a linear input sequence path. This path is subsequently utilized to initialize an empty target FTO-DAG or incrementally expand an existing one (Figure 1f). The merging stage is for fusing an existing input FTO-DAG (Figure 1g) into a target FTO-DAG. In both stages, the integration process follows a rigorous three-phase workflow to ensure the graph’s integrity and acyclic nature.

##### Phase 1: Anchor Search

For each input node of an input sequence path (or each main path node of an input FTO-DAG), the algorithm searches in the target FTO-DAG for an anchor, which is a node for the input node to merge into. The search may fail for some nodes, and the rate of success is dependent upon similarity of sequences represented by the input sequence path (or FTO-DAG) and the target FTO-DAG. An anchor must meet the following three criteria:

- Sequence Match: The sequence fragment contained in an anchor must be identical to that of the input node.
- Node Type Match: A start/end/intermediate node (see STAR Methods, FTO-DAG Construction Algorithm Set: Definitions) only search for nodes of the same type in the target FTO-DAG as its anchor.
- Highest Weight: Among all candidate nodes with a matching sequence fragment, the anchor must have the highest weight value (representing its number of historical merges).
- Uniqueness: No other nodes with an identical sequence and the same highest weight exist, thus avoids ambiguity.

Notably, for the merging of an input FTO-DAG, the anchor search is performed only for its main path (see STAR Methods, FTO-DAG Construction Algorithm Set: Definitions) nodes in the whole target FTO-DAG to ensure the graph’s key structural elements are prioritized for accurate alignment.

##### Phase 2: Potential cycle detection and elimination

Direct merging of every initial matching input nodes into its identified anchor may introduce cycles to the target FTO-DAG (see Figure 2(a) bottom), thus violating its acyclic property. An active “identification-elimination” procedure is performed to prevent formation of potential cycles (see Figure 2(b)) as described below. The potential cycle elimination step is performed when TC conflicts were detected, the initial matching input nodes and their anchors will be directly utilized for merging otherwise.

**Figure 2:**
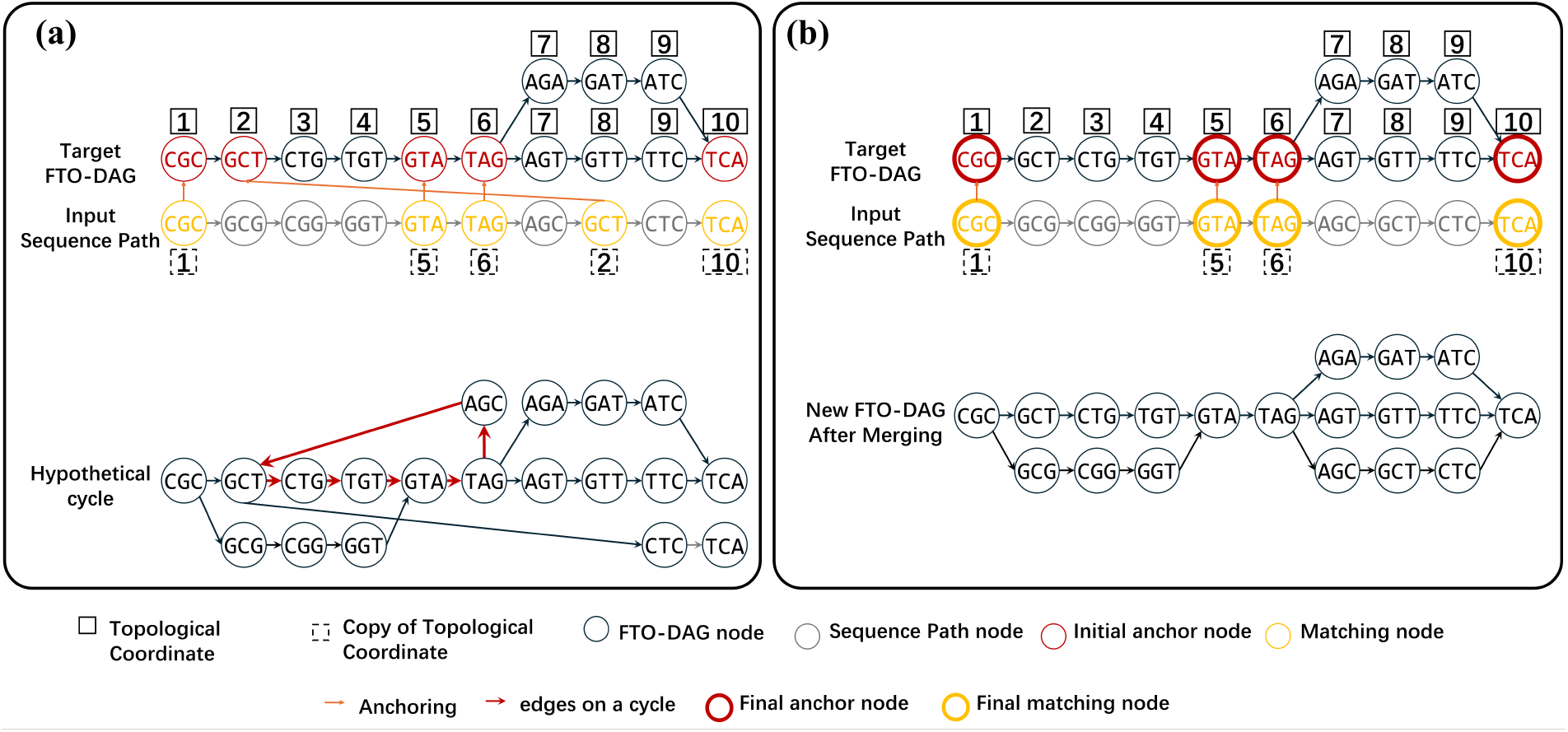
Schematic illustration of potential cycle detection and elimination process. (a) Top: a scenario where in the initial matching input nodes, TC conflict occurs with the extracted TC sequence [1, 5, 6, 2, 10], violating monotonic increase property of TO on a directed path. Bottom: a hypothetical cycle with its edges shown as red arrows would have been formed if direct merging of these initial matching input nodes to their anchors were performed. (b) Top: the final TC conflict free matching input nodes and their corresponding final anchors returned from the dynamic programming algorithm (see Algorithm 4 in STAR Methods) searching for the longest and maximum-weight-sum TC conflict free subset. Bottom: a proper target FTO-DAG after integration based on TC conflict free matching input nodes and their anchors.

- Identification of potential cycle(s) (Figure 2(a) Top) with TC conflict (defined in caption of Figure 1) in two steps for FTO-DAG expansion or merging:
  – After the initial anchor search and before the merging, the TC of each anchor in the target FTO-DAG is copied and assigned to its matching input node.
  – Newly assigned TCs for matching input nodes are extracted to form a TC sequence in the direction specified by directed edges in the input sequence path (or FTO-DAG), and existence of TC conflicts are checked.
- Potential cycle elimination. Upon detecting of TC conflict(s), a dynamic programming algorithm for finding the longest and maximum-cumulative-weight TC conflict free subsequence (see Algorithm 4 in STAR Methods) is called on the extracted TC sequence, and returns a final matching input node set and corresponding anchor set.

##### Phase 3: Node Merging

With the conflict-free matching input nodes and anchors are established, the merging step proceeds as follows:

- For each matching input node, all its information (such as weight, path relationships, etc.) is completely transferred and aggregated onto its anchor in the target FTO-DAG.
- For each input node without an anchor, they are added directly to the target graph as a new node, with connections to relevant matching input node in the input sequence path (or FTO-DAG) transferred to the corresponding anchor in the target FTO-DAG.

##### Post-processing and Compression Evaluation of the Graph

After the initial graph construction, the following optimization may be performed to further reduce redundancy and increase compression ratio (see Equation (1)).

##### Graph Resolution Adjustment

The resolution of a graph (i.e., the length of the sequence fragment contained in each node) can be adjusted post-construction (only shortening is supported at present). This process (illustrated in Figure S4) involves two steps:

- Sequence Truncation: Save a copy of all start nodes, subsequently the sequence fragment is truncated for each node to retain only its final 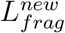 bases.
- Start Node Completion: The truncated part of all start nodes are added back as new start nodes and 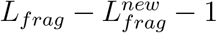 of its successor nodes to maintain integrity of each sequence.

Notably, when 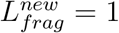, this process is referred to as the atomization of a FTO-DAG.

##### Secondary node merging

To further reduce redundancy, a topology-based node merging may be carried out. Specifically, all pairs of nodes in different paths sharing both identical sequence fragment and TC are merged (see details in STAR Methods: FTO-DAG Construction Algorithm Set, Detailed workflow of FTO-DAG construction). This maximizes the compression rate while maintaining the acyclic property. To quantify the graph’s compression efficiency, we define the compression ratio *η* as follows:

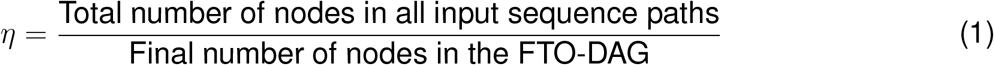

A higher compression ratio indicates a more effective elimination of sequence redundancy, and consequently computation repetition in alignment by the FTO-DAG. As shown in Figure S13, *η* increases significantly with increasing genome set sizes, which is the fundamental basis of the DAG-align’s high scalability (Figure S14) with respect to the sequence number.

#### The DAG-TPHMM and DAG-Viterbi algorithm sets

The Profile Hidden Markov Model (PHMM)^4,31^ is a foundational probabilistic model in bioinformatics. Besides a start and an end state position for initiating and finishing the model, it has a specified number of regular state positions, each of which may have one of the following three states: Match (*M*), Insertion (*I*), and Deletion (*D*). Possible transitions between states of the same and/or neighboring state positions are illustrated in Figure S5. Conventional PHMM are constructed for linear sequences. To deal with branching/gathering structures in a DAG, we created virtual child/parent node (see Figure 3(e,f)) to convert complex “many-to-one”/”one-to-many” dependencies into “one-to-one” pseudo-linear relationships that are compatible with standard recurrence formulas. Subsequently probability summation (Equations (6), (10) and (18)) is utilized to realize forward/backward of branching/gathering structures in training and decoding stages (see STAR Methods, DAG-TPHMM and DAG-Viterbi Algorithm sets). The overall pipeline is illustrated in Figure 3(a,b,c,d) and described below:

**Figure 3:**
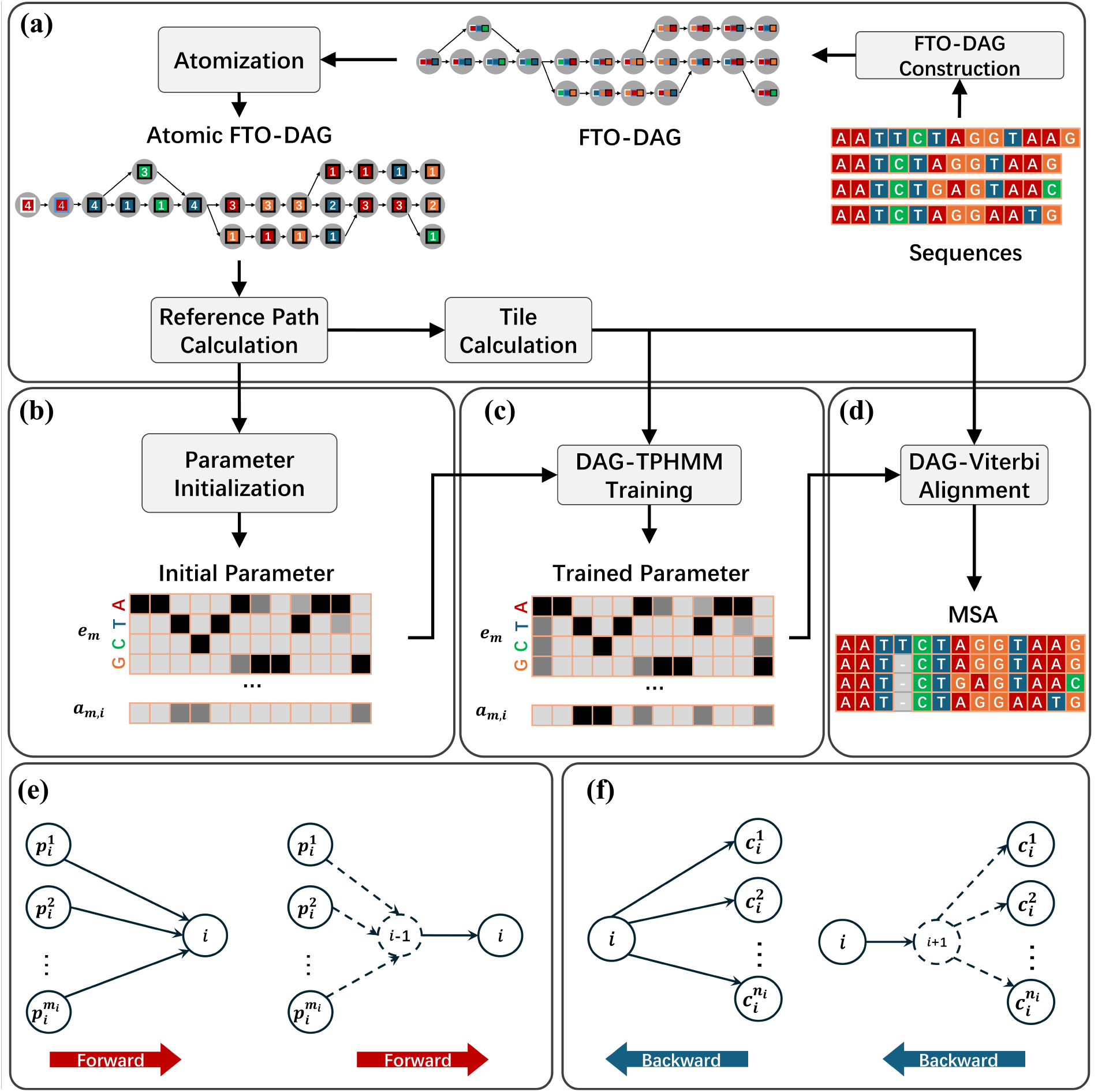
Schematic representation of the algorithmic framework for DAG-align. Details of each step are explained in STAR Methods: DAG-TPHMM Algorithm Set (a) Fragment based construction of FTO-DAG from input sequences followed by atomization, reference path and tile calculation for preparation of DAG-TPHMM. (b) TPHMM parameter initialization based on global graph information. (c) Parameter training on the global FTO-DAG using the Baum-Welch algorithm. (d) Decoding the optimal path and generation of the final MSA using the DAG-Viterbi algorithm. (e) Illustration of a virtual node summing for multiple parent nodes. Left: A forward computation unit (*i*) and its parent nodes. Right: Constructing the virtual predecessor node (*i* − 1) for the forward computation. The probability summation is shown in Equation (6) in STAR Methods. (f) Illustration of a virtual node summing for multiple children nodes. Left: A backward computation unit (*i*) and its child nodes. Right: Constructing the virtual successor node (*i* + 1) for the backward computation. The probability summation is shown in Equation (10) in STAR Methods.

##### Preprocessing: Atomization of the FTO-DAG

As alignment are formulated for single bases, the constructed FTO-DAG is first atomized (see Figure 3(a), also illustrated in Figure S4). This process decomposes each node, which originally represents a sequence fragment, into atomic nodes representing single bases, while preserving all topological relationships.

##### The Alignment Pipeline

It consists of five key stages briefed below (detailed in STAR Methods, DAG-TPHMM Algorithm Set):

- State position establishment through reference path calculation. After pruning and coarsening of the atomized FTO-DAG, the main path is calculated and defined as the reference path. Each node on the reference path corresponds to a regular state position.
- Parameter Initialization. We analyze the global sequence conservation information embedded within the FTO-DAG to construct an initial set of parameters, including emission and transition probabilities for all state positions (Figure 3(b), Figure S8).
- Tile Calculation. Each node in the FTO-DAG is assigned a calculated tile range, and only state positions within that range is considered in forward/backward computation of DAG-TPHMM (Figure S7).
- Parameter Training. One round of Baum-Welch algorithm is applied to optimize the TPHMM parameters on the atomized and tiled FTO-DAG (Figure 3(c)).
- Alignment Decoding. After the training is complete, the DAG-Viterbi algorithm is used to decode the optimal hidden state path on the FTO-DAG, which is then translated into the final MSA (see Figure 3(d),and Figure S9).

#### Parallelization Strategies

Different strategies are utilized for FTO-DAG construction, training and decoding and are briefed below:

- In FTO-DAG construction, parallel computation is realized through partition of the total dataset into subset, followed by parallel construction of subgraphs from each of the subsets, and finalized by 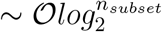 rounds of iterative two-to-one merging operations. This process is schematically illustrated in Figure S6(a). The scaling of the FTO-DAG construction for various numbers of processors is shown in Figure S10.
- In DAG-TPHMM training, parallel computation is controlled by the topological order on the FTO-DAG. Specifically, the global FTO-DAG experience a graph coarsening operation where each maximal unbranched path is converted into a super node as a task unit (Figure S6(b) top). Then a graph traversal strategy that strictly adheres to topological order is used to schedule these computational task units, thereby effectively utilizing multi-core computing resources (see Figure S6(b) bottom).
- In DAG-Viterbi decoding, parallelization is realized at two different levels. Firstly, by mapping the global FTO-DAG reference path into each subgraphs, the decoding computation may be performed independently and in parallel on each subgraph(see Figure S6(c)). Secondly, on each subgraph, the same TO based computation task queue as in training may be constructed and utilized when necessary (see Figure S6(b)) except that the reverse TO is used for guidance. The scaling of the decoding process for various number of processors is shown in Figure S11.

### Evaluation of Alignment Efficiency and Accuracy

To quantitatively assess the alignment accuracy and efficiency of our new algorithm set, and compare with other state-of-the-art mainstream algorithms, we utilized two types of metrics. They are reference-based Q-score and TC-score that require a “true” alignment standard and therefore only applicable to simulated data; and reference-free scaled sum-of-pairs(SP)-score and negative total entropy, which are applicable to all test scenarios (details in STAR Methods, Alignment Accuracy Metrics). Meanwhile to evaluate the comprehensive performance of our algorithm set across key dimensions including sequence number, lengths, and similarity, we constructed four independent benchmark datasets designed with respect to these variations as briefed below (see details in STAR Methods, Construction of Test Datasets):

1. SARS-CoV-2 Number-Tiered Benchmark, to assess the algorithm’s scalability and accuracy for processing real-world sequence sets of different sizes (from 50 to 40,000).
2. Ultramassive SARS-CoV-2 Benchmark, to evaluate performance on a ultra-massive scale against other high-speed algorithms.
3. MPoxBR Length-Tiered Benchmark, to assess the algorithm’s capability in handling 1000 sequences of varying lengths.
4. Human Mitochondrial Based Similarity-Tiered Benchmark, to evaluate precision across different evolutionary distances, with similarities ranging from 99% to 70%.

With the first three benchmarks being derived from real sequences, the similarity-tiered benchmark was generated by us through simulation with^32^.

As shown in Figure 4(a), DAG-align exhibits lowest efficiency when sequence number is below 1000 due to its data structure complexity and Python implementation, but nonetheless far surpass MAFFT and SR-align when sequence set size is larger than 10,000. Halign is the most efficient among these four algorithms. DAG-align has intermediate memory consumption (Figure 4(b)) and achieved highest accuracy in all four methods (Figure 4(c,d)) except for three data points where its performance lags behind MAFFT by a very small margin (Figure 4(c)). In ultra-massive datasets of millions (Figure 5), only DAG-align, SR-align and Halign are tested as other methods (e.g. MAFFT, Muscle) would have taken millions of hours by estimation. DAG-align demonstrates superior accuracy and intermediate memory cost when compared with SR-align and Halign4. The efficiency difference between DAG-align and Halign become smaller for larger datasets with CPU time ratio drop from a few thousands to below 8 (Figure S14). The superior scalability of DAG-align with respect to the number of sequences is mainly due to its better ability in eliminating repetitions caused by shared segments in non-reference sequences as illustrated in Figure S15. In the MPoxBR dataset of 1000 sequences with various lengths, DAG-align exhibits intermediate CPU and memory cost, much lower that naive SR-align but significantly higher than MAFFT and Halign, again caused by its significantly more complex data graph structures that excels for massive datasets. DAG-align demonstrated comparable accuracy with MAFFT in all four metrics, and both are significantly more accurate than SR-align and Halign as indicated by two reference-based metrics (Q-score in Figure 6(c) and TC-score in Figure 6(d)). We added Muscle-v3^33^ and -v5^34^ in the comparison on the simulated human mitochondrial datasets of 100 sequences with various simulated evolutionary distances indicated by similarities in Figure 7. Again, DAG-align exhibit intermediate CPU and memory efficiency, but performed consistently comparable with the most accurate Muscle, which is much more expensive that MAFFT and therefore only tested in this small dataset. The superiority of DAG-align and Muscle over other methods increased significantly for further simulated distances in all four metrics, with the extent being the most for Q-score (Figure 7(c)). Overall, it is apparent that DAG-align strikes the best balance between efficiency and accuracy, and brings MSA of ultra-massive genome sets into the high accuracy era for the first time. The excellence of DAG-align at relatively lower sequence similarity (Figure 7(c,d,e,f)) comes as a pleasing surprise as our initial goal is to mainly accelerate MSA of highly similar sequences through elimination of computation repetition.

**Figure 4:**
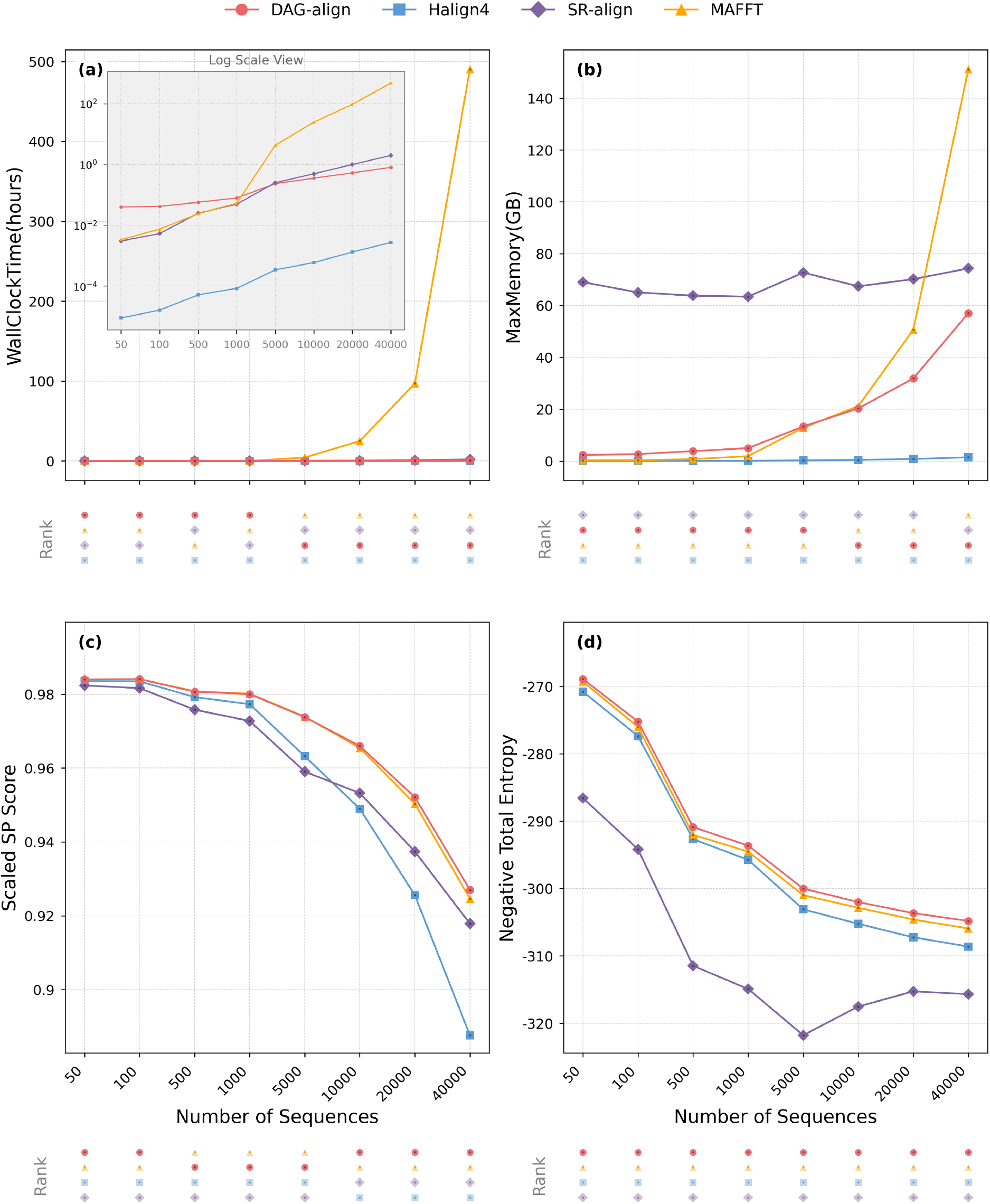
Assessment of accuracy and efficiency of DAG-align, Halign4, SR-align and MAFFT on the SARS-CoV-2 Number-Tiered Benchmark, with genome set sizes varying from 50 to 40,000. Rank of different methods are shown underneath each plot. (a) CPU wall clock time (in hours), a log scale view is provided as inset to better show differences among DAG-align, Halign4 and SR-align. (b) Peak memory usage (in GB). It is important to note that SR-align has a peak memory essentially not related to sequence size. (c) Scaled SP-score. (d) Negative Total Entropy.

**Figure 5:**
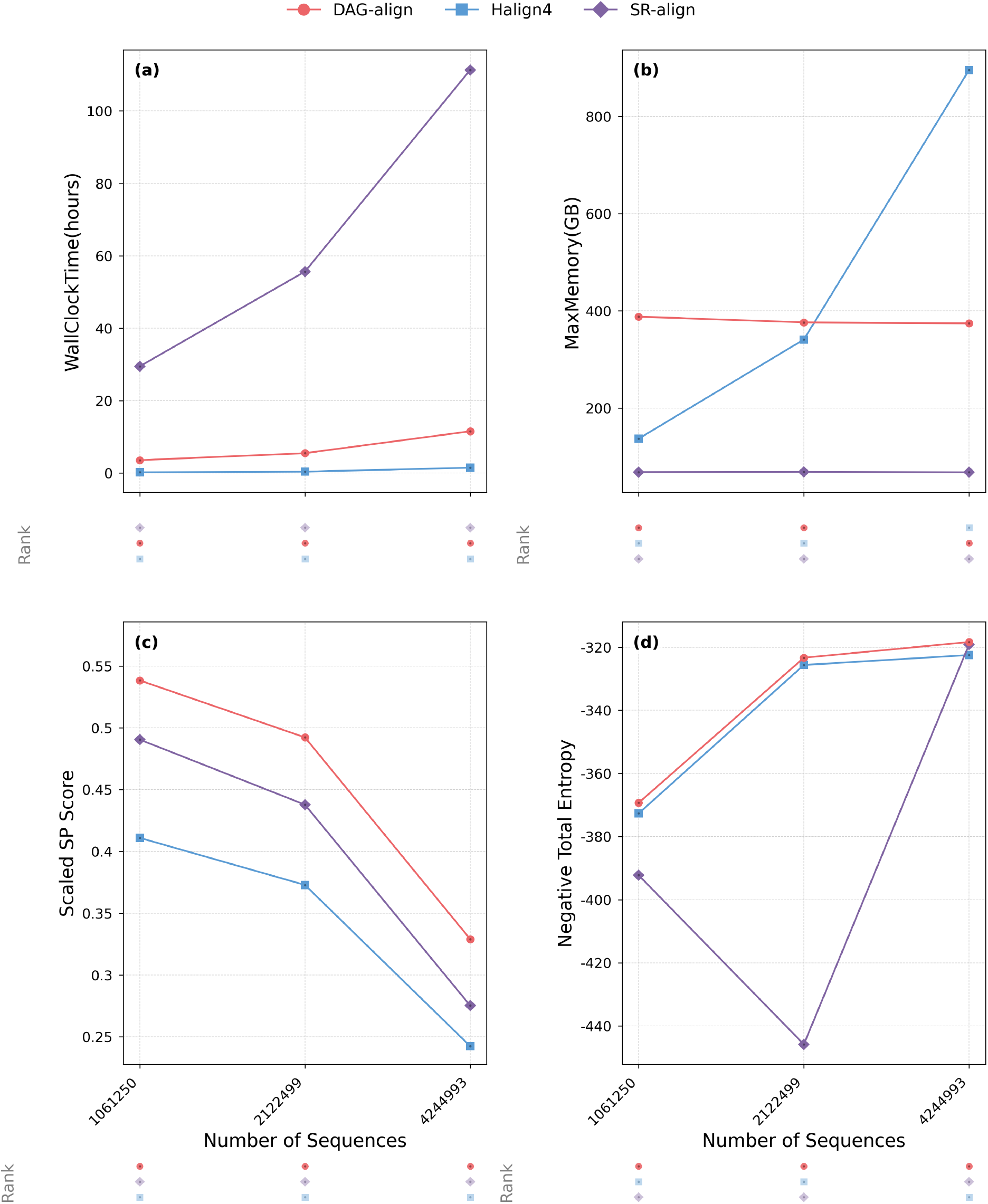
Assessment of accuracy and efficiency of DAG-align, Halign4 and SR-align on the Ultramassive SARS-CoV-2 Number-Tiered Benchmark. Rank of different methods are shown underneath each plot. (a) CPU wall clock time (in hours). (b) Peak memory usage (in GB). (c) Scaled SP-score. (d) Negative Total Entropy. It is noted that there is a dramatic drop of negative total entropy with Halign4 for the 4 million data set. This is a demonstration of the reference sensitivity for Halign4, as the longest sequence in this dataset is somewhat unusual (see more details in STAR Methods, Construction of Test Datasets, Real-Data-Based Datasets, Dataset 2). In this study, SR-align test always have the first sequence of each data set as the reference and no unusual sequences are selected as reference.

**Figure 6:**
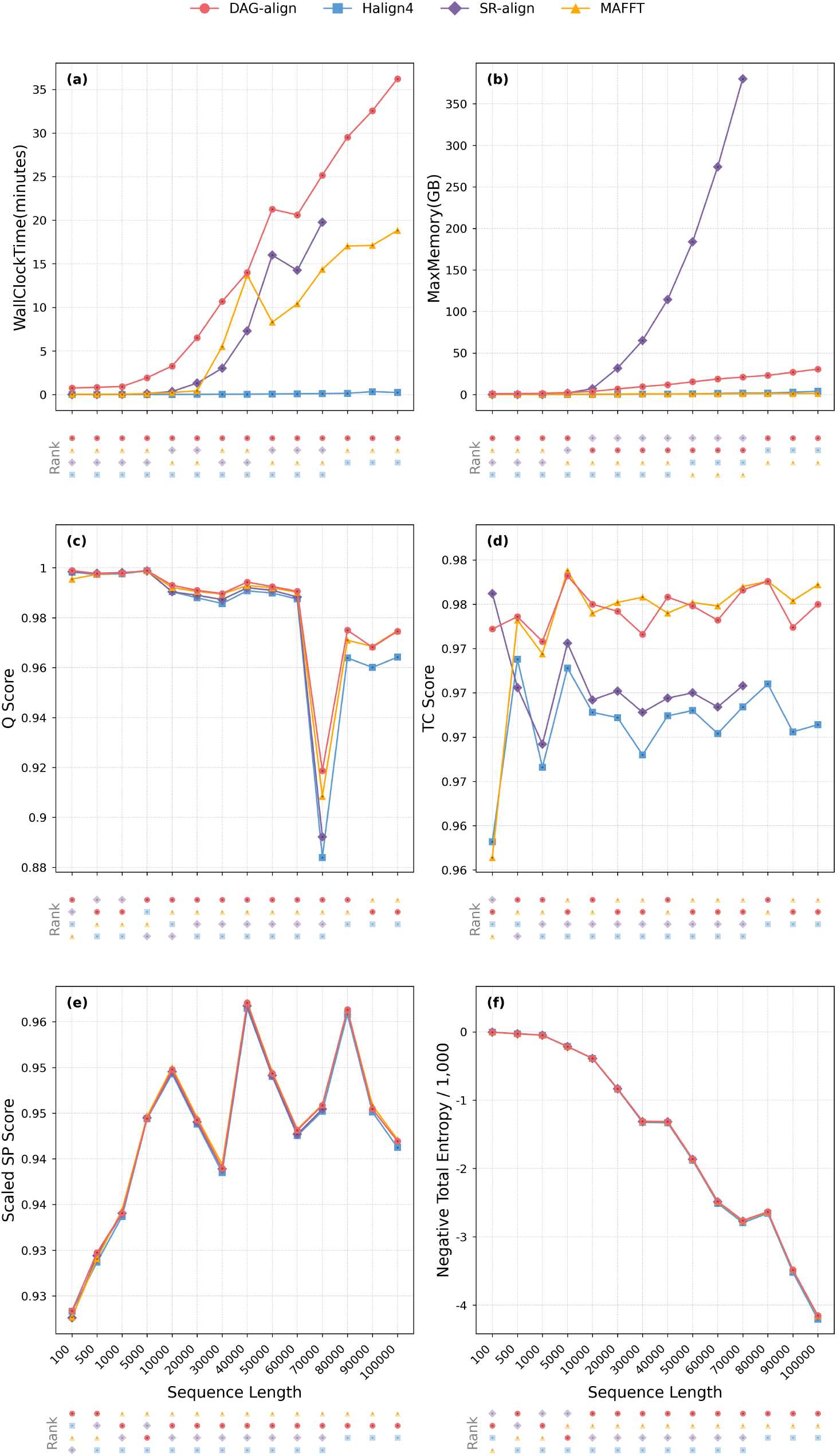
Assessment of accuracy and efficiency of DAG-align, Halign4, SR-align and MAFFT on the MPoxBR Length-Tiered Benchmark. Rank of different methods are shown underneath each plot. (a) CPU wall clock time (in minutes) (b) Peak memory usage (in GB). (c) Q-score. (d) TC-score. (e) Scaled SP-score. (f) Negative Total Entropy. SR-align lacks the last three data points due to the excessive memory demands beyond capacity of our machines.

**Figure 7:**
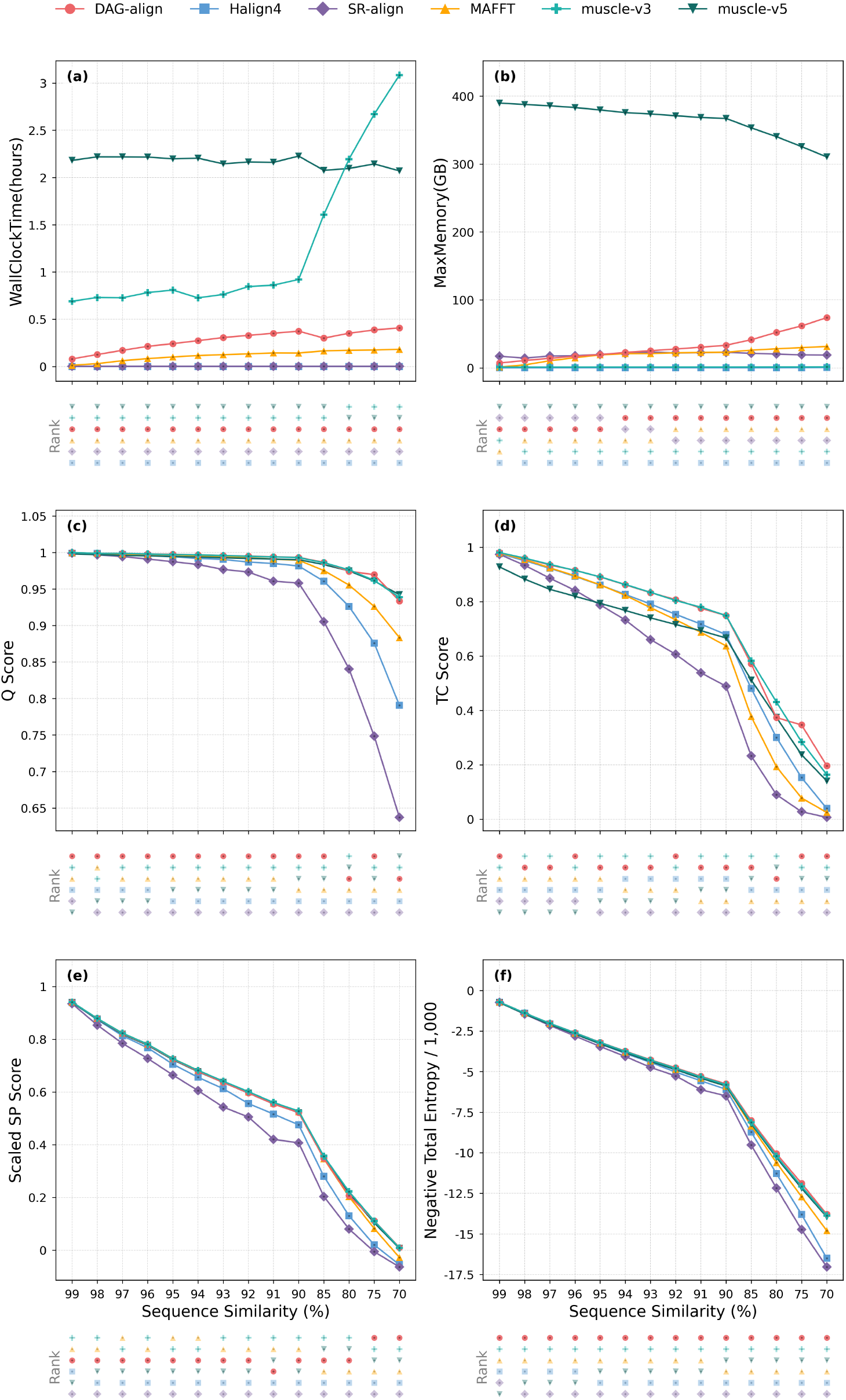
Assessment of accuracy and efficiency of DAG-align, Halign4, SR-align, MAFFT, Muscle v3 and Muscle v5 on the Generated Human Mitochondrial Similarity-Tiered Benchmark. Rank of different methods are shown underneath each plot. (a) CPU wall clock time (in hours). (b) Peak memory usage (in GB). (c) Q-score. (d) TC-score. (e) Scaled SP-score. (f) Negative Total Entropy.

## DISCUSSIONS

The sequence fragment length *L*_*frag*_ is the only adjustable parameter in FTO-DAG construction, with its upper and lower limit being the minimal sequence length *L*_*min*_ in a given sequence set and 1 respectively. If *L*_*frag*_ = *L*_*min*_, there would be negligible TC conflicts and accompanying computation cost for potential cycle elimination. However, such a FTO-DAG is essentially useless as barely any node merging and computation repetition reduction will be achieved. On the other hand, when *L*_*frag*_ = 1, numerous occurrences of single-character “fragments” will lead to a large number of potential cycles, and engender expensive computation for potential cycle elimination. To avoid such dilemma caused by a fixed value of *L*_*frag*_, a stepwise fragment reduction strategy may be utilized by graph resolution adjustment (see Algorithm 7 and Figure S4). Specifically, a sufficiently large *L*_*frag*_ is selected first to reduce potential cycle elimination computation to minimum at the global level, the resulting FTO-DAG may be further compressed locally by stepwise decrease of *L*_*frag*_ so as no significant computation for potential cycle elimination occurs. Reduction of *L*_*frag*_ to 1 is termed atomization of FTO-DAG. Two key factors contributing to the observed dramatic MSA accelerations are the FTO-DAG compression ratio (*η*, see Equation (1) and Figure S13) and average tile widths 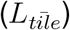 in DAG-TPHMM training, with a larger *η* and a smaller 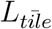 correspond to higher efficiency. Both factors depend critically on the extent of sequence similarity, reduction of which leads to decrease of *η* and increase of 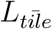.

The simplest approximate MSA is single reference alignment (SR-align), where each non-reference sequence is aligned against a selected reference sequence, with relationship among all non-reference sequences ignored. As illustrated by Figure S2, large amount of computation repetition exists for SR-align. Halign^29,35^ utilizes a number of strategies (suffix trees for Halign3^35^ and BW transform for Halign4^29^) to identify and align non-identical parts, thus greatly reduces such repetition and achieves extreme high efficiency. Partially ordered alignment (POA)^36^ and its variants^36,37^ utilize DAG for MSA, where an sequence sees only preceding (but no subsequent) sequences before commit itself to MSA. While Halign and POA methodologies are seemingly very different, both share critical common points with SR-align. The first is that both (essentially all dynamic programming based) methods utilizes some scoring schemes, which always have limitations. The second is that each sequence fully commit its positions without seeing all other sequences in a given dataset. The third is significant reference dependent for Halign and SR-align or the order of sequence incorporation dependence for POA methods. These are fundamental accuracy limitations and deteriorate with increasing number of sequences. Conventional strategy for overcoming the second/third limitations is to perform pairwise (for each sequence to see all other sequences) or triple (for each sequence to see all pairs) alignments as the basis of MSA assembly. Unfortunately, such strategy is not scalable to large number of sequences. In construction of FTO-DAG, we only merge identical part of sequences through anchors such that no scoring scheme is needed at all. All sequences remain not fully committed until the final decoding stage of DAG-TPHMM, thus excludes premature commitment. The dictionary based anchoring with partial commitment, in contrast to regular dynamic programming alignment, engenders possibility of generating cycles. This is solved by detection and elimination of potential cycles with the aid of TC conflicts. In reality, TC conflicts are rare when sufficiently long fragment length is selected such that the algorithm for eliminating them are rarely called. Training of DAG-TPHMM parameters takes information from all sequences to determine model parameters, which are subsequently utilized to assemble MSA through DAG-Viterbi decoding procedure (detailed in STAR Methods and Figure S9).

We are pleasingly surprised by the superior accuracy of DAG-align in par with Muscle^33^ at relatively lower sequence similarity (Figure 7(c,d,e,f)). This observation indicates more general applicability of DAG-align beyond highly similar sequences such as SARS-CoV-2 genome set. Presently, the expectation-maximization scheme is utilized to train TPHMM model parameters in high dimensional space, there are potential opportunities to further improve accuracy through introduction of additional more effective training strategies, which will be one of the important future directions. It is important to note that MSA itself becoming a fundamentally flawed idea for far divergent sequences. This is due to the fact that by forcing sequences from the further divergent branches of evolution to the same linear coordinate, the more mistakes are needed. Such statement does not undermine the importance of either MSA nor studies of remote homologues. More advanced technologies beyond MSA are needed to accurately describe far divergence of biological sequences.

The major output of our algorithms are FTO-DAGs, DAG-TPHMM models and MSAs for selected datasets. While both FTO-DAGs and DAG-TPHMMs are constructed mainly to achieve accurate MSA, they both have additional important utilities. As ultramassive genome sets being generated for more and more important species, identification of the closest genome for a new sequence becomes more difficult, and comparative genomics questions can not be tackled by naive alignment of single sequences anymore. FTO-DAGs generated by our algorithms may be utilized for accurate and rapid identification of the closest previous genome(s) for continuously arising new ones, and for variant calling or other similar tasks. Probabilistic DAG-TPHMMs models are suitable tools to achieve comprehensive comparison between different species. Accurate MSAs generated for large genome sets in this work may be utilized to perform better evolution analysis (e.g. phylogenetic and recombination investigations), and to carry out genome segments correlation analysis, which may reveal difficult-to-discover molecular interactions and disclose presently unknown encoding segments and regulation pathways. When extended to more complex genomes (e.g. yeast, mouse, human), FTO-DAGs and MSAs together may serve as a reliable framework and a high resolution map for integrating information from other types of sequencing data (e.g. RNA-seq^38^, epigenetic modifications^39^, single cell data, spatially resolved transcriptomics^40^). Last but not the least, with ever increasing quantity and improving quality of sequence data (which are loyal records of evolution experiments in hundreds of millions years), language models^41^ will certainly player more and more important roles in our future investigation of biology. FTO-DAGs may provide information-dense input of genome diversity by riddance of repetition. Accurate MSAs and genome segments correlations derived from which may provide important step stones for training and aligning future language models.

Our FTO-DAGs provide an alternative to current *de facto* standard cyclic pan-genome graphs^42–48^. Besides the demonstrated advantage of facilitating fast MSA, the absence of cycles is also helpful in rapid sifting of virtual paths formed by stretches from distinct genomes. Cyclic pan-genome graphs, on the other hand, provide more agility for handing complex genome structures such as inversions and translocations, addressing of which is not as elegant in a FTO-DAG. Given their respective advantages and limitations, we may select either or both types of graph representation in our analysis based on specific goals and data sets. It is also important to note that our construction strategy of FTO-DAG is to prevent the formation of rather than unroll cycles in a cyclic graph representation, efficient and reliable realization of which remains to be solved. Conversion of a FTO-DAG to a cyclic representation, on the other hand, is a much less challenging task.

In summary, we developed a set of new parallel algorithms, including that for construction of FTO-DAGs, for training of DAG-TPHMMs and subsequent decoding on FTO-DAG with DAG-Viterbi. These algorithmic and implementational efforts jointly realized acceleration of accurate MSA for more than 4 million SARS-CoV-2 genomes by an estimated five orders of magnitude, thus transforming similar tasks from intractable to routine. The linear scaling property and high efficiency of these algorithms reduce the cost of accurate MSA to a negligible fraction of that for sequencing, and therefore are applicable to *arbitrarily large* genome sets expected in future. The FTO-DAGs, DAG-TPHMM models and MSAs generated for many data sets in this study may be utilized by the scientific community for further high resolution analysis of evolution, characterization of genome diversity, and discovering of molecular interactions. Application of these algorithms on other viral genome sets is straight froward. We are actively working on extension of these algorithms to more complex genomes (e.g. bacteria, human) and more species. Completion of the present and planned future work are expected to lay an important cornerstone and open up new frontiers of research for the upcoming era of ultra-massive genome sets.

## RESOURCE AVAILABILITY

### Lead contact

Further information and requests for resources should be directed to and will be fulfilled by the Lead Contact, Pu Tian.

### Materials availability

N/A.

### Data and code availability

- The source code, datasets constructed and necessary tools for manipulating datasets in this study is available at https://github.com/putian74/DAG-align and at https://github.com/putian74/DAG-align/releases/tag/Datasets_and_tools
- Any additional information required to analyze the data reported in this paper is available from the lead contact upon request.

## ACKNOWLEDGMENTS

We gratefully acknowledge all data contributors, i.e., the Authors and their Originating laboratories responsible for obtaining the specimens, and their Submitting laboratories for generating the genetic sequence and metadata and sharing via the GISAID Initiative, on which this research is based. We thank professors Hao Ge at Peiking University, Yang Ji at Zhejiang University and Hao Wu at Wenzhou Institute University of Chinese Academy of Sciences for enlightening discussions in preparation of the manuscript.

## AUTHOR CONTRIBUTIONS

X.L. prepared SARS-CoV-2 data sets, developed and implemented the DAG construction algorithms. X.L. and H.L. jointly performed development and implementation of graph data structure, profile HMM training and Viterbi alignment on graphs. X.L and H.L participated in the manuscript preparation. P.T. conceived the study, designed the algorithm framework (FTO guided construction of DAG, followed by TPHMM training and alignment on such graphs), and wrote the paper.

## SUPPLEMENTAL INFORMATION INDEX

Supplemental Text S1. Redundancy in Existing MSA Algorithms on Similar Sequences.

Supplemental Text S2. Compressed Sparse Alignment Format.

Supplemental Text S3. Single-Reference Alignment (SR-align).

Supplemental Text S4. Parameter Handling in PHMM Training (M-Step).

Supplemental Text S5. Experimental Environment and Parameter Settings

Table S1. Fixed Values for Parameter Updates. These parameters ignore E-step expectations and are reset to these log-space values in each M-step.

Table S2. Log-Space Pseudocounts Added to E-Step Expectations. These values are added to the log-space E-step expectations before normalization.

Figure S1. Repetition Riddance by Dynamic Programming (DP). Schematic illustration on how dynamic programming saves repetition from brute force comparing of all possible sequence arrangement in alignment of two sequences described in the Supplemental text S1.

Figure S2. Repetition Block in Alignment of Similar Sequences. Schematic illustration of blocks of computation repetition engendered by identical segments in pairwise based SR-align and conventional PHMM alignment described in the Supplemental text S1.

Figure S3. Schematic illustration of SR-align algorithm, the baseline of MSA approximation.

Figure S4. Schematic illustration of the Graph Resolution Adjustment Process.

Figure S5. The Canonical State Architecture of a Profile Hidden Markov Model (PHMM).

Figure S6. Multi-stage Parallelization Strategies for Ultramassive Sequence Alignment.

Figure S7. Illustration of the Tiling Calculation for Reducing Computational Complexity.

Figure S8. Schematic Illustration of Graph-guided Initialization of TPHMM Parameters.

Figure S9. Schematic Illustration of Decoding and Path Reconstruction with DAG-Viterbi.

Figure S10. Scaling of Parallelized Graph Construction.

Figure S11. Scaling of Parallelized Viterbi Decoding.

Figure S12. Schematic for the Construction of the Number-Tiered Benchmark Dataset.

Figure S13. Compression ratio for fragment lengths *L*_*frag*_ = 16 and 32 on various sized SARS-CoV-2 genome sets.

Figure S14. CPU wall clock time ratio between DAG-align and Halign4 on various sized SARS-CoV-2 genome sets.

## STAR METHODS

### Construction of Test Datasets

To systematically evaluate algorithm performance, we constructed four distinct benchmark datasets. These datasets are designed to test performance across different dimensions: sequence number, length, and similarity. Three sets are constructed directly from real-world biological sequences and one set generated through simulation.

### Real-Data-Based Test Sets

- Dataset 1: SARS-CoV-2 Genome Number-Tiered Test Set: To test scalability with respect to sequence number, we first constructed a high-quality superset from the GISAID database^1^ (data as of May 15, 2023). We selected sequences marked with ‘high coverage’ and ‘Is complete’ metadata tags, and removed sequences where the total number of degenerate bases exceeded 50% of the sequence length or contained 30 or more consecutive degenerate bases. This process yielded a high-quality superset of 4,245,000 sequences, which was then sorted chronologically and partitioned into 849 basic sets of 5,000 sequences each. A multi-tiered sampling strategy was then applied (Figure S12). We began by selecting 50 of the basic sets (sampling interval of 17, selecting the 1st, 18th, 35th, …, 834th basic sets for a total of 50 sets) and drew varying numbers of sequences (1, 2, 10, 20, 100) from each to create five initial tiers, ranging from 50 to 5,000 sequences. Each of these tiers contains 32 independent replicate datasets. Finally, these replicates were progressively merged pairwise to build three larger-scale tiers of 16 10, 000-genome, 8 20, 000-genome, and 4 40, 000-genome sequence sets.
- Dataset 2: Ultramassive SARS-CoV-2 Number-Tiered Test Set: To specifically evaluate performance on a scale that challenges even high-speed aligners, we constructed an ultra-massive benchmark. Starting from the high-quality superset of 4,245,000 sequences described under **Dataset 1**, we applied an additional length filter, removing any sequences shorter than 28,000 bp or longer than 32,000 bp, resulting in a clean dataset of 4,244,993 sequences. This set formed the largest tier. The motivation for removing these 7 very long/short sequences is that they were found to be very different from all other sequences. If these rare cases were happened to be selected as reference, the result would have been very unfair to Halign4 and SR-align. A medium tier was created by taking approximately half of these sequences (2,122,499), and a minimal tier was created by taking approximately half of the medium tier (1,061,250). As computational cost for all other algorithms are prohibitive, this benchmark was used to compare the performance of DAG-align against SR-align and Halign4.
- Dataset 3: MPoxBR Length-Tiered Test Set: To evaluate performance across various sequence lengths, we utilized a benchmark dataset originally used in the HAlign4 study^29^. This dataset comprises 1,000 monkeypox virus genomes obtained from the MPoxVR database^49^. To create the length-tiered benchmark, prefixes of these genomes were systematically extracted at lengths ranging from 100 to 100, 000 bp, providing a standardized set for assessing performance on sequences of varying length scales.

### Simulated Test Sets

Dataset 4: Human Mitochondria Similarity-Tiered Test Set: To create a controlled benchmark for assessing alignment accuracy across a wide range of evolutionary divergence, we generated a similarity-tiered dataset through a reproducible, script-automated simulation pipeline. The process began by establishing a biologically realistic evolutionary context. We first constructed a high-quality reference alignment from 1,000 complete human mitochondrial genomes using the L-INS-i option in MAFFT (v7.490)^50^. This reference alignment then served as the basis for inferring the best-fit nucleotide substitution model via the ModelFinder functionality in IQ-TREE 2 (v2.2.0)^51^, with the optimal model selected according to the Bayesian Information Criterion (BIC)^52^. With this evolutionary model as a foundation, we simulated the benchmark datasets using AliSim^32^. For each similarity tier, a star-like guide tree was constructed where all 100 sequences evolved from a common ancestor with a uniform branch length. This branch length was calculated to approximate the desired pairwise similarity using the Jukes-Cantor (JC69) distance correction formula^53^. This guide tree, along with the inferred substitution model, served as the input for the AliSim simulator to generate the final datasets. The benchmark comprises 14 distinct tiers of sequence similarity, including levels of 70%, 75%, 80%, 85%, and fine-grained 1% increments from 90% to 99%. Each simulated dataset contains 100 sequences of a fixed length corresponding to the human mitochondrial rCRS genome (∼ 16.5 kb). To enhance realism, the simulation incorporated insertion and deletion events at rates of 0.01 and 0.1 relative to substitution, respectively. The final output is a comprehensive benchmark suite. For each of the 14 similarity tiers, we generated 10 independent replicate datasets. Each replicate consists of two critical components for evaluation: (1) a FASTA file of the unaligned sequences, serving as the input for MSA tools, and (2) the corresponding true alignment generated by the simulator, which acts as the gold standard for accuracy assessment.

### Alignment Accuracy Metrics

#### Q-score and TC-score

The Q-score and Total Column (TC) score are reference-based metrics for evaluating MSA accuracy, particularly prominent in the development of the Muscle algorithm^33^. The Q-score measures the fraction of correctly aligned residue pairs in a test alignment relative to a reference alignment. It is calculated as the number of correctly aligned pairs in the test alignment divided by the total number of aligned pairs in the reference alignment. The TC-score is a stricter metric that measures the fraction of columns in the test alignment that are identical to the corresponding columns in the reference alignment. A column is considered correctly aligned only if all residues in a column are identical to that of the reference. Both scores range from 0 to 1, with higher values indicating better alignment quality.

#### Scaled Sum-of-Pairs (SP) Score

The Sum-of-Pairs (SP) score is a reference-free metric used for evaluating alignment quality^54^. It quantifies the overall quality of a single alignment by summing the substitution matrix scores for all pairs of aligned residues across all columns. The raw SP score for an alignment *A* with *N* sequences and *Z* columns is given by:

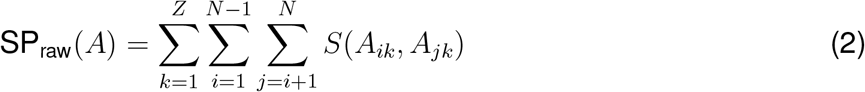

where *A*_*ik*_ is the character for sequence *i* in column *k*, and *S*(*a, b*) is the score from the substitution matrix used in this study. Specifically, we employed a unified scoring matrix where a match between identical bases scores +1, a mismatch between different bases scores -1, an alignment between any base and a gap scores -2, and an alignment between two gaps scores 0. For scoring purposes, degenerate bases were decomposed into their constituent standard bases, and their counts were distributed proportionally. To ensure comparability across alignments of varying lengths and numbers of sequences, we use a scaled SP score. This score represents the average score per aligned non-gap residue pair and is calculated by dividing the total raw SP score by the total number of non-gap residue pairs across all columns:

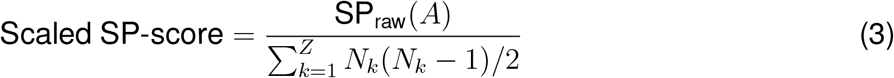

where *N*_*k*_ is the number of non-gap characters in column *k*. This normalization provides a score that reflects the intrinsic quality of the alignment, independent of its size.

#### Negative Total Entropy

Entropy is a reference-free metric that measures the extent of conservation of each column in an alignment. Lower entropy indicates higher conservation and is generally associated with a higher quality alignment^4^. The Shannon entropy for a single column *j* is calculated as:

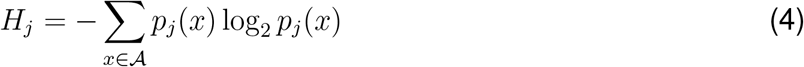

where 𝒜 is the alphabet of characters (e.g., {A, C, G, T, -}) and *p*_*j*_(*x*) is the probability of character *x* in column *j*. The total entropy of the alignment is the sum of the entropies of all columns. To be consistent with other metrics where a higher score corresponds to a better alignment, we use the negative total entropy as our evaluation metric.

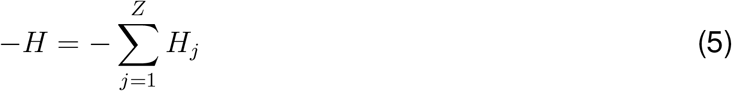

### FTO-DAG Construction Algorithm Set

#### Definitions

- Part 1: Basic Components and Sequence Representation in the Graph
  – Sequence Fragment: A contiguous segment of bases/residues generated by moving a sliding window of length *L*_*frag*_ along an original sequence with a step size of 1. A sequence of length *L* will produce *L* − *L*_*frag*_ + 1 fragments.
  – Node: The basic unit of the graph, with each node encapsulating a sequence fragment and node weight, which is the length of the source genome information list that will be detailed below.
  – Edge: Representing the sequential adjacency of nodes. It is represented as an ordered node pair (*u, v*), where the fragment represented by node *u* immediately precedes the fragment represented by node *v* in the original sequence.
  – Sequence Path: The graph representation of a single biological sequence, which is a simple linear path composed of nodes representing the sequence’s contiguous fragments and the edges connecting them.
  – Start/End/Internal Nodes: In a sequence path, a node with an in-degree of 0 is a start node, a node with an out-degree of 0 is an end node, and all other nodes are internal nodes.
  – Topological Order (TO): The precedence of nodes in a DAG (see Figure 1(c,d)). The normal TO is the inherent 5’ to 3’ arrangement of nodes in a sequence path. This order is a core property of all DAG operations, ensuring that for any directed edge (*u, v*), *u* appears before *v* in the ordering. The reverse TO that goes from 3’ to 5’ is also utilized.
- Part 2: Core Data Structures and Key Properties
  – FTO-DAG (Fragment Topological Order guided DAG): The core data structure of this study. It is a weighted DAG constructed by merging one or more sequence paths via node fusion (see Figure 1(f)). Its most critical design principle is the strict preservation of the TO for all paths in the graph during the construction process. It is important to note that besides original sequence paths corresponding to input sequences, there are also many virtual paths in the constructed FTO-DAG.
  – TO-guided Breadth-First Search (TO-BFS): The standard operation for traversing the FTO-DAG (Algorithm 5). In a forward traversal, a node can only be visited after all its predecessors have been visited. Conversely, in a backward traversal (conceptually reversing the edges), a node can only be visited after all its successors have been visited.
  – Topological Coordinate (TC): A scalar value assigned to each node to efficiently represent its rank in the topological sort (see Figure 1 (c,d) and Algorithm 9). For a forward traversal, the coordinate of a node *v* is one greater than the maximum coordinate of all its predecessor nodes *p*, i.e., *TC*(*v*) = max({*TC*(*p*) | *p* ∈ pred(*v*)}) + 1. Start nodes (with no predecessors) have a coordinate of 1. A reverse TC is defined for backward traversals, where the coordinate of a node *v* is one greater than the maximum coordinate of all its successor nodes *s*. End nodes (with no successors) have a reverse coordinate of 1.
  – Source Genome Information List: A core property of a node, it stores information about all genomes that share the node’s sequence fragment (e.g., genome ID and fragment position) with consistent TO. For example, if a genome have multiple segments sharing the same fragment as a node in a FTO-DAG, it will appear at most once in this list, or does not appear at all if all the segments do not have correct TO on that node.
  – Node Weight: A numerical attribute of a node, its value is defined as the cardinality of its Source Genome Information List, intuitively reflecting the number of original sequences sharing the sequence fragment in this node with correct TO.
  – Edge Weight: A numerical attribute of an edge, its value quantifies the frequency of TO-consistent and physically contiguous co-occurrence in all original input sequences for the two fragments represented by the connected nodes. Formally, the weight of an edge (*u, v*) is the number of sequences that satisfy the condition of having ‘(sequence ID, position p)’ in the source genome information list of *u* and ‘(sequence ID, position p+1)’ in the source genome information list of *v*.
- Part 3: Graph Construction and Merging Mechanisms
  – Input Sequence Path: The most basic building block in the incremental construction process, converted from a single original linear sequence (see Figure 1 (e), Algorithm 1).
  – Input FTO-DAG: In a graph-to-graph merging process, this is a structurally complete FTO-DAG that serves as the source graph.
  – Target FTO-DAG : In a graph construction or merging operation, this is the persistent graph that acts as the destination graph.
  – Anchor: A special class of nodes identified in the target FTO-DAG when integrating an input unit (see Algorithm 2, details defined in main text The DAG-align algorithm set, The FTO-DAG graph construction algorithm set, Phase 1 Anchor Search).
  – Node Merging: A graph operation that transfers information (primarily the Source Genome Information List) from a matching input node to a target anchor, followed by the deletion of the source node and the redirection of all its incoming and outgoing edges to the anchor (Algorithm 4).
  – TC Conflict: the violation of monotonic increase for TC on a FTO-DAG path (see Figure 1(c,d) and caption for illustration and definition.)
- Part 4: Advanced Concepts and Application Components
  – Compression Ratio *η*: A global metric that measures the degree of compression of the FTO-DAG relative to the original sequence data (see Equation (1) in the main text).
  – Secondary Merging: A process of streamlining the graph structure by merging pairs of nodes that share both the same sequence fragment and identical TC, thereby improving the compression ratio (Algorithm 8). Note this process is done for both normal forward TO and reverse TO iteratively until no mergeable node pairs is found. The motivation of this operation is that the strict anchor criteria leave some of mergeable node pairs in separate paths. It is important to note that aggressive merging by loosing anchor criteria in the earlier phase of FTO-DAG construction may risking wrong commitment and therefore decrease final alignment quality.
  – Graph resolution adjustment: A procedure wherein all nodes in the graph are modified to retain only the last 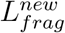 bases of their original sequence fragments. Nodes are then supplemented upstream of start nodes to compensate for information lost due to truncation (see Figure S4, Algorithm 7). When 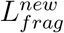 is 1, this process is termed Atomization.
  – Main path: The path with the maximum cumulative weight, as identified by a maximum weight path algorithm on the FTO-DAG (Algorithm 6). When merging graphs, anchor search for the input FTO-DAG is performed exclusively for main path nodes to improve efficiency.
  – Maximal Unbranched Path: A path stretch in a FTO-DAG where all internal nodes have exactly one parent and one child, while its terminal nodes are either start/end nodes or nodes having multiple parent/child nodes. Such paths are work units for downstream parallel training and decoding.
  – Alignment Reference Path: The main path of the atomized, pruned and coarsened FTO-DAG. This path represents the core conserved features of the sequence set. It is mapped back to the original graph or other graphs (DAG-TPHMM Multiple Sequence Alignment Algorithm: Workflow and Details) and used for defining the state positions and initializing parameters of the subsequent DAG-TPHMM model. (see Figure S7, Figure S8 and Figure S9)
  – DAG-TPHMM Node Tile Width: A node attribute introduced to optimize the time and space efficiency of downstream DAG-TPHMM training; it serves as an application-level performance optimization parameter (see Figure S7, Algorithm 21). During DAG-TPHMM forward/backward computation, only the values for state positions within the node’s tile width are calculated.

#### FTO-DAG construction process details

There are two stages for FTO-DAG construction as briefed in the main text and illustrated in Figure 1(f,g).

- Incremental Construction:
  1. Path Conversion: Convert an input single linear sequence into an Input Sequence Path using a fixed fragment length (default *L*_*frag*_ = 32) (see Figure 1(e), Algorithm 1).
  2. Anchor search: For each node in the input sequence path, search for a qualifying anchor in the target FTO-DAG. The search may return an anchor or none (see Algorithm 2).
  3. TC Conflict Detection and Elimination: Check (see Algorithm 3) for TC Conflicts in initial matching input nodes. If detected, Algorithm 4 is applied to return a TC conflict free final matching input nodes. Otherwise, the initial matching input nodes are final matching input nodes.
  4. Node Merging: Merge each final matching input node into the corresponding anchor. Subsequently, integrate the remaining input FTO-DAG nodes into the target FTO-DAG based on their connection to the final matching input nodes (now merged into the corresponding anchor) (see Algorithm 5).
- Graph Merging:
  1. Main Path Determination: Calculate the main path in the input FTO-DAG (Algorithm 6).
  2. Anchor search: Search anchor in the target FTO-DAG for main path nodes of the input FTO-DAG (Algorithm 2), return initial matching input nodes and corresponding anchor.
  3. TC Conflict Detection and Elimination: Same as in the incremental construction stage.
  4. Graph-to-Graph Merging: The final matching input nodes (on the main path) are merged first into corresponding anchors in the target FTO-DAG. Subsequently, integrate the remaining input FTO-DAG nodes into the target FTO-DAG based on their connection to the final matching input nodes (now merged into the corresponding anchor) (Algorithm 5).

#### Post-Construction Graph Structure Optimization

The initial fragment length for building a FTO-DAG maybe adjusted in preliminary test runs with randomly sampled sequences, the goal is to have a good balance between achieving high compression rate and low TC conflict occurrence. After the initial FTO-DAG construction is accomplished with a selected default fragment length (*L*_*frag*_ = 32 in this study), it may be further optimized with the following two operations.

1. Graph Resolution Adjustment: Once all sequences are integrated, perform a resolution adjustment operation to refine the global fragment length from the initial length to a smaller value (e.g. 16 in this study) to enhance the graph’s resolution (see Figure S4, Algorithm 7).
2. Secondary Merging: Subsequently, execute a global secondary merging process. The algorithm traverses all nodes in the graph and merges any pairs of nodes that share the same fragment content and TC. This merging step is a critical finishing touch for constructing a high-resolution, high-compression FTO-DAG, as it aims to eliminate structural redundancies that arise from both the resolution adjustment (Algorithm 8) and missed mergeable node pairs in the initial construction stage due to the strict criteria of anchor search.

### DAG-TPHMM and DAG-Viterbi Algorithm Sets

#### Graph-guided DAG-TPHMM Parameter Initialization

To ensure that model training begins from a proper starting point, we have designed a graph-guided parameter initialization strategy. The core of this strategy is to leverage the topological structure and statistical information of the constructed FTO-DAG to establish a biologically meaningful initial parameter set for a DAG-TPHMM (Figure S8). The specific steps are listed below:

- Extraction of the Alignment Reference Path. The core of the initialization is the extraction of the reference path that represents the central, shared features of the sequence set. To overcome path length induced biases in direct searches, we devised a two-stage preprocessing strategy:
  – Phase 1: Graph Pruning: This stage aims to filter out structural noise. We streamline the graph by removing any edge whose weight is less than a threshold (e.g. 1% in this study) of the weight of its source or target node, thereby preserving the core topological information flow (Algorithm 10).
  – Phase 2: Graph Coarsening: This stage aims to eliminate the length-induced bias. We compress unbranched linear segments in the graph (i.e., contiguous nodes where the in-degree and out-degree are both 1) into super nodes, as in the case of constructing TO guided training task units (see Figure S6(b)). The weight of a super node is defined as the average weight of all atomic nodes it contains (Algorithm 11). Finally, we apply a dynamic programming algorithm to this preprocessed, coarsened graph to efficiently identify the main path in this graph. It noted that extraction of this main path is exactly the same for that in preparation of subgraph merging except no pruning/coarsening preprocessing is performed in the later case. This path is then expanded back into its constituent atomic nodes to reconstruct the final main path, which is termed alignment reference path, in the original atomic graph (Algorithm 6).
- Fine-grained Parameter Initialization Based on Graph Topology. The extracted reference path is used to define the skeleton of the DAG-TPHMM, with each node in the reference path defines a regular state position in the model. Subsequently, through a fine-grained graph analysis process, we set initial values for the model’s emission and transition probabilities that are consistent with the statistics from FTO-DAG. The core of this process lies in analyzing “bypass partial graphs” that surround the reference path. The specific steps are as follows (Algorithm 12):
  1. bypass partial graph Identification: We first remove the reference path from the graph, thereby partitioning the original graph into a series of independent connected components, which are referred to as bypass partial graphs. These are structurally distinct from the subgraphs generated by partitioning the input data during the parallel construction phase, where each of original sequence may be fully traced. In a bypass partial graph, many original sequences only exist in part. The algorithm identifies those bypass partial graphs that connect to the main path at two or more “Attachment Point”. They represent variations or structural variants from the main path.
  2. bypass partial graph Classification and Parameter Initialization: Next, the algorithm classifies each bypass partial graph by comparing the length of its longest path to the length of the main path segment it spans, and initializes different parameters accordingly:
    – Emission Probability Initialization (for Parallel Paths): When the bypass partial graph length is equal to the corresponding reference path segment length, it is considered a “parallel path” (typically representing single nucleotide polymorphism or alleles). At this point, we calculate the TC of each node in the subgraph relative to the start of the main path segment. The emission probability of a base at any state position defined by the reference path is then determined by the weighted base frequencies of the node at that position itself, as well as all nodes from parallel paths that are aligned to it via their TCs.
    – Transition Probability Initialization (for Insertion Events): When the bypass partial graph length is greater than the corresponding backbone segment, it is identified as a potential “insertion hot spot”. The algorithm marks the entire segment of the reference path spanned by this bypass partial graph (from the divergence point up to, but not including, the convergence point) as a high-insertion probability region. In the model, the relevant transition probabilities for this entire segment (e.g., the probability of transitioning from a match state to an insert state) will be correspondingly increased. Through this graph topology-based analysis, we not only accurately infer the base emission propensity for each model state position from the global data, but also identify potential “insertion hotspot”, thereby generating an initial DAG-TPHMM parameter set that is consistent with the intrinsic structure of the data.
- Initialization Rules and Parameter Space Exploration. To increase the likelihood of obtaining a good local minima in the high dimensional parameter space for alignment, we enhance our search by exploring different starting points in the parameter space. Specifically, we generate two independent sets of initial parameters by adding two different pseudo-counts (see below). The parameter set that yields a higher final overall graph likelihood after one full round of Baum-Welch training is then selected for subsequent decoding steps. Some transition probabilities (e.g., *a*_*D,M*_, *a*_*D,D*_) are fixed to specific values based on empirical settings that we found to produce good results through extensive experimentation.
  – Emission Probability Matrix (*E*_*S*_(*x*))
    * Match States (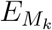): For each regular state position *k* as defined by the reference path, its emission probability is derived from the weighted base frequencies of all nodes sharing the TC with corresponding reference path node. Additionally, for each base a pseudo-count (*λ*_1_ = *e*^−3^ or *λ*_2_ = *e*^−5^) is added to the count with a subsequent renormalization.
    * Insert States (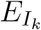): The emission probabilities for all insert states are initialized to a uniform distribution (i.e., a probability of 0.25 for any base).
  – Transition Probability Matrix (*a*_*S,S*′_)
    * Transitions from Start State: *P* (*M*_1_ | Start) = *P* (*I*_0_ | Start) = *P* (*D*_1_ | Start) ≈ 1*/*3
    * Transitions from Match State (*M*_*k*−1_): The default values for *a*_*M,M*_, *a*_*M,I*_, and *a*_*M,D*_ are set to 1 − *e*^−2^ − *e*^−5^, *e*^−5^, and *e*^−2^, respectively.
    * Transitions from Insert State (*I*_*k*−1_): *a*_*I,M*_ = 0.5, *a*_*I,I*_ = 0.5, *a*_*I,D*_ = 0
    * Transitions from Delete State (*D*_*k*−1_): *a*_*D,M*_ = 0.5, *a*_*D,D*_ = 0.5, *a*_*D,I*_ = 0
    * Transitions to End State: 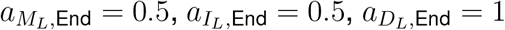
  – Graph-guided Dynamic Adjustment: If the length of a branch in the graph is found to be longer than the main path segment it spans, that region is identified as a high-insertion area. In these regions, the corresponding *a*_*M,I*_ probability is increased by 0.1 from its default value with a subsequent transition matrix renormalization for that state position.

#### DAG-TPHMM Node Tile Computation

In conventional PHMM^4^, each node *v* in a sequence needs to iteratively compute its match probability with all state positions. This need remains for each node in a probabilistic model on a FTO-DAG. However, the vast majority of state positions (corresponding to the reference path nodes in the preprocessed FTO-DAG) contribute negligibly to the final result. Additionally, the topological structure of the FTO-DAG suggests that a graph node *v* will most likely align with high probability to only a local, contiguous region of the state positions as defined by the reference path. To avoid a massive amount of computation with state positions of negligible contribution, we introduce a heuristic tiling strategy. The core objective is to use the graph’s topology to define a high-probability range (the “tile”) of state positions for each graph node *v*. All subsequent calculations will be strictly confined to this tile, thus reducing training computation from being proportional to number of state positions *L*_*DAG*−*T PHMM*_ to the average tile length 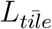. This calculation process relies on a reference path within the graph, whose nodes have a direct correspondence with the regular state positions. There are two different scenarios need to be addressed:

1. For preparation of tiles in the global FTO-DAG for training, calculation is based on the global reference path as expected.
2. For preparation of subgraphs for parallel in DAG-Viterbi decoding, a critical step is to transfer the reference path in the global FTO-DAG to each subgraph. This is achieved by transform the global reference path into a input sequence path and merge it to each subgraph using the standard construction workflow.

With the reference path established for all graphs needing it. The tiling computation (see Figure S7, Algorithm 21) can be performed independently and consistently with the following two steps:

1. Locate Reference Path Interval: For any non-reference path node *v* in the graph, the algorithm first analyzes its shortest paths to the reference path to determine the two nearest reference path nodes, between which the node *v* is “inserted”. For a reference path node, the interval is its own position.
2. Padding: The reference path interval where the node is located is considered the core state position region where its optimal state is most likely to occur. To ensure the robustness of the calculation, we add a padding buffer (default to 100 state positions) to each side of this core region, ultimately forming the computational “tile” for that node.

With tiling, computational resources are effectively utilized on the subset of state positions with the highest match/insert probability, achieving an increase in computational efficiency of about 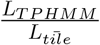 while ensuring minimal impact on the accuracy of the final alignment result. For the largest SARS-CoV-2 genome data set, *L*_*T PHMM*_ and 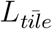 are 29902 and 234 respectively, implies an upper bound acceleration of 128 with a small engendered cost for calculating tiles and bookkeeping.

#### DAG-TPHMM Training

One round of the Expectation-Maximization (EM) algorithm is applied to optimize the model parameters with the aim of maximizing the likelihood of the observed data (the atomized FTO-DAG). The forward and backward calculations are performed by sequentially computing the *α* and *β* vectors for each node via the TO Guided Multi-process Graph Traversal Algorithm (Algorithms 5 and 13).

- Step 1: Expectation Step (Forward-Backward Algorithm)
  – Forward Algorithm (*α* calculation): This algorithm computes the probability 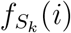 (with *S* representing *M, D* or *I*), which is the probability of the FTO-DAG arriving at node *i* and being in state *S*_*k*_. Cases with multiple parent nodes are handled by probability aggregation (Equation (6)). The recurrence relation is as follows:

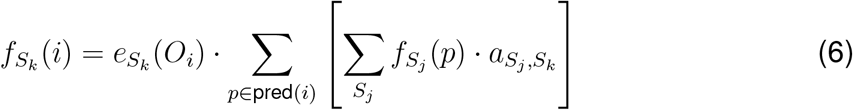

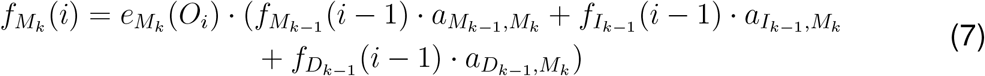

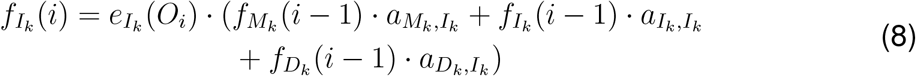

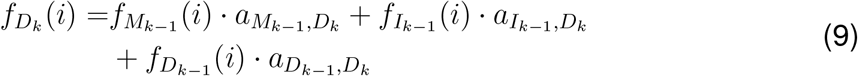
  – Backward Algorithm (*β* calculation): This algorithm computes the probability 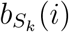, which is the probability of generating the subsequent sequence given that the FTO-DAG is in state *S*_*k*_ at node *i* (with *S* representing *M, D* or *I*). Multiple child nodes are handled by probability aggregation (Equation (10)). The recurrence relation is as follows:

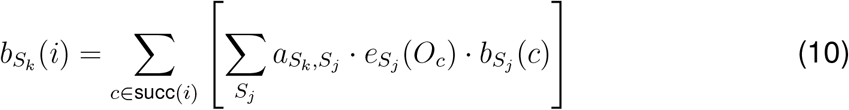

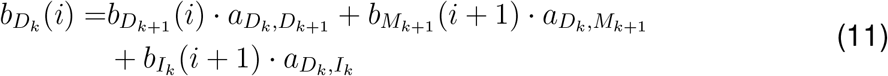

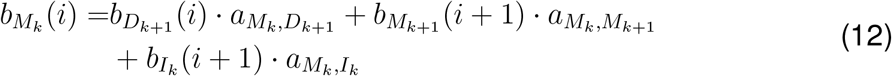

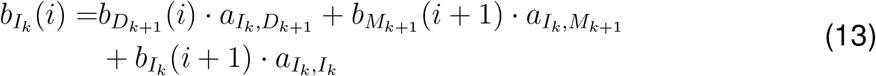
- Step 2: Maximization Step (Parameter Re-estimation) The results from the forward-backward algorithm are used to re-estimate the parameters (Algorithm 16).
  – Transition Probability Update 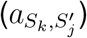: The new probability is calculated as the expected total number of a specific transition type divided by the expected total number of all outgoing transitions from the source state.

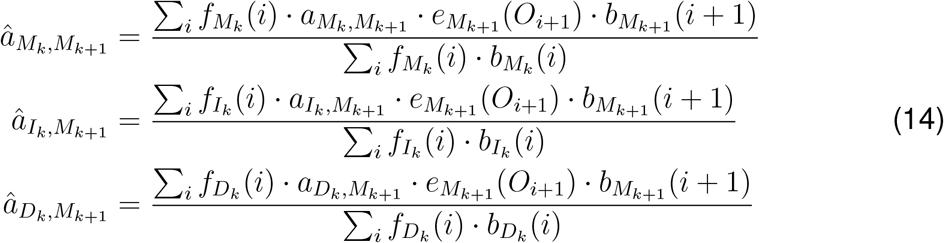

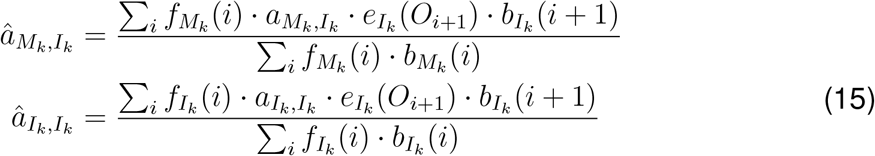

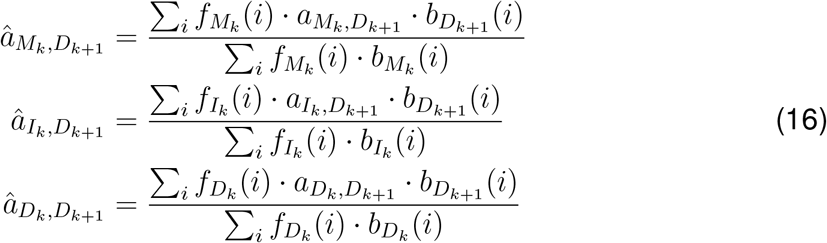
  – Emission Probability Update 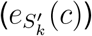: Updated based on the expected number of times a specific character is emitted from a given state.

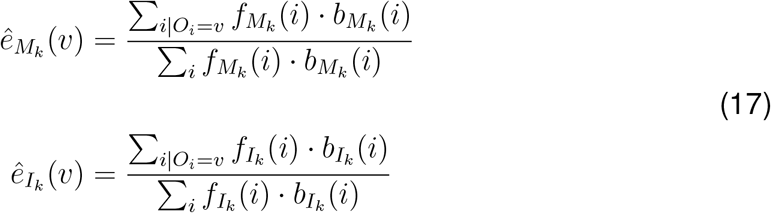
  – Regularization: Pseudo-counts are added to all learned parameters to prevent overfitting. More details are available in SI Text S4.

#### Decoding: The DAG-Viterbi Algorithm

After parameter training, the next core task is to find the most probable sequence of hidden states for the given FTO-DAG using this optimized parameter set. This process is known as decoding, and it aims to identify the Viterbi paths that generates the observed FTO-DAG with the highest probability. To this end, we have designed an efficient DAG-Viterbi algorithm (Algorithm 17) that can be executed in parallel by utilizing the subgraph structure generated during the initial graph construction phase (Figure S6(c)). The rationale for this hybrid approach, with training on the global FTO-DAG and decoding on subgraphs in parallel, stems from the different computational requirements of each phase. The parameter training phase primarily relies on aggregated node and edge weight data from the entire graph and does not require querying the detailed `Source Genome Information List`. In contrast, the Viterbi decoding and traceback phase requires frequent and intensive lookups into this list for each node to attribute states to original sequences. By parallelizing the decoding on smaller subgraphs, these lookups become significantly faster, drastically reducing the computational bottleneck of this step, as demonstrated by our performance benchmarks (Figure S11). There is also a trade-off between repetition and genome information list lookup. With the global FTO-DAG, there is least repetition and excessive lookups, while with single sequences, there will be no lookups but excessive repetition. Using subgraph decoding is a balanced decision. After the DAG-TPHMM parameters have been globally trained on the complete FTO-DAG, each subgraph is dispatched to an independent process. Within each process, the DAG-Viterbi algorithm is executed exclusively on the nodes and edges of that subgraph, using the globally optimized parameters. This operation yields a partial `FinalOptimalStateMaps` data structure for each subgraph. The final, global alignment is then reconstructed by aggregating the results from these partial state maps in the subsequent MSA generation step. The entire decoding process is divided into the following two main stages:

- 1. Viterbi Forward Pass. This state compute likelihood score and fill the one-step tracing pointer matrix for each node. The algorithm employs a topologically-guided parallel traversal (this is another level of parallelization of decoding computation independent from subgraph parallelization) from the start nodes to the terminal nodes of the graph to compute the optimal Viterbi score 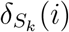 and the tracing pointer matrix *ψ*(*i, S*_*k*_) for each node *i* in each state position *S*_*k*_ (see Algorithm 18), where *S* representing *M, D* or *I*. For any non-start node in the graph, its calculation is clearly divided into two steps: Likelihood Score Aggregation. To handle paths from multiple parent nodes (*p* ∈ pred(*i*)), the algorithm first independently calculates the optimal source score aggregated from all parent nodes for each possible predecessor state *S*^′^, where *S*^′^ representing *M* ^′^, *D*^′^ or *I*^′^, uniformly represented as *δ*_*S*′_ (*i* − 1), and records the optimal predecessor node that provides this score. This score is calculated as follows:

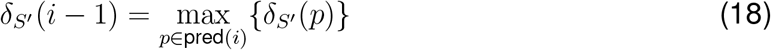

Recursive Update & Path Tracking. Subsequently, the algorithm uses these aggregated scores to calculate the final score for the current node *i* (corresponding to observation *O*_*i*_) in each state via the standard Viterbi recurrence relations, and finds the optimal predecessor state:

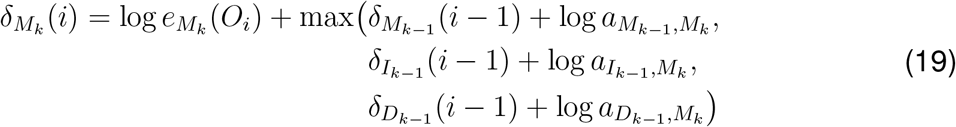

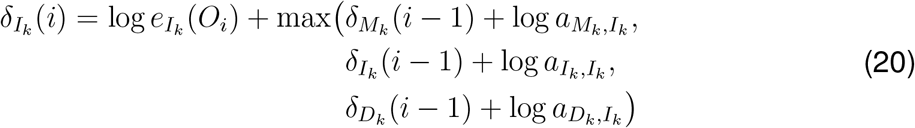

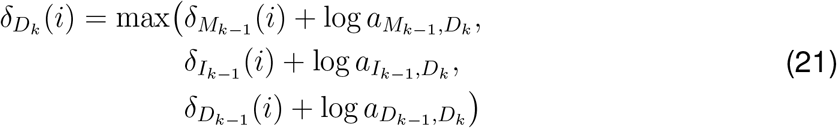

The final tracing pointer matrix *ψ*(*i, S*_*k*_) is jointly determined and stored the optimal predecessor node and the optimal predecessor state based on the results of these two steps.
- 2. Viterbi Backward Pass: Optimal Path Reconstruction and Source Attribution. After the Viterbi forward pass is complete, the Viterbi scores for all nodes have been computed and tracing pointer matrix saved, the algorithm initiates a TO guided parallel backward traversal process to reconstruct the unique optimal path for each original genome (Algorithm 19), ultimately generating the `FinalOptimalStateMaps` data structure, which has the format `node -> optimal state -> set of source information` (Figure S9(a)). More specifically, this data structure is a nested dictionary: its outer keys are the nodes in the graph, and its inner keys are precise state representations *S*_*k*_, where *S* represents the state type (Match, Insertion, Deletion) and *k* is its positional index in the DAG-TPHMM model. The value corresponding to this state key is a set that stores all the genome IDs and corresponding positions that adopt this state at this node. The entire population process of the `FinalOptimalStateMaps` is accomplished through a child-to-parent reverse traversal guided by the graph’s topological order, and is divided into the following two stages:
  – Stage 1: Initialization by backtracking from Terminal Nodes The backtracking process does not start from the beginning of the graph, but rather is initiated from the endpoints of all paths (i.e., nodes with an out-degree of 0). The algorithm initializes by starting from all terminal nodes in the graph, determining their optimal final state *S*_*k*_ based on their highest Viterbi scores, and populating the initial entries of FinalOptimalStateMaps with their entire source information `{(genome_id, position)}`. This step establishes a clear, traceable starting point for the traceback path of each sequence.
  – Stage 2: Path Attribution Based on Physical Adjacency. After initialization, the algorithm begins a child-to-parent reverse traversal. For any parent node *i* currently being processed, the algorithm examines its processed child node *w*. Using the pre-stored tracing pointer matrix *ψ*, it identifies the optimal source state in the parent node *i* that corresponds to the child’s state *S*_*k*_. The crucial step of ensuring physical path continuity is accomplished through a set intersection operation. First, it converts all source information `{(genome_id, pos_w)}` for the child’s state *S*_*k*_ into an “expected” set of predecessor sources `{(genome_id, pos_w - 1)}`. Then, it finds the intersection of this set with all “available” original source information at the parent node *i*. The elements in the intersection are the physically contiguous sources that followed this optimal path, and they will be accurately attributed to the corresponding optimal state of the parent node *i*. After the entire process is complete, `FinalOptimalStateMaps` contains the precise source genome information corresponding to each optimal state of each node in the graph, implicitly defining the final MSA.
- 3. Alignment Reconstruction and MSA Generation. Once the graph traceback is complete, the `FinalOptimalStateMaps` data structure has assigned a unique optimal state to the base of each genome. The final alignment reconstruction step aims to efficiently convert this complex, graph-based path information into a standard MSA matrix (Figure S9(b) and Algorithm 20). This process follows an efficient two-pass graph traversal strategy, avoiding the costly overhead of reconstructing the full path for each sequence:
  1. Layout Planning Traversal: The algorithm first performs a topological traversal of the graph to directly compute and plan the overall layout of the MSA. In this pass, it calculates the maximum number of insertion columns required between each pair of neighboring state positions (defined by state positions *k* and *k* + 1) and simultaneously records the specific insertion length for each sequence at each position *k*.
  2. Matrix Population Traversal: After constructing a pre-filled MSA matrix with gap characters (‘-’) based on the above statistics, the algorithm performs another topological traversal. In this pass, it places the base corresponding to each node and state precisely at its predetermined coordinate in the matrix (Figure S9(b)).
    – Bases emitted by a match state *M*_*k*_ are placed in the corresponding state position *k* for that sequence.
    – Bases emitted by an insert state *I*_*k*_ are placed, right-aligned, in the block of columns dynamically inserted for that position, according to the insertion length of the sequence itself.

When all bases are filled into the matrix according to their match or insert states, any remaining pre-filled units (‘-’) naturally form the alignment gaps for that sequence, which correspond to the delete states *D*_*k*_ in the path. Through this precise and efficient process, all sequences are integrated and arranged into a dynamically sized matrix, ultimately generating a high-precision MSA.

### Pseudocode Algorithms

#### Algorithm 1 Sequence to Path Conversion

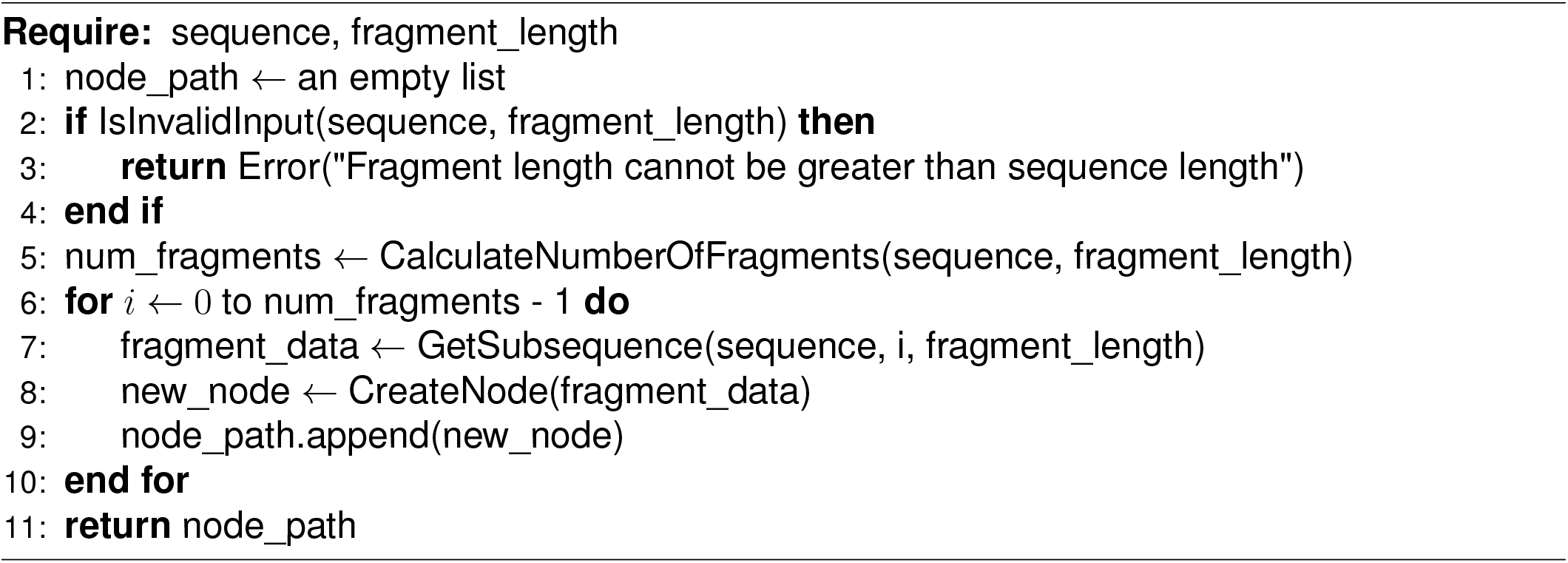

#### Algorithm 2 Anchor Finding

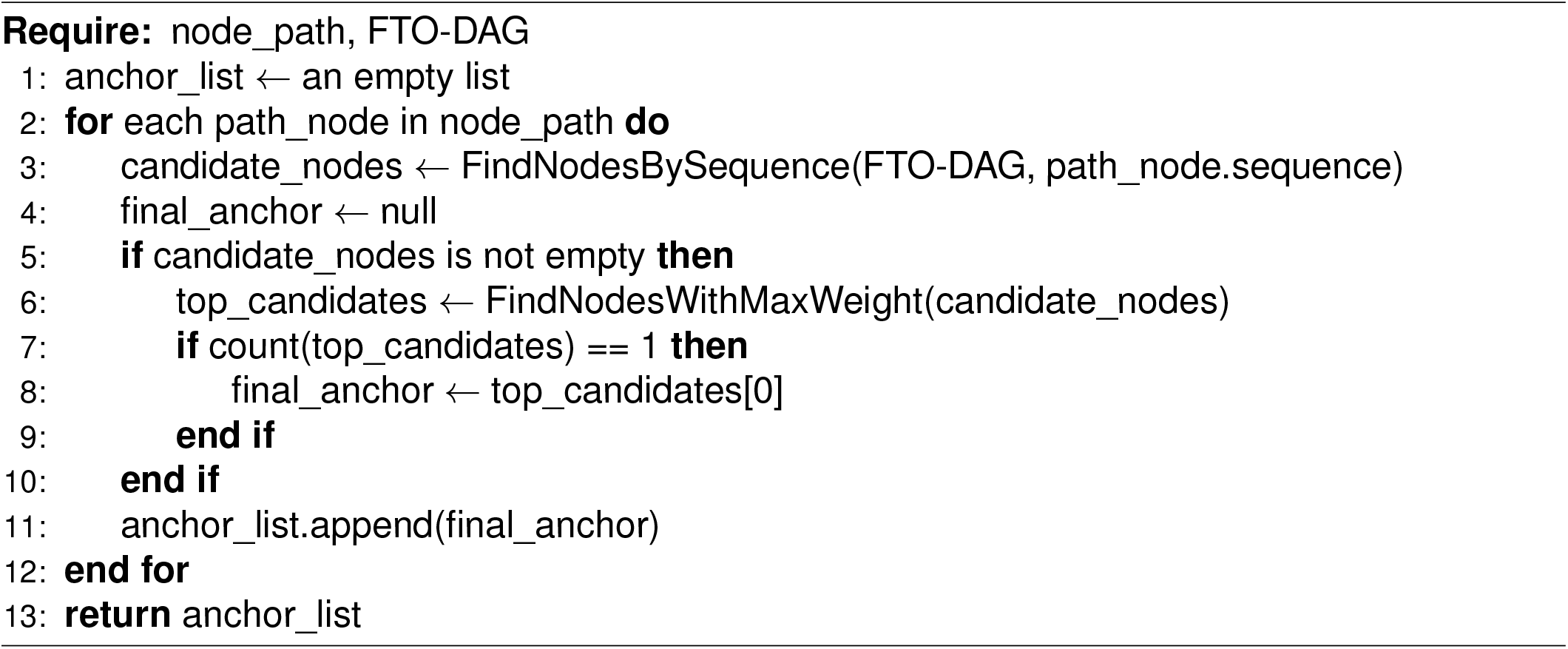

#### Algorithm 3 Find Best Anchor Chain

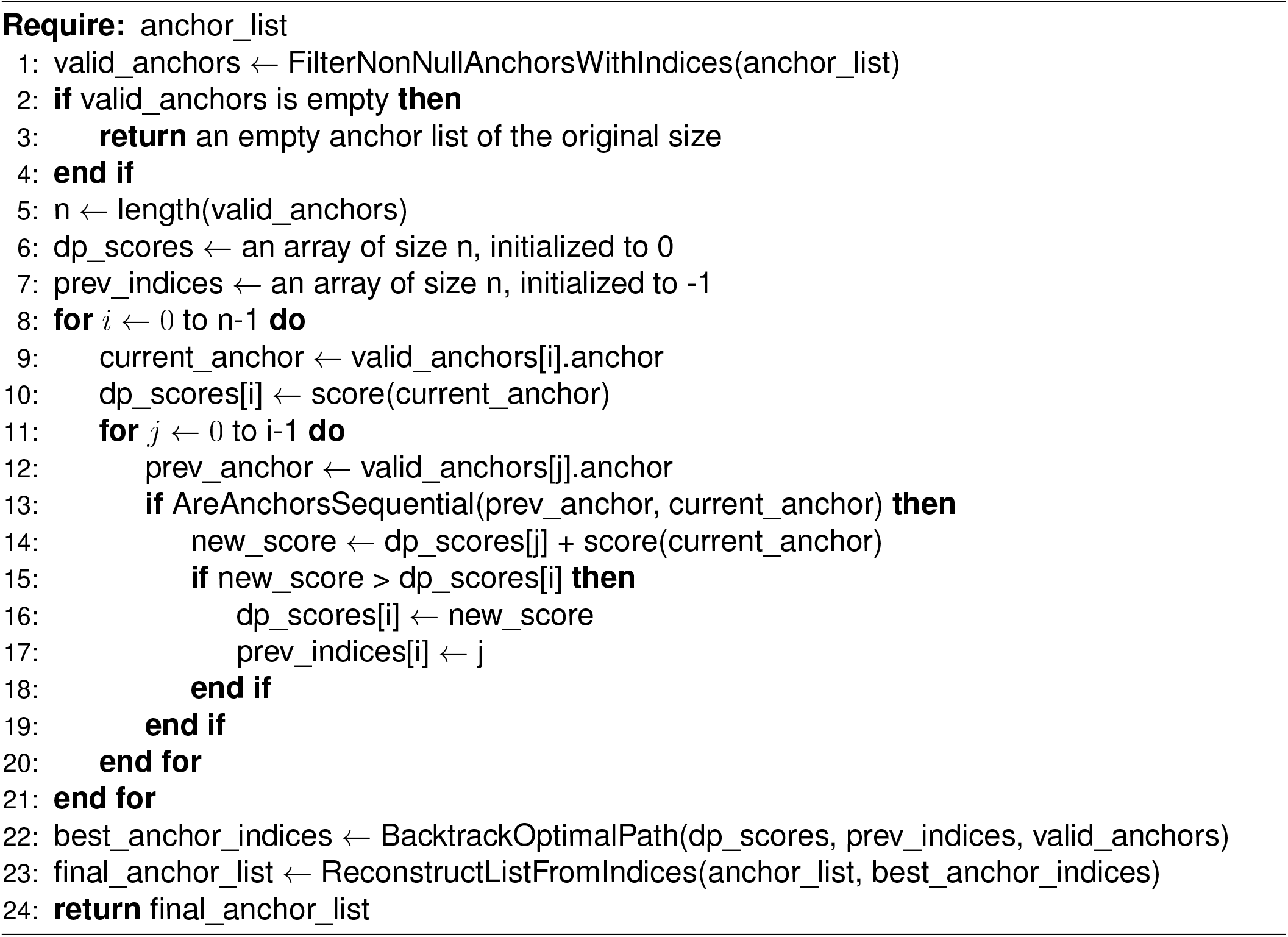

#### Algorithm 4 Merge Path into Graph

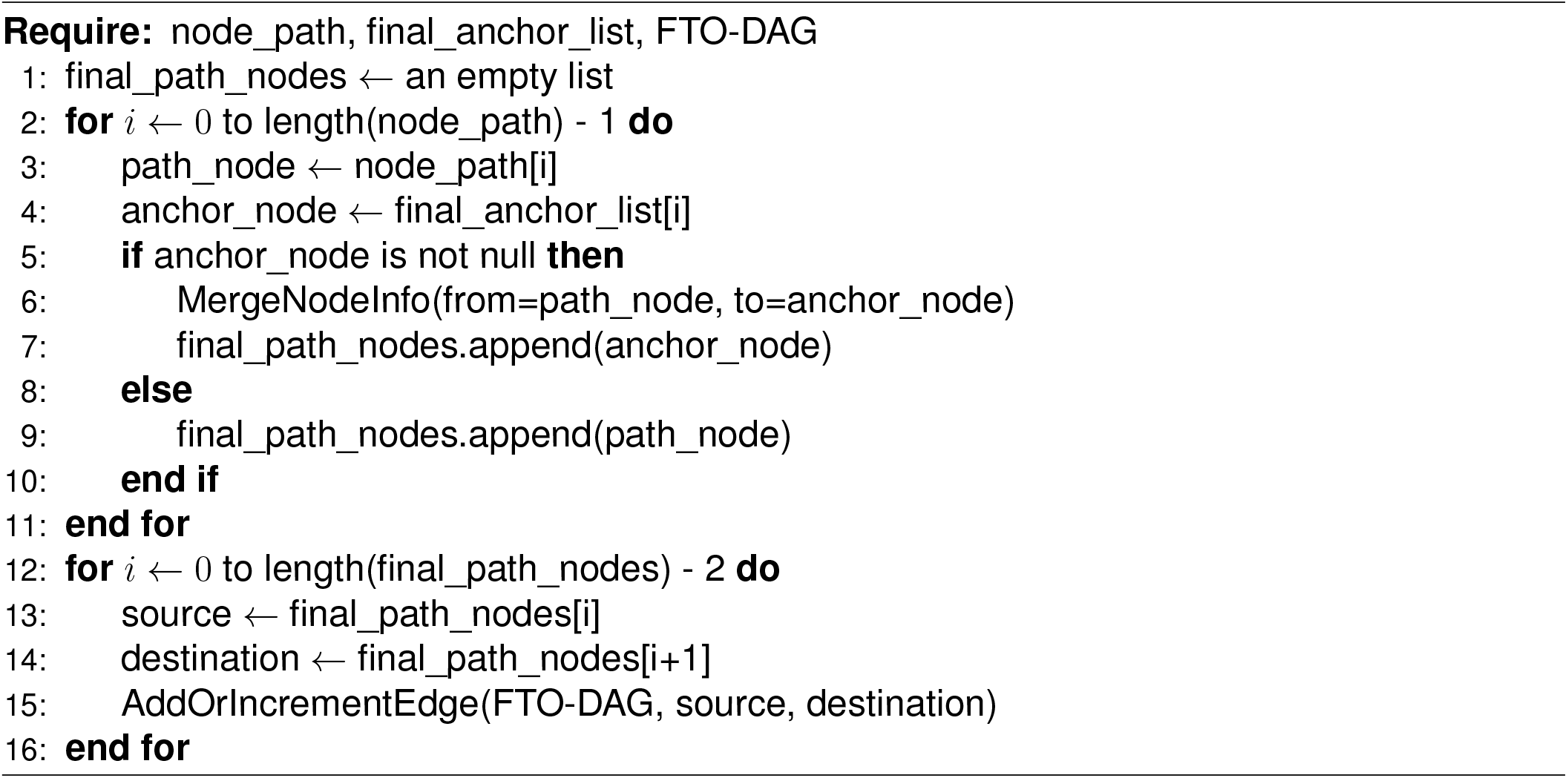

#### Algorithm 5 Topological Sort Guided Multi-process Traversal

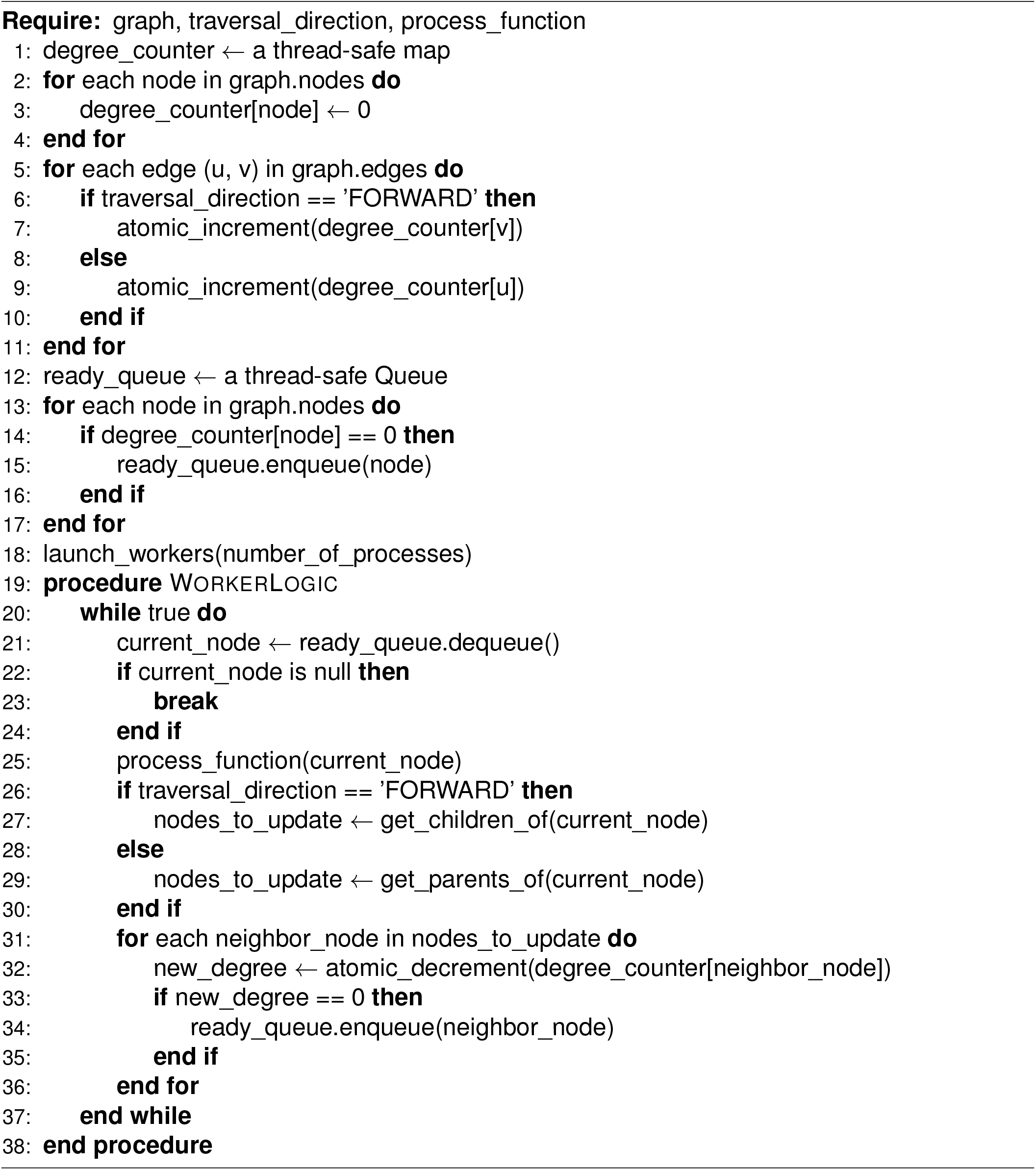

#### Algorithm 6 Maximum Weight Path Calculation

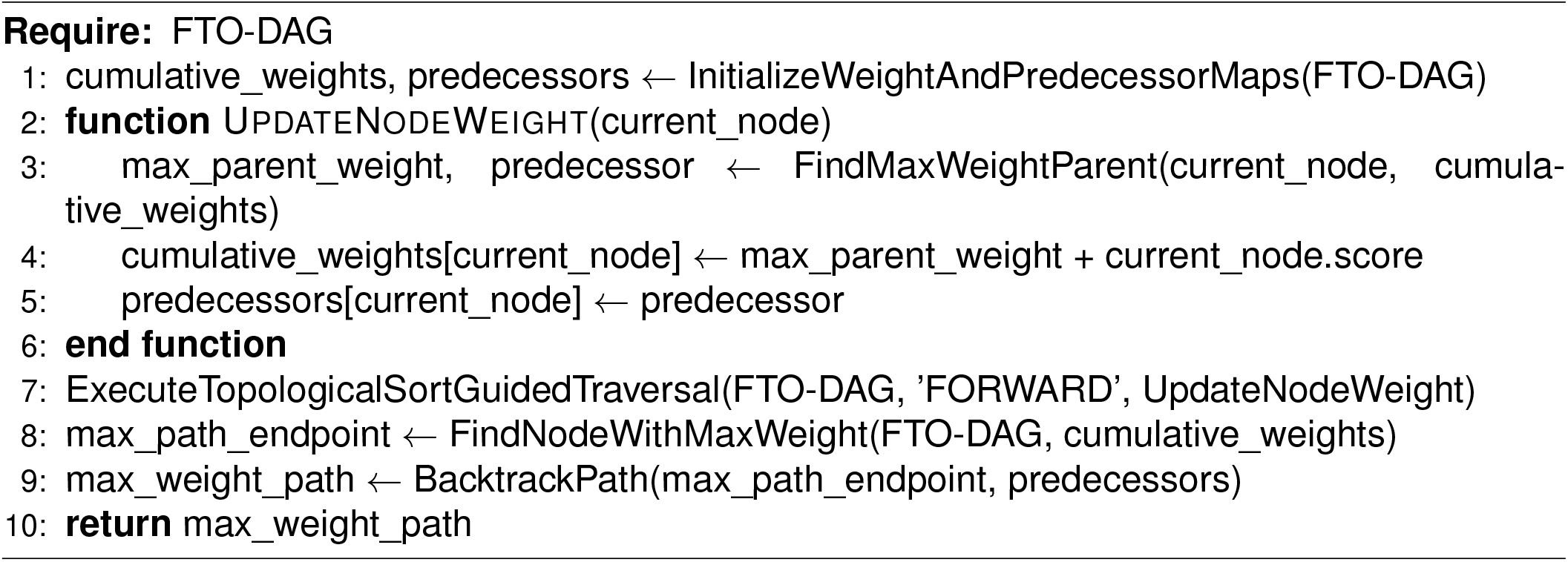

#### Algorithm 7 Fragment Resolution Adjustment

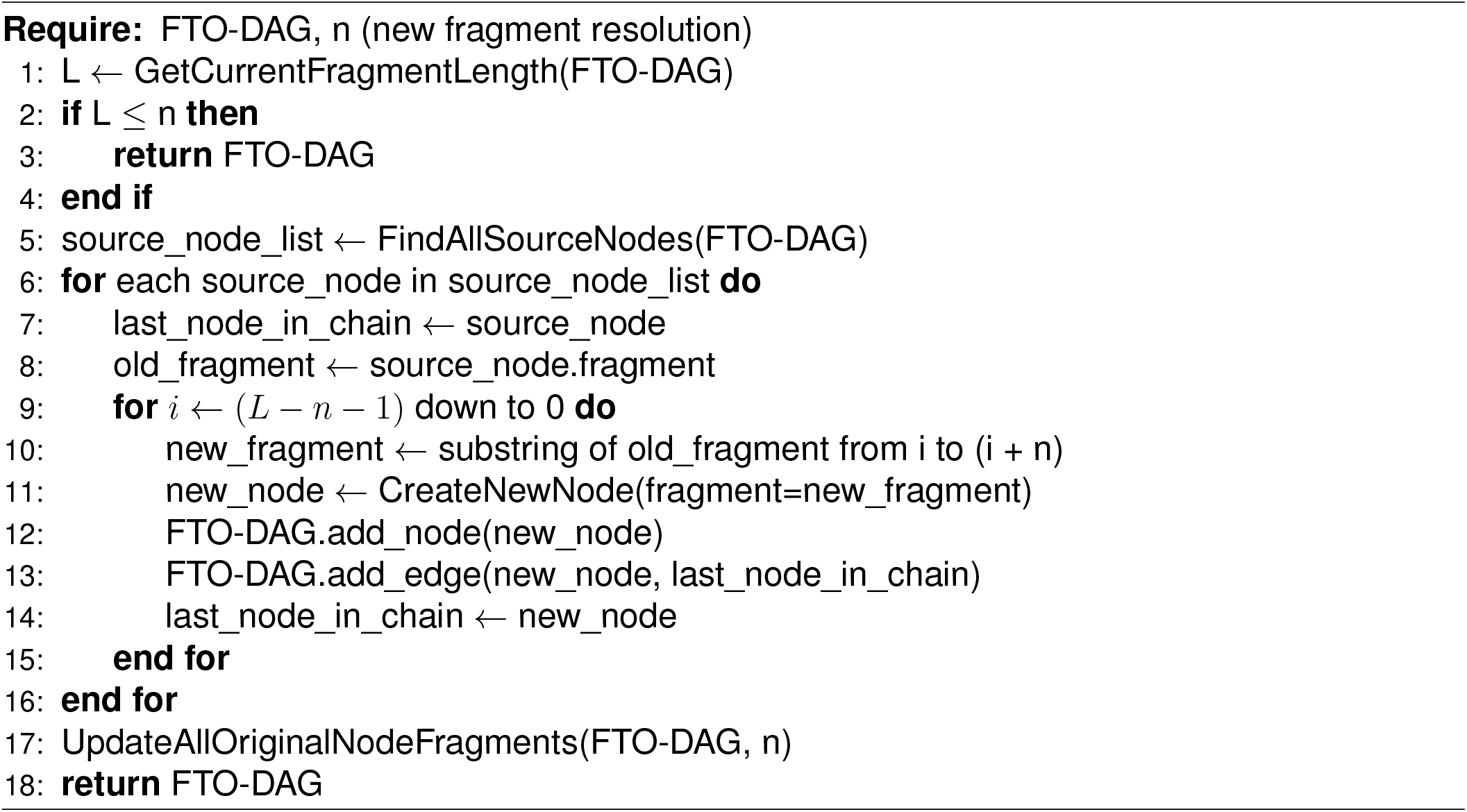

#### Algorithm 8 Graph Structure Optimization

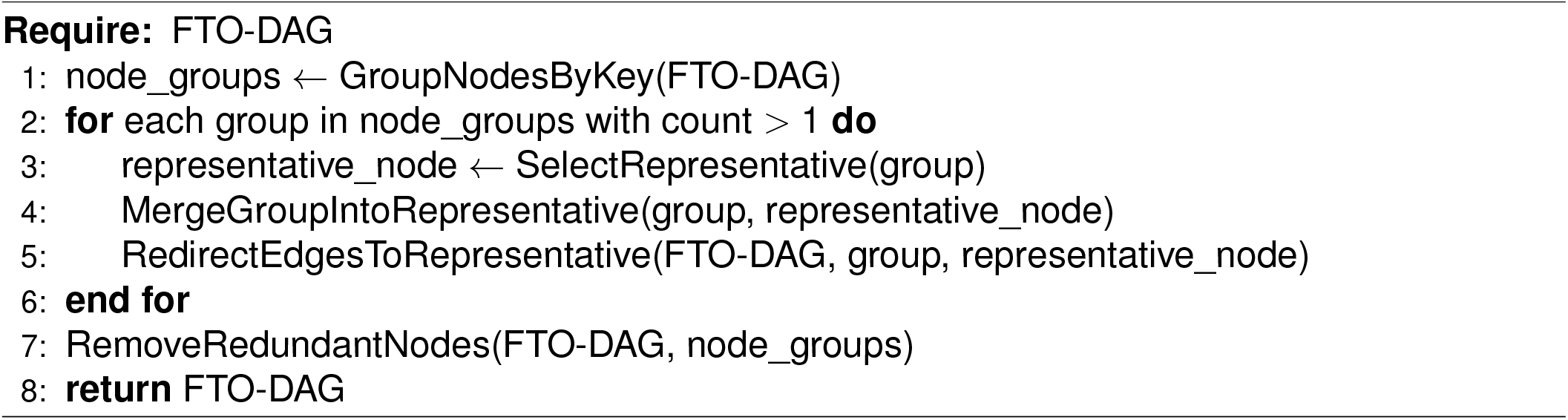

#### Algorithm 9 Topological Coordinate Calculation

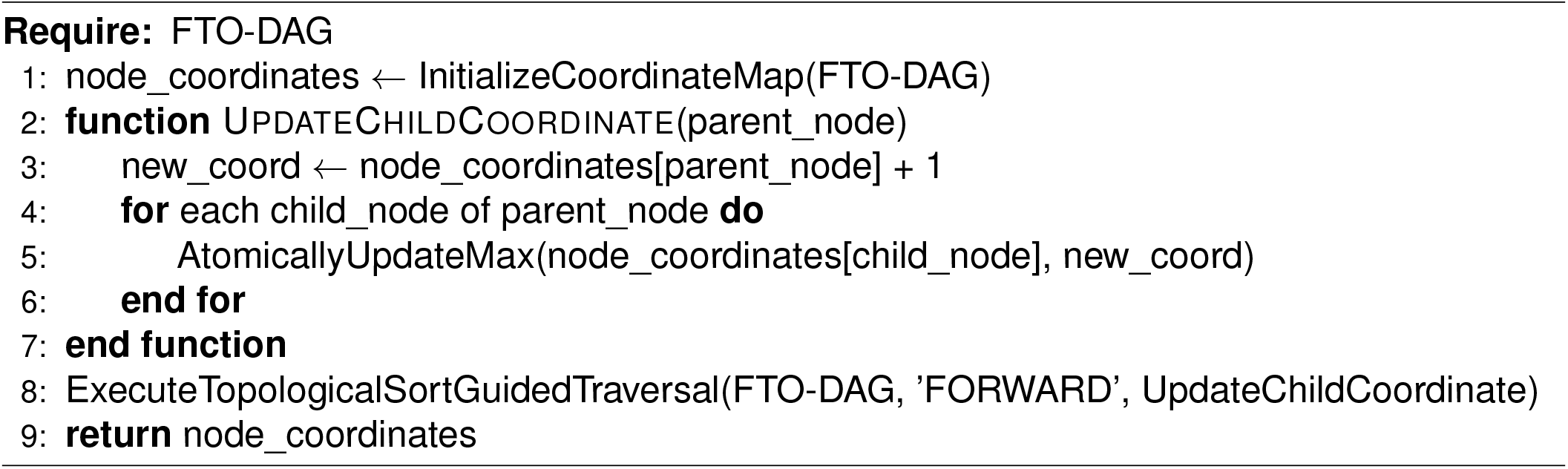

#### Algorithm 10 Sequential Edge Filtering

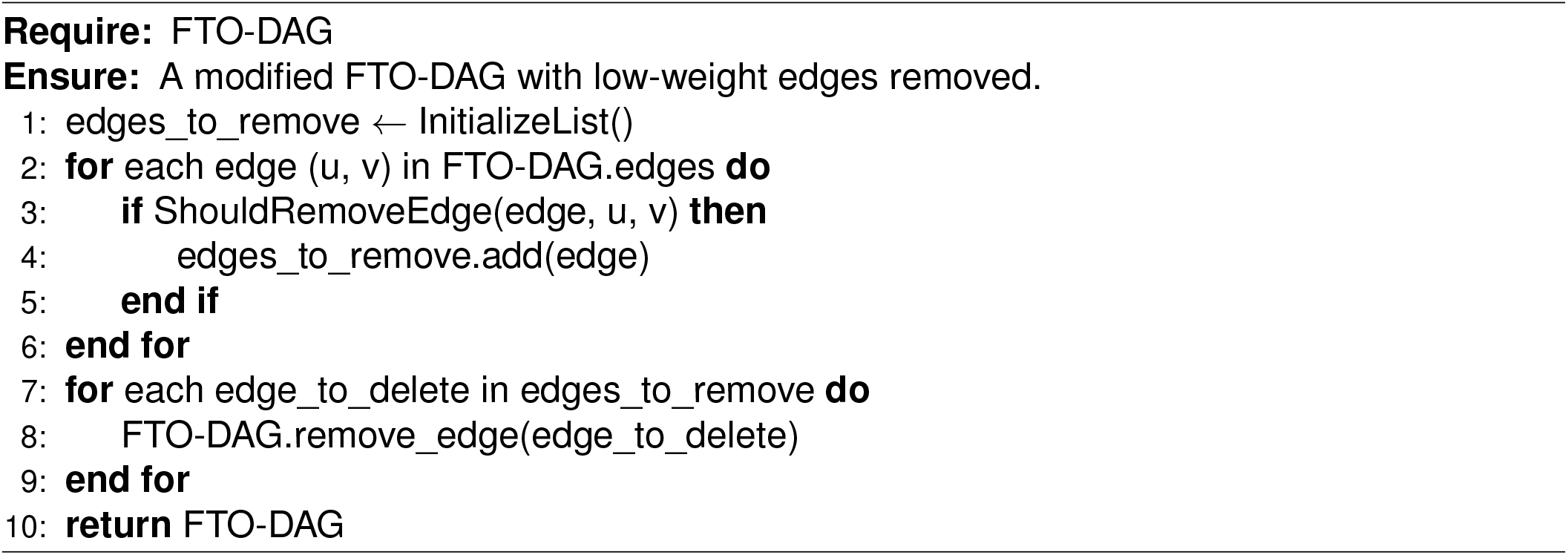

#### Algorithm 11 Graph Coarsening by Collapsing Linear Chains

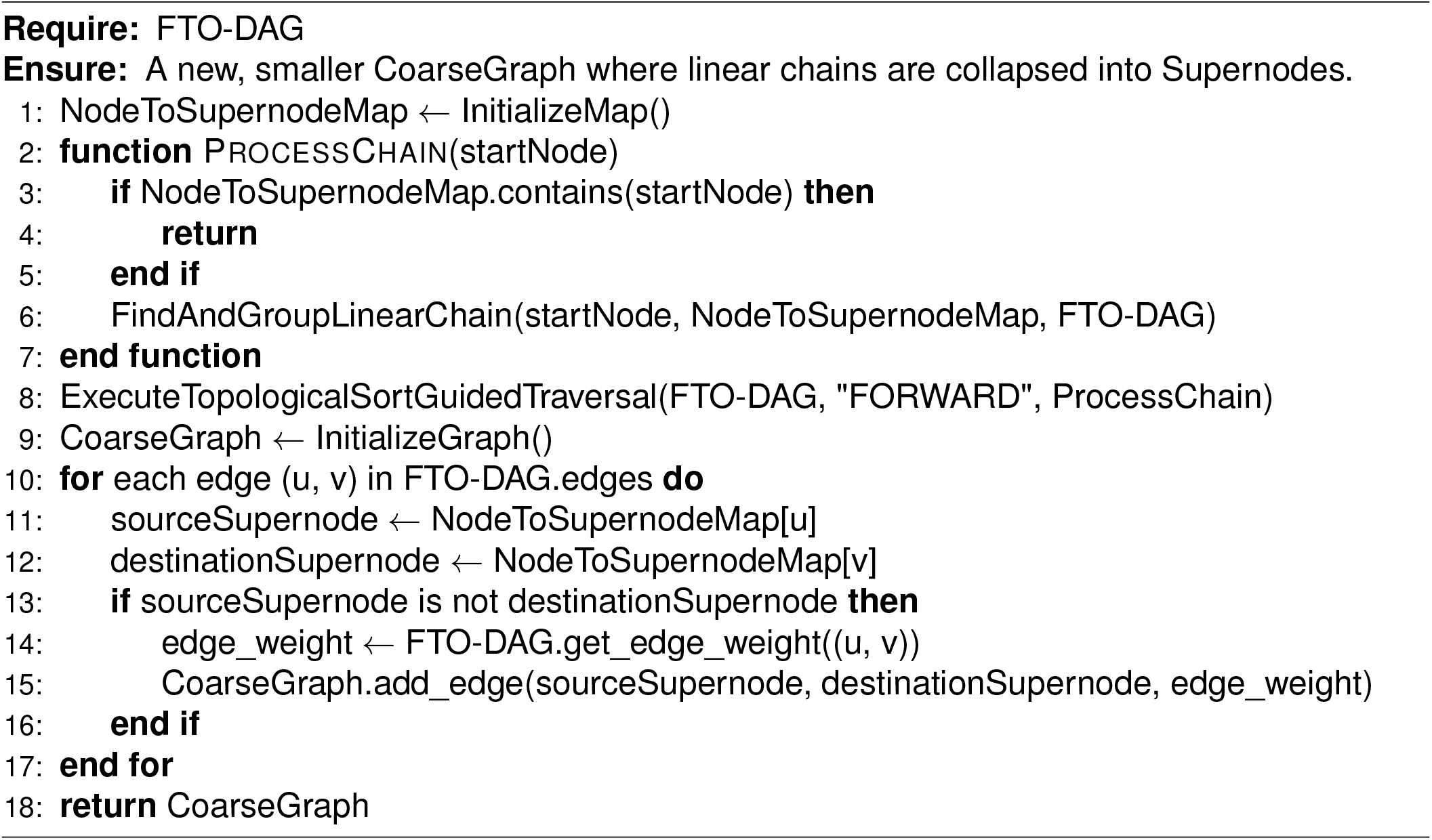

#### Algorithm 12 Path Statistics Analysis by Subgraph Coordinates

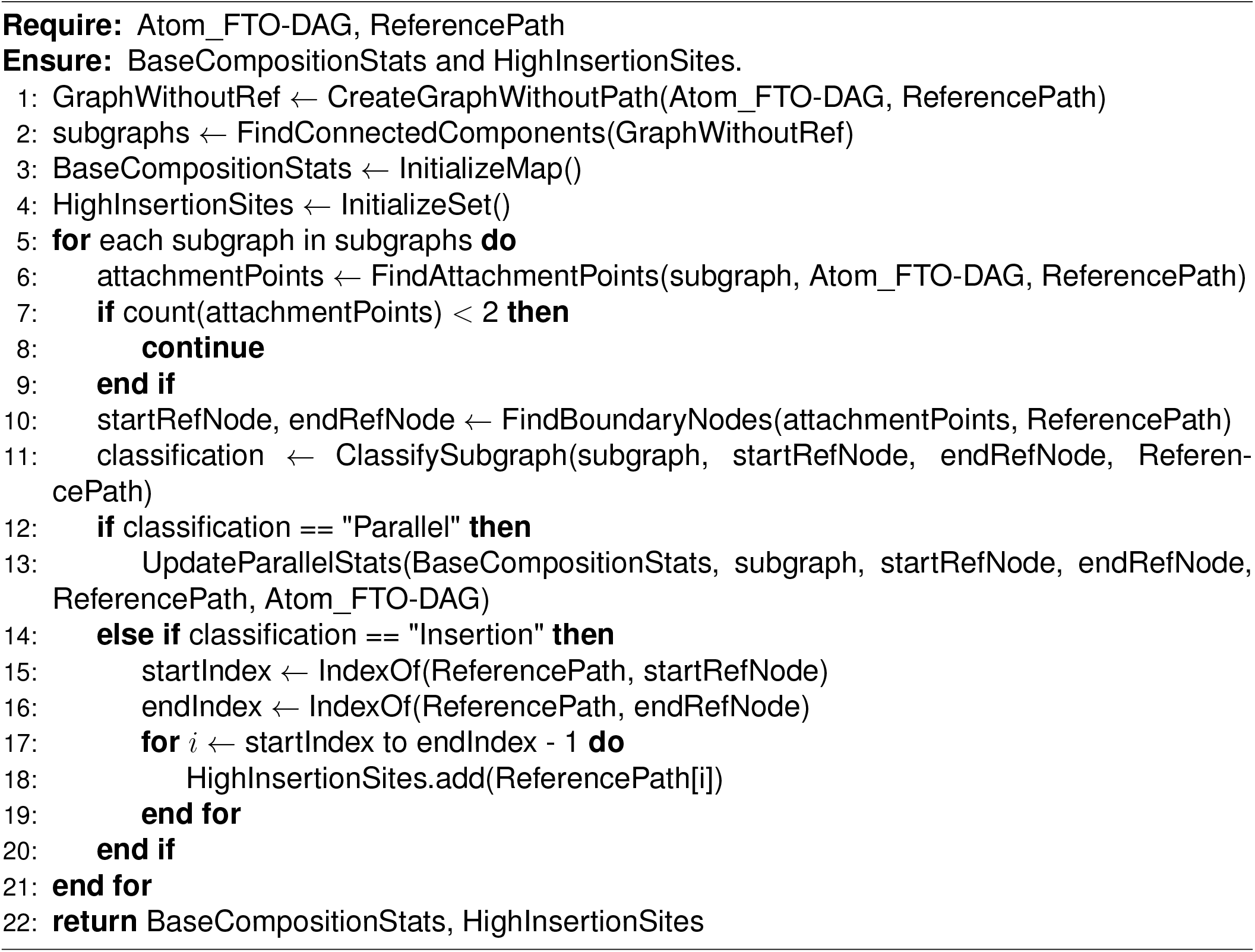

#### Algorithm 13 Iterative Baum-Welch for DAGs (Master)

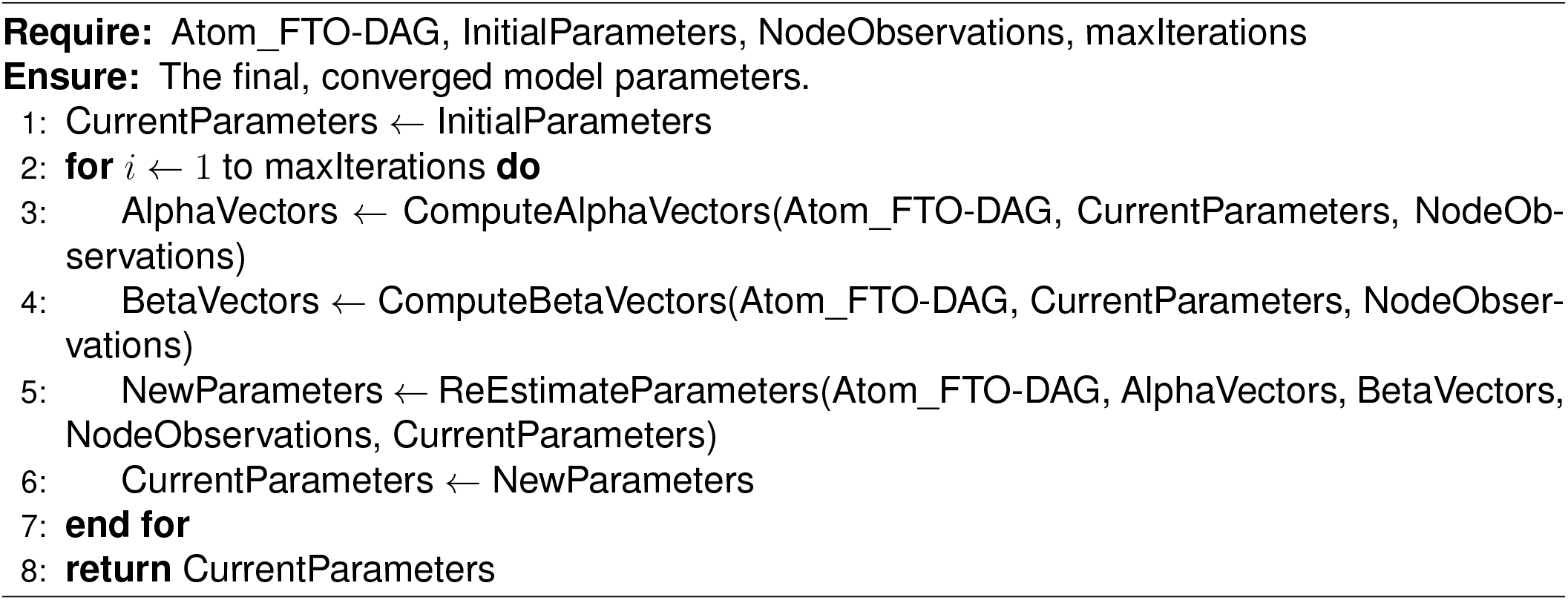

#### Algorithm 14 Compute Alpha Vectors (Forward Pass)

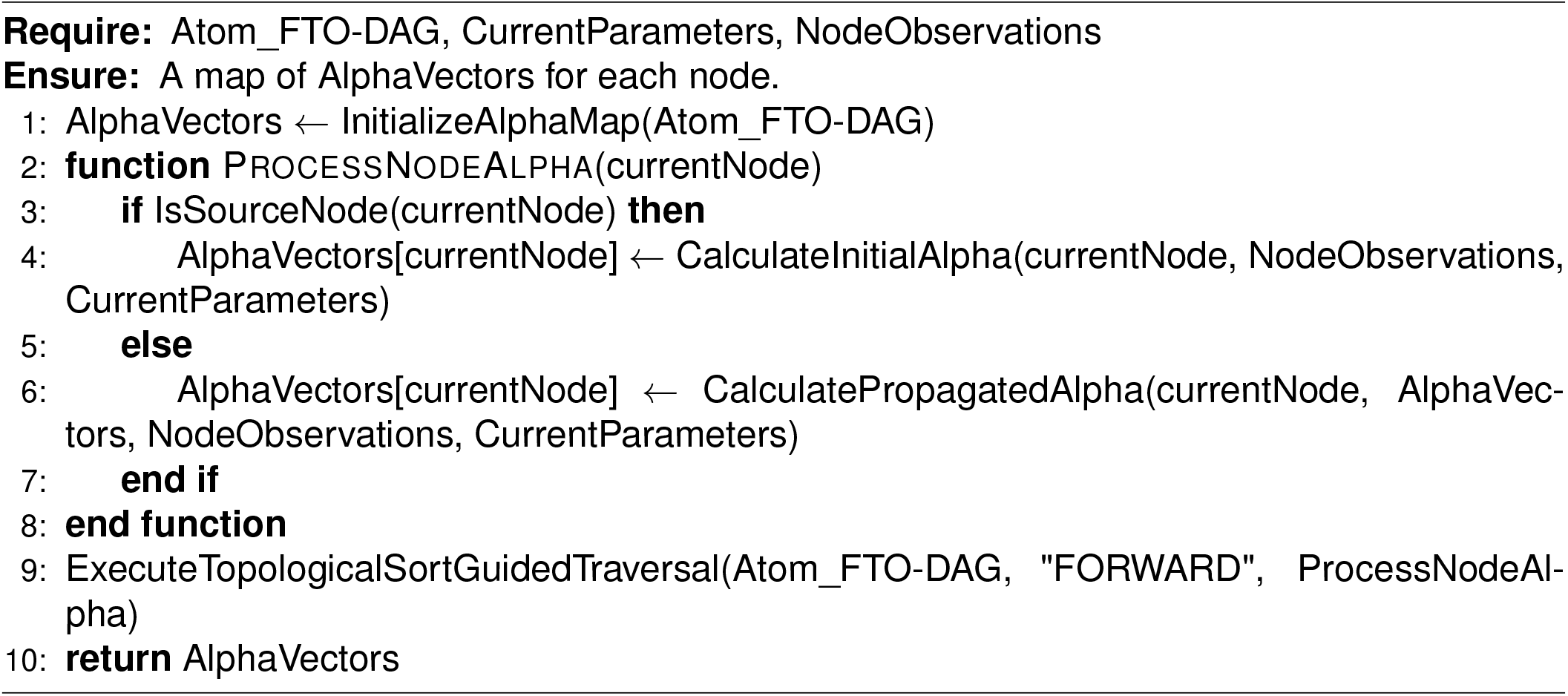

#### Algorithm 15 Compute Beta Vectors (Backward Pass)

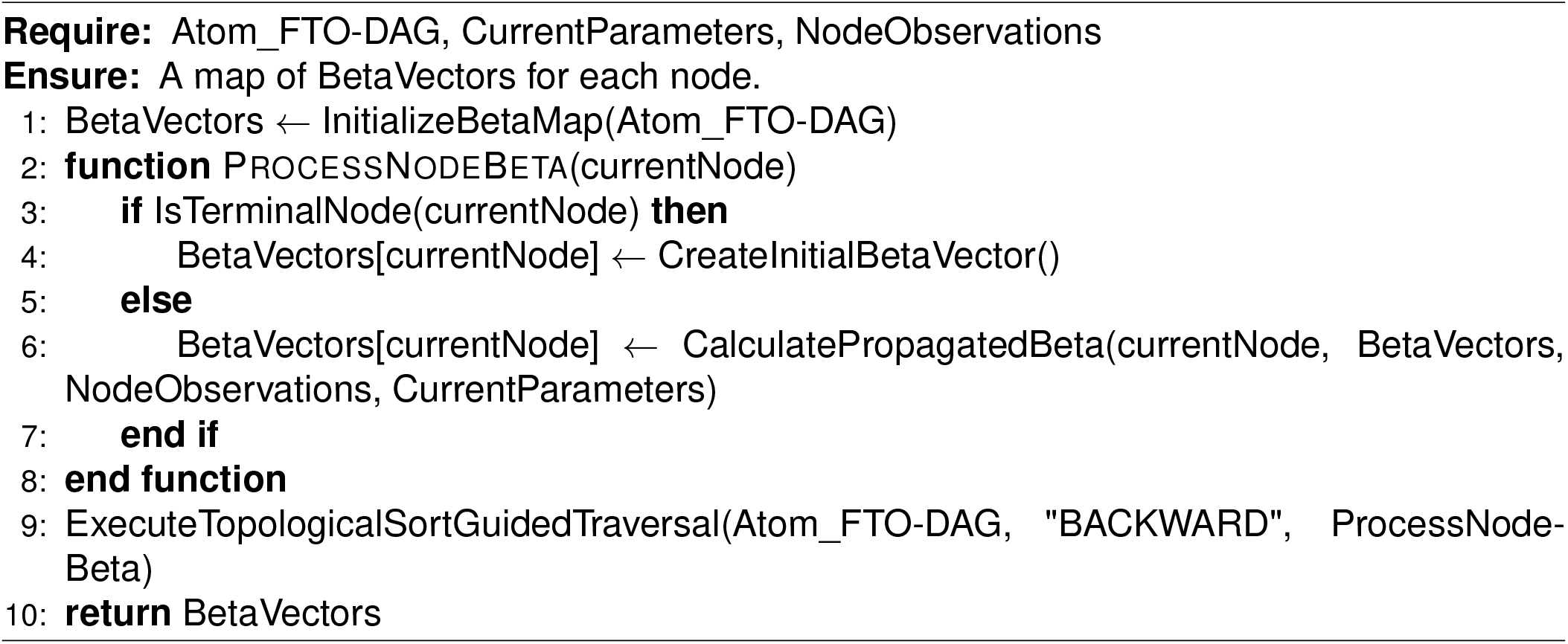

#### Algorithm 16 Re-Estimate Parameters (M-Step)

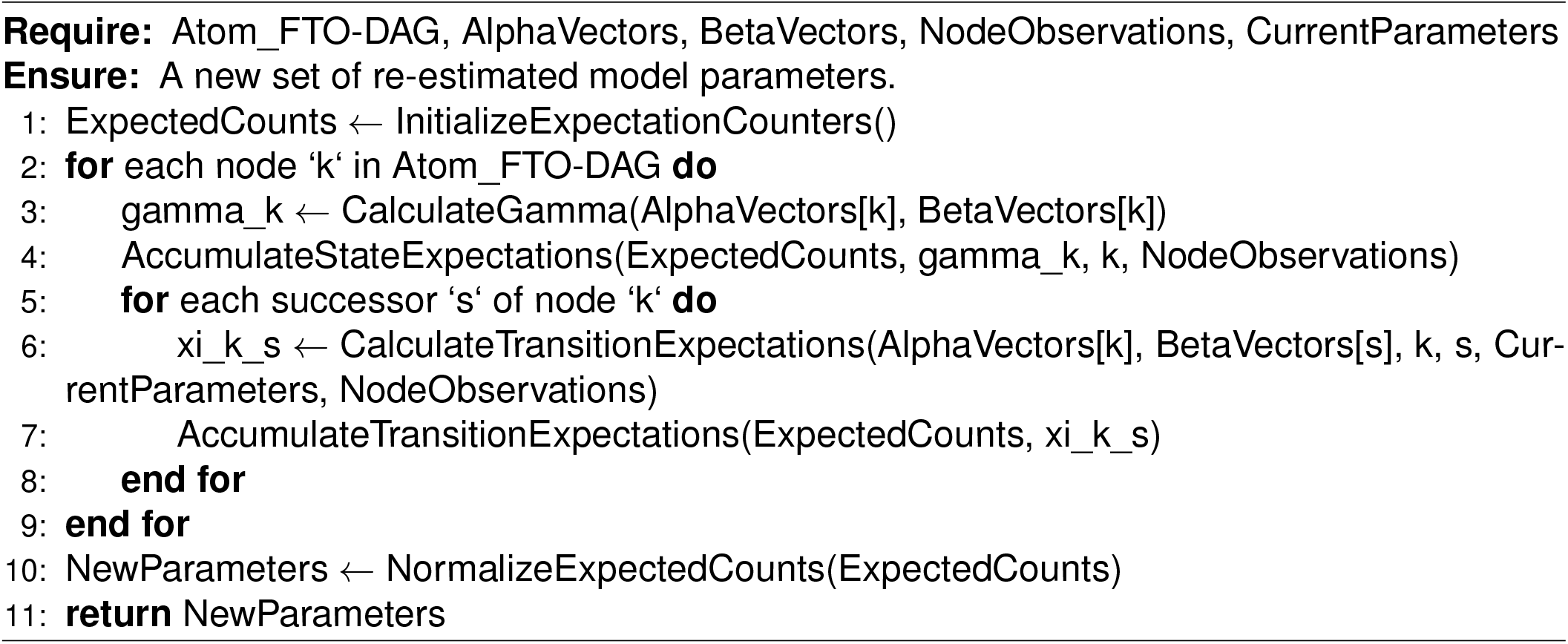

#### Algorithm 17 DAG-Viterbi for DAGs (Master)

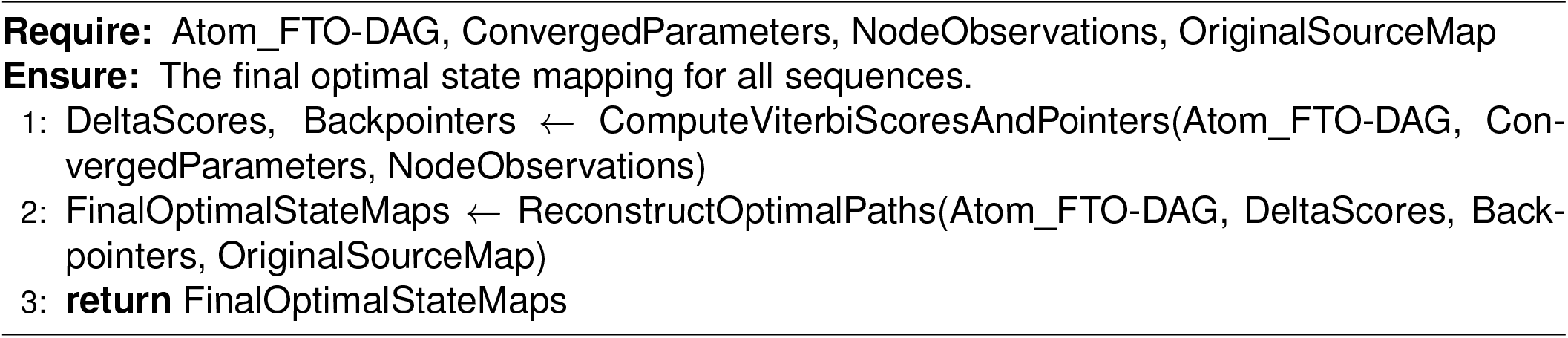

#### Algorithm 18 Compute Viterbi Scores and Pointers (Delta Pass)

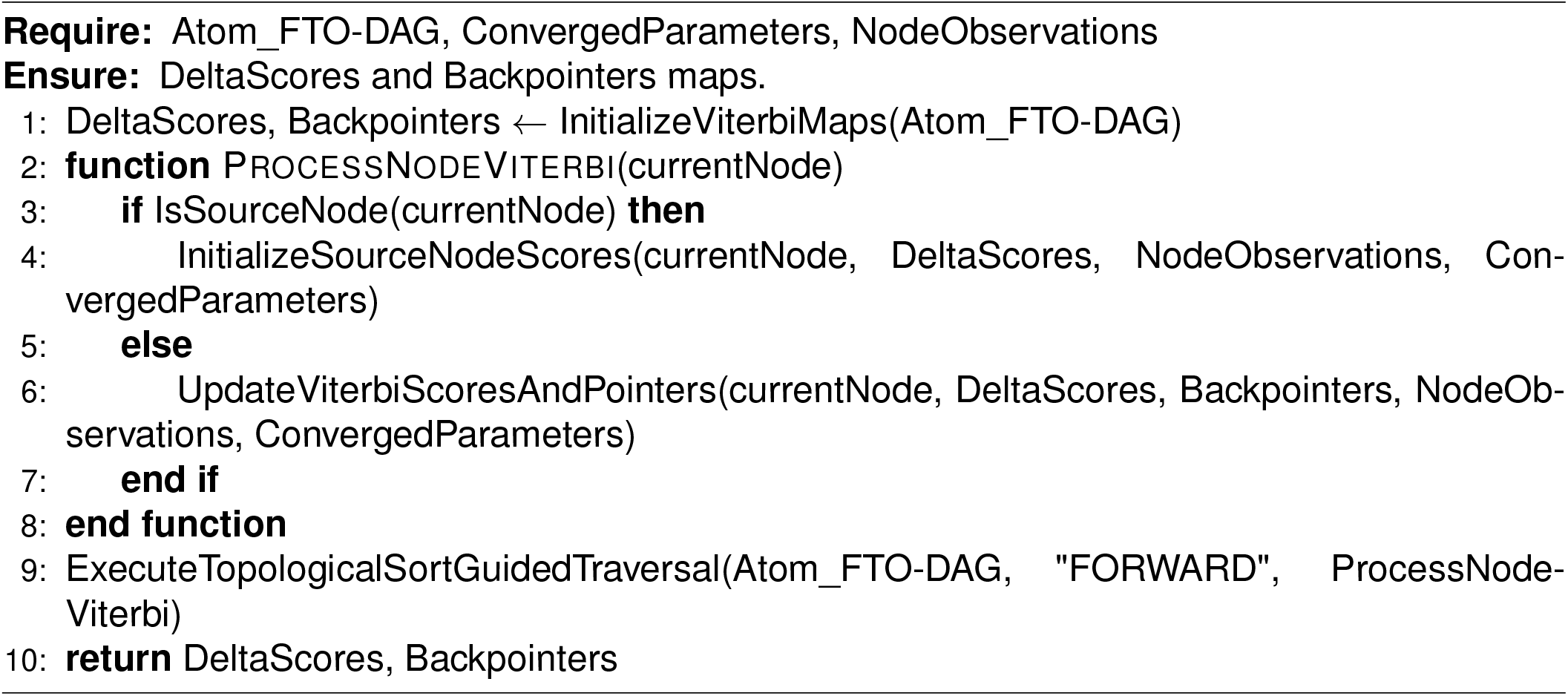

#### Algorithm 19 Reconstruct Optimal Paths (Traceback)

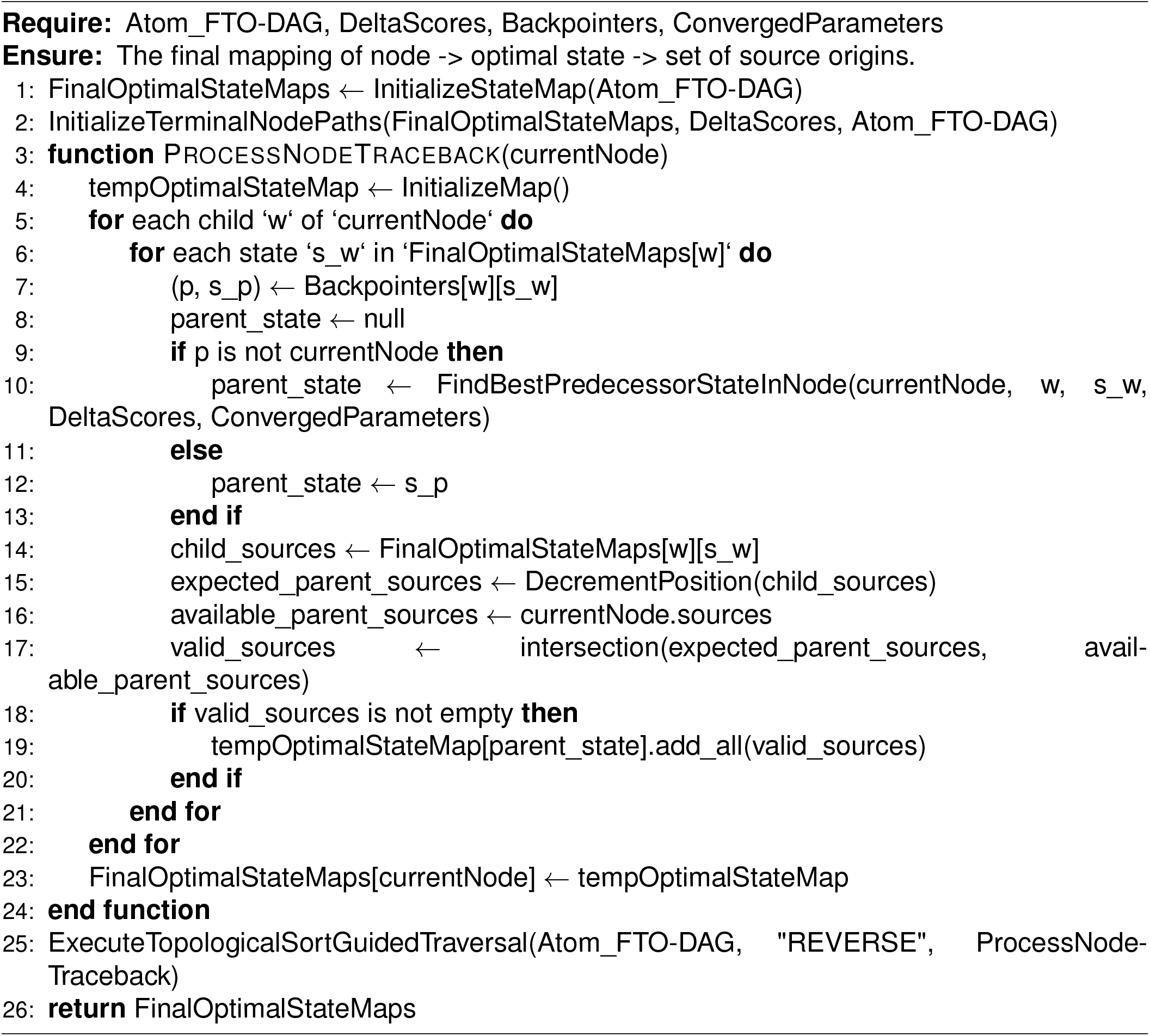

#### Algorithm 20 Multiple Sequence Alignment Generation

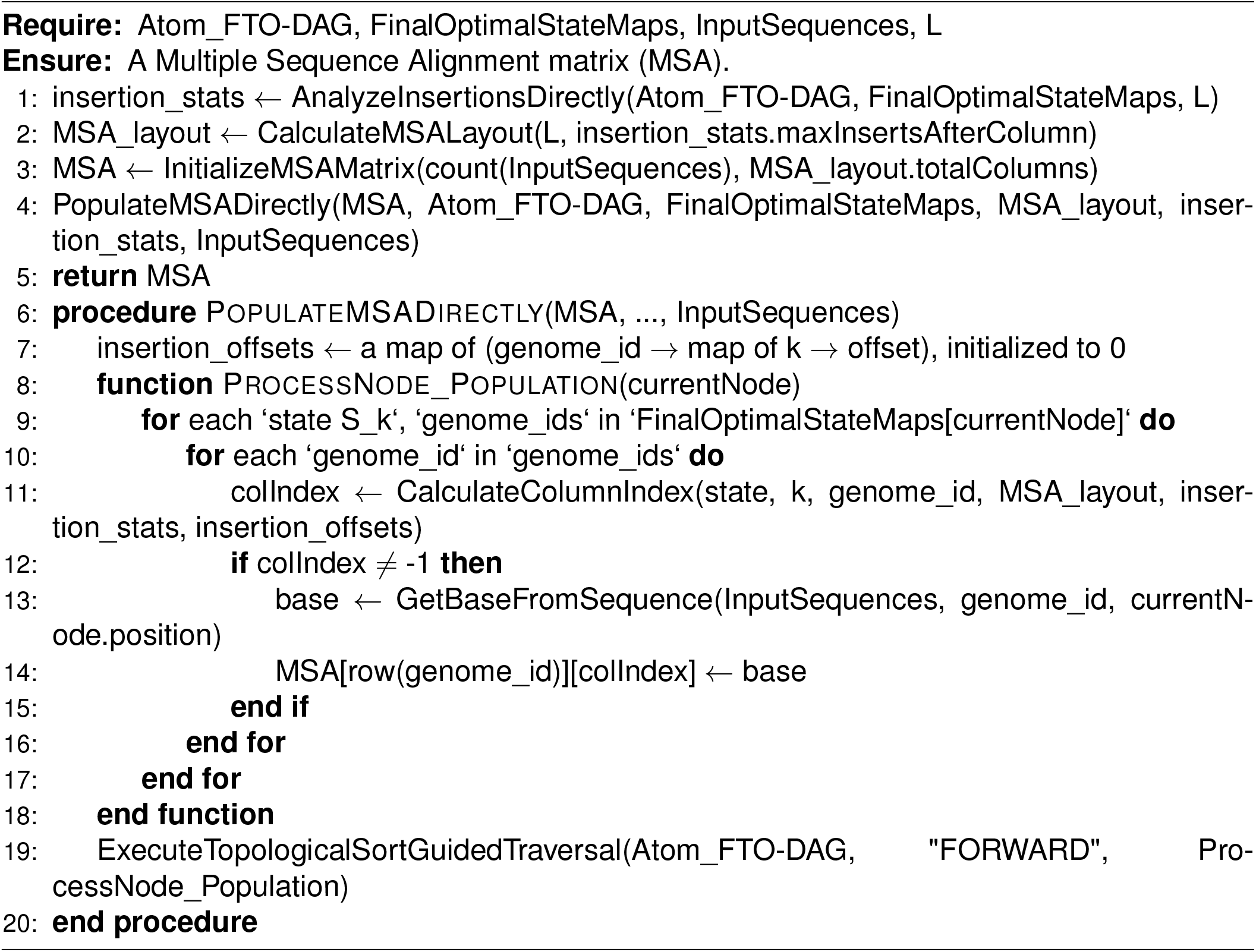

#### Algorithm 21 Tiled Computation Heuristic

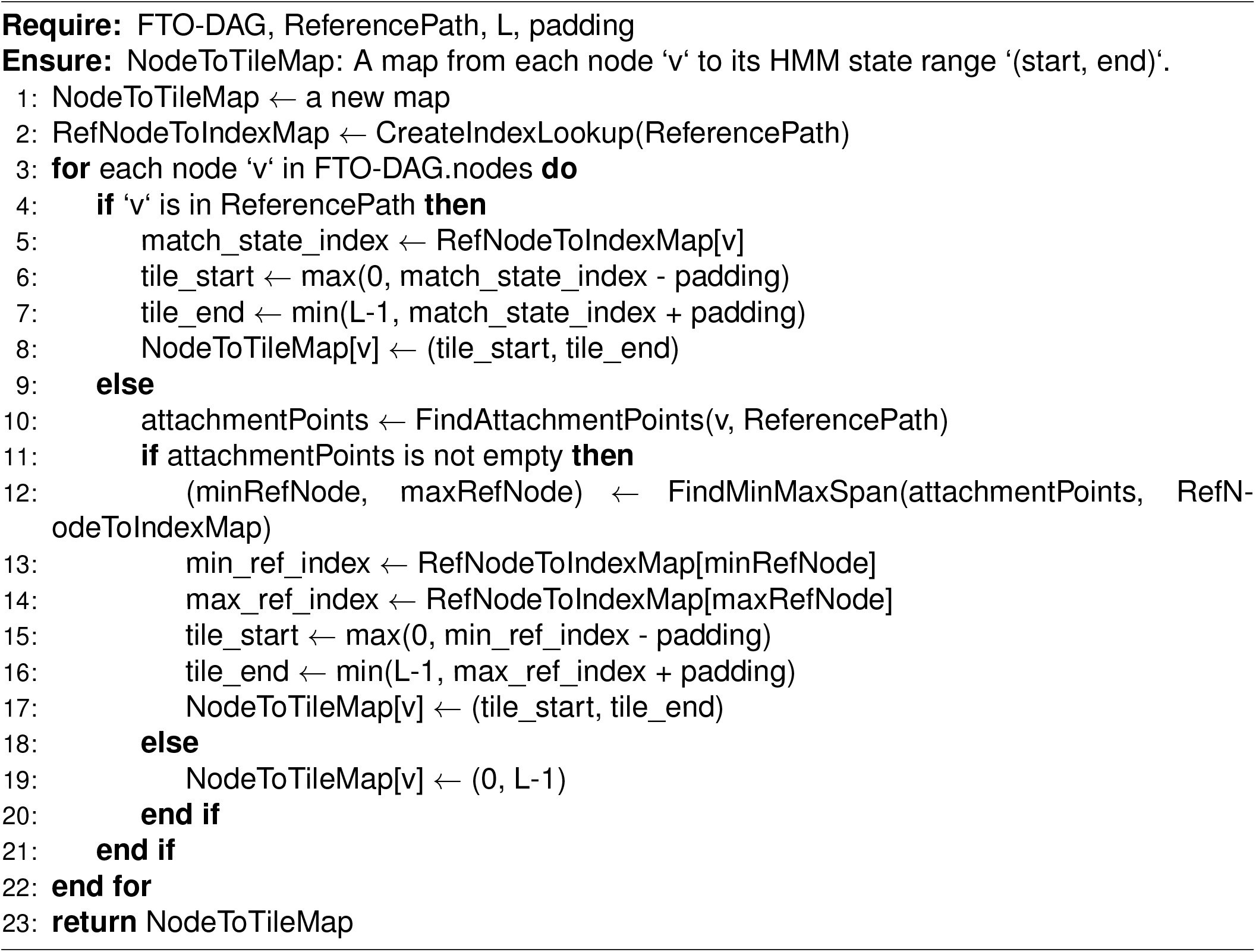

