## Supplemental texts and figures for "Accurate Multiple Sequence Alignment of Ultramassive Genome Sets"

### SUPPLEMENTAL INFORMATION

#### S1 Redundancy in Existing MSA Algorithms on Similar Sequences

##### S1.1 Redundancy Eliminated by Dynamic Programming

For two sequences $X$ and $Y$ of lengths $L_x$ and $L_y$, a brute-force alignment approach would enumerate all possible relative positionings to find the one(s) with the highest score. The maximum possible length of the resulting alignment is $L_{tot} = L_x + L_y$. Without loss of generality, we may assume $L_x \geq L_y$ and the total number of configurations to be scored are within a range of $\sum_{i=0}^{L_y} C_i^{L_x+i} C_{L_x-L_y}^{L_x}$. This formula may be easily derived by considering the arrangement of gaps only. The number of candidates to be scored is computationally prohibitive for sequences of several hundred bases or more. Needleman and Wunsch introduced dynamic programming to pairwise sequence alignment, eliminating the massive redundancy inherent in the aforementioned brute-force scoring scheme. Specifically, pairwise alignment is simplified to the task of filling a matrix, as illustrated in the dynamical programming formulation (Figure S1(a)) and its matrix representation (Figure S1(b)). Consequently, both the space and time computational complexities are reduced to $O(L_x L_y)$.

##### S1.2 Redundancy Generated by Existing Algorithms on Similar Sequences

Pairwise sequence alignment, which forms the most critical foundation for progressive, iterative, and consistency-based MSA, generates a vast amount of computational redundancy when applied to a set of highly similar sequences. Specifically, within a set of N sequences, if a fragment p ($Seg_p$, shown in green in Figure S2(a)) exists in each of the $n_p$ sequences in a set P, and a fragment q ($Seg_q$, shown in red in Figure S2(a)) exists in each of the $n_q$ sequences in a set Q, then the corresponding alignment block (shown in purple in Figure S2(a)) will be repeated $n_p \times n_q$ times if sets P and Q are disjoint; $n_p(n_p-1)/2$ times if sets P and Q are identical; and a number of times that varies between these two values if the sets are neither identical nor disjoint. Since the sizes of sets P and Q can both range from 1 to N, the maximum possible number of repetitions for a single alignment block is $N(N-1)/2$, which amounts to approximately 5 * $10^{11}$ for a set of one million sequences. Consequently, some computational blocks are repeated hundreds to trillions of times, leading to a colossal waste of computational resources (Figure S2(b)). Even greater amounts of redundancy are generated during the construction of guide trees for progressive and iterative MSAs, and still more massive redundancy occurs when building the library for consistency-based MSAs. By successfully eliminating this redundancy, our algorithmic framework achieves both a significant speedup and a notable improvement in accuracy compared to quadratic-scaling methods, such as the fastest options in MAFFT.

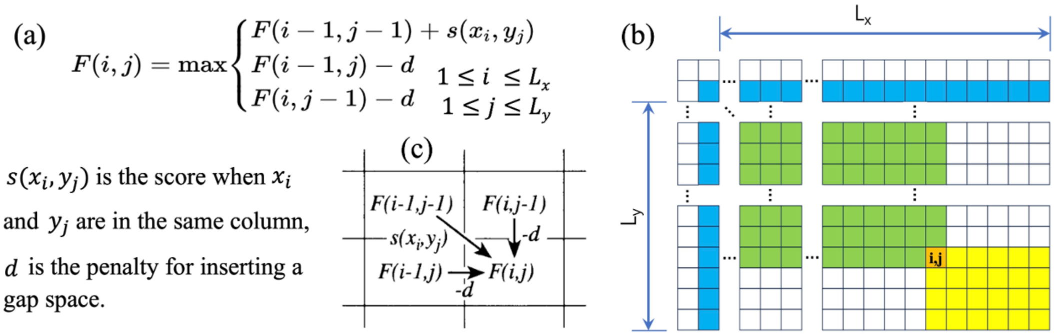


**Figure S1: Repetition Riddance by Dynamic Programming**. (a) The formulation developed in Needleman-Wunsch algorithm. (b) Matrix representation of the dynamical programming for two sequences $x$ and $y$ with lengths $L_x$ and $L_y$ , the 2nd row and the 2nd column (in blue) are initialization row and column representing a gap. To calculate score $F(i,j)$ of the alignment between $x_{1,2,\cdots,i}$ and $y_{1,2,\cdots,j}$, all upper and left slots (shown in green) need to be calculated first, and the result is stored in $(i,j)$ slot in orange. Calling slots of $(i,j)$ (all lower and right slots shown in yellow) need the results of $(i,j)$, so caching of which removes repetition by the number of corresponding calling slots.

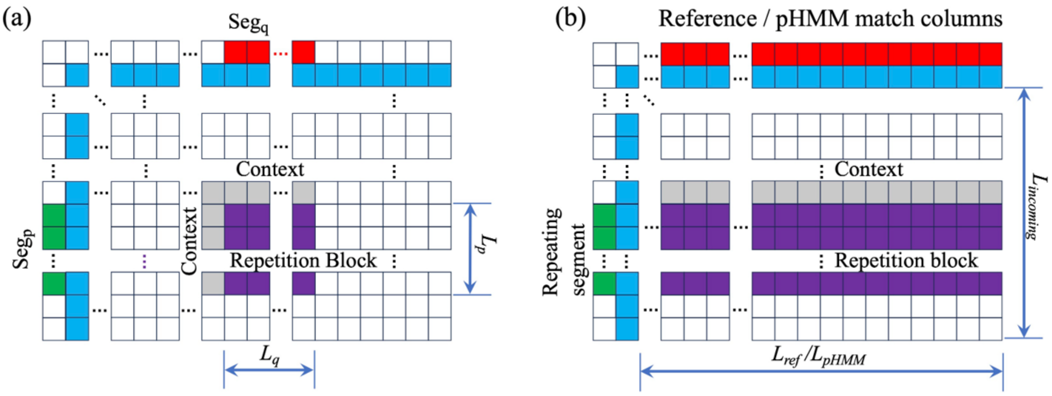


**Figure S2: Repetition Block in Alignment of Similar Sequences**, related to STAR Methods. (a) Pairwise alignment scenario, where two segments $Seg_p$ and $Seg_q$ with lengths $L_p$ and $L_q$ (green and red slots) define a repetition block of size $L_p \times L_q$ (purple slots), it is important to note that the content of these blocks are determined not only by their defining segments, but also by their context (preceding bases in two sequences) shown as gray slots. (b) PHMM or SR-align pair scenario, all sequences are aligned with the PHMM state positions or the given reference sequence (both are represented by the same set of red slots), a repeating segment is shown in green. Similar to the pairwise scenario, the content of repetition block (purple slots) corresponding to the repeating segment depends not only on the repeating segment and PHMM/reference, but also on the context (the preceding base of the repeating segment) shown as gray.

#### S2 Compressed Sparse Alignment Format (.npz)

The compressed sparse alignment format (.npz) is a highly efficient representation of MSAs (MSAs), designed to minimize storage requirements and computational overhead. This format is compatible with both DAG-align and SR-align algorithms, making it a versatile choice for ultra-large-scale genomic datasets. Below, we describe the structure and components of the .npz format in detail.

**Structure of the .npz File** The .npz file contains two main arrays:

- `namelist`: An array of strings of length $N$, storing the ordered names of all $N$ sequences in the alignment.
- `align`: An array of lists of length $L + 1$, where $L$ is the total number of columns in the alignment.
  - For the first $L$ elements (from index $0$ to $L - 1$), each element is a list representing one column of the alignment. The structure is: `[ConsensusCode, [VariantCode1, [index_list_1]], [VariantCode2, [index_list_2]], ...]`
    - `ConsensusCode`: An integer representing the most frequent base/residue in that column.
    - Each subsequent element (e.g., `[VariantCode1, [index_list_1]]`) is a two-element list where the first item is an integer for a variant base, and the second is a NumPy array of integer indices specifying which sequences (referencing `namelist`) have this variant in this column.
  - The final element (at index $L$) is a single integer $N$, representing the total number of sequences.

**Base/Residue Encoding** The integer codes for bases/residues are as follows:

```
0: '-', 1: 'A', 2: 'T', 3: 'C', 4: 'G', 5: 'R', 6: 'Y',
7: 'M', 8: 'K', 9: 'S', 10: 'W', 11: 'H', 12: 'B', 13: 'V',
14: 'D', 15: 'N'
```

**Advantages of the .npz Format**

- **Storage Efficiency:** By storing only the differences relative to the consensus sequence, the .npz format significantly reduces the size of alignment files, especially for highly similar sequences.
- **Fast Access:** The sparse representation allows for rapid reconstruction of specific columns or rows of the alignment without the need to load the entire dataset into memory.
- **Compatibility:** The format is designed to be easily parsed and manipulated using standard Python libraries such as NumPy, making it accessible for downstream analysis.

**Usage in DAG-align and SR-align** Both DAG-align and SR-Align algorithms support the .npz format as an output option. This ensures that users can seamlessly integrate alignments generated by either method into their workflows, leveraging the format's efficiency and flexibility.

#### S3 Single-Reference Alignment (SR-align)

Performing pairwise alignments of all sequences against a single selected reference sequence (Figure S3), while ignoring all relationships among the non-reference sequences, serves as a very crude approximation of an MSA. This approach is referred to as "SR-align" in the main text. This approximation was used as a baseline for comparison. For all test sets, the first sequence is chosen as the reference by default. After all query sequences are pairwise aligned to the reference, they are assembled into a final MSA based on the reference coordinates. Bases in a query sequence that align to a specific reference base (as matches or mismatches) are placed in the corresponding column of the MSA. Insertions relative to each of the reference sequence base/residue create new columns in the final alignment. When multiple query sequences have insertions at the same position relative to the reference, these insertions are left-aligned within the newly created block of columns, thus completing the MSA (Figure S3(c)).

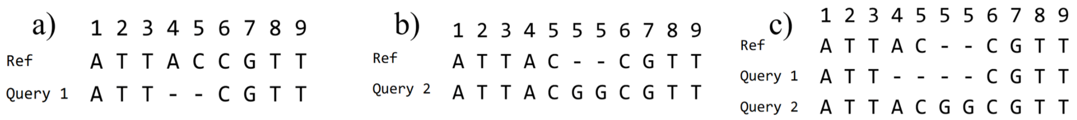


**Figure S3. Principle and Workflow of Single-Reference Alignment (SR-align)**. The figure schematically illustrates the SR-align method, a rapid approximation for MSA. (a) A query sequence is aligned to a selected reference sequence. (b) This process is repeated independently for all other query sequences against the same reference. (c) A final MSA is assembled by mapping all query sequences onto the reference sequence's coordinate system. Insertions relative to the reference create new columns, and when multiple sequences have insertions at the same locus, they are left-aligned within the newly created block of columns.

#### S3.1 SR-align: A High-Performance Python Implementation

`SR-align` is a high-performance, single-reference alignment tool designed for the rapid MSA of massive genomic datasets. Its primary goal is to efficiently align a large number of query sequences against a single reference sequence and generate both a standard alignment file (for smaller datasets) and a highly compressed sparse alignment format suitable for ultra-large-scale data analysis.

**Core Features** The core features of the script include:

- **High-Performance Alignment Core:** Utilizes the `parasail` C library(`https://github.com/jeffdaily/parasail`), which implements SIMD-optimized Smith-Waterman and Needleman-Wunsch algorithms, offering alignment speeds far exceeding pure Python implementations.
- **Multi-process Parallelization:** Leverages Python's `multiprocessing` module to distribute alignment tasks across multiple CPU cores, significantly reducing the time required to process large datasets.
- **Memory Optimization:** Employs a stream-based reading of the input file and a difference-based result collection strategy. During parallel computation, worker processes return only the "difference information" (mismatches, insertions, deletions) between sequences rather than the full aligned sequences, substantially reducing inter-process communication overhead and memory consumption.
- **Adaptive Output Strategy:** The script provides two output formats. For datasets with 40,000 or fewer sequences, it generates a standard MSA (MSA) in FASTA format. For all datasets, it saves the alignment in a custom, highly compressed sparse `.npz` format to manage disk space effectively.

##### Core Dependencies

- **BioPython:** For efficient parsing and writing of FASTA-formatted sequence files.
- **parasail:** Provides high-performance sequence alignment functions.
- **NumPy:** Used for efficient array operations, especially when reconstructing alignment results and creating the sparse `.npz` file.
- **tqdm:** Offers a user-friendly command-line progress bar to monitor the processing status.

**Workflow** The script's execution flow is carefully designed for maximum efficiency:

1. **Initialization and Reference Loading:** Upon launch, the script first reads the first sequence from the input FASTA file to serve as the reference. It then creates the necessary scoring matrix for alignment based on predefined scores. The output directory specified by the `-o` argument is created if it does not already exist. Output filenames are automatically derived from the input filename.
2. **Parallel Task Distribution:** The script creates a process pool (`multiprocessing.Pool`) with the number of processes specified by the user. Using an `initializer`, the reference sequence string, scoring matrix, and penalty values are pre-loaded into the global memory of each worker process. This critical optimization avoids the repetitive transfer of this large, read-only data for each alignment task.
3. **Streaming and Alignment:** The main process uses a generator function (`task_generator`) to read query sequences one by one from the input file, submitting each as an independent task to the process pool. A worker process receives a task, calls the `perform_alignment_task` function, which uses `parasail` to perform a global alignment against the pre-loaded reference.
4. **Difference Information Extraction:** After alignment, the `perform_alignment_task` function parses the alignment "traceback" to extract and return only the differences of the query relative to the reference. These differences (mismatches, insertions, deletions) are encoded as integers for efficiency.
5. **Result Aggregation and Statistical Analysis:** The main process asynchronously collects the difference information from all worker processes. Once all alignments are complete, the `calculate_alignment_statistics` function calculates and prints overall quality metrics, such as the total number of mutations, Scaled Sum-of-Pairs (SP) Score, and Entropy.
6. **Alignment Reconstruction and Output:**
   - The `save_as_sparse_npz` function is always called. It reconstructs a virtual alignment template based on all observed insertions and then converts the difference information into the sparse column-based format before saving it as a compressed `.npz` file.
   - If the total number of sequences is 40,000 or less, the `reconstruct_and_write_fasta` function is also called. It uses the same alignment template to reconstruct the full, aligned sequence strings for every entry and writes them to the output FASTA file.

**Usage** This script can be run from the command line.

**Command Format:**

```
python3 SR_Align.py -i <input_file.fasta> -o <output_directory/> -t <num_threads>
```

**Arguments:**

- `-i, -input`: (Required) Path to the input FASTA file. The first sequence in this file will be used as the reference.
- `-o, -output`: (Required) Path to the output directory. The script will create this directory if it does not exist. Output files will be named based on the input file's name (e.g., 'input.fasta' will produce 'input.npz' and 'input.fasta' inside the specified directory).
- `-t, -threads`: (Optional) The number of CPU threads to use for parallel alignment. Defaults to all available CPU cores.

**Example:**

```
python3 SR_Align.py -i sequences.fasta -o results/ -t 16
```

This command will align all sequences in `sequences.fasta` against the first sequence, using 16 threads. The results will be saved in the `results/` directory as `sequences.npz` and, if there are 40,000 or fewer sequences, also as `sequences.fasta`.

#### Alignment Quality Calculator: A Memory-Efficient Parallel Scorer

The `alignment_quality_calculator.py` script is a command-line tool designed to efficiently calculate quality metrics for massive MSAs stored in FASTA format. To handle files that are too large to fit into memory, the script employs a parallel, stream-based processing strategy.

**Core Features and Workflow** The script's workflow is optimized for both speed and memory efficiency:

1. **Automatic Format Detection:** The script checks the input file's extension. If it is `.npz`, it uses a fast, single-threaded method to read the pre-computed sparse data. For any other extension, it defaults to a parallel, stream-based processing strategy suitable for large FASTA files.

2. **Parallel FASTA Processing:** For FASTA files, the script reads the alignment in sequential chunks (e.g., 1000 sequences at a time) to keep memory usage low. It distributes these chunks to a pool of worker processes, where each worker computes a partial "counts matrix" (tabulating character frequencies per column). These partial matrices are then aggregated by the main process into a final, global counts matrix.

3. **Direct NPZ Calculation:** For NPZ files, the script directly reconstructs the character counts for each column from the sparse format. This process is computationally inexpensive and is performed in a single thread, as parallelization offers no advantage.

4. **Unified Final Calculation:** Regardless of the input format, once the character counts for each column are determined, the script uses a single, shared function to calculate the final quality scores. The metrics calculated are:

- **Scaled Sum-of-Pairs (SP) Score:** This score is computed based on the aggregated counts for each column. The pairwise score for each character combination is defined by a unified scoring matrix with the following rules: a match between identical bases (e.g., A vs. A) receives a score of +1; a mismatch between different bases (e.g., A vs. C) receives a score of -1; an alignment between any base and a gap receives a score of -2; and an alignment between two gaps receives a score of 0. Degenerate bases are decomposed into their constituent standard bases, and their counts are distributed proportionally for scoring.
- **Total Negative Entropy:** The Shannon entropy is calculated for each column based on character frequencies, and the total entropy is the sum across all columns. The negative of this value is reported.

#### Usage

##### Command Format:

```
python3 alignment_quality_calculator.py <input_file.fasta> [num_workers]
```

##### Arguments:

- `input_file`: (Required) Path to the input FASTA alignment file.
- `num_workers`: (Optional) The number of parallel processes to use. Defaults to the total number of available CPU cores.
- `-chunk_size`: (Optional) The number of sequences to be processed in a single batch by each worker. Default is 1000.

##### Example:

```
python3 alignment_quality_calculator.py large_alignment.fasta 16
```

### S4 Parameter Handling in PHMM Training (M-Step)

This section details the M-step (Maximization) of the Baum-Welch algorithm, which re-estimates the Profile HMM (PHMM) parameters (see Figure S5 for PHMM architecture). This step uses the expected values (e.g., expected transition counts, expected emission counts) calculated during the E-step (Expectation). All calculations are performed in log-space to maintain numerical stability.

The update strategy for each parameter falls into one of two categories, which serves to regularize the model and prevent overfitting:

1. **Expectation + Pseudocount:** The parameter is re-estimated by adding a log-space **pseudocount** to the E-step's log-space expectation, followed by normalization.
2. **Fixed Value:** The parameter is reset to a **fixed value**, completely ignoring the E-step expectation. This is used to enforce a specific model behavior, such as uniform probabilities for insertions or deletions, preventing the model from learning complex, dataset-specific indel profiles.

**Match State Emission Probabilities (Me_Matrix)** These probabilities are updated using **E-step expectations + pseudocounts**.

- **Method:** The initial log probability is computed from the E-step's expected observations (gamma_o_M) and expected state visits (MEi). A log-space pseudocount (self.emProbAdds_Match) is added to the log-space probabilities for the four standard bases (A, T, C, G).
- **Normalization:** The resulting values for each Match state are normalized (by subtracting the log-sum-exp of all emissions for that state) to ensure the probabilities sum to 1.

**Insert State Emission Probabilities (Ie_Matrix)** These probabilities are updated using a **fixed value**.

- **Method:** E-step expectations are ignored. All insert state emission probabilities are uniformly reset to $\log(0.25)$.
- **Rationale:** This enforces the assumption that insertions are random and not position-specific, a common simplification in PHMMs.

**Delete State Transition Probabilities (D2M_array, D2D_array)** These probabilities are updated using a **fixed value**.

- **Method:** E-step expectations are ignored. Both D$\rightarrow$M and D$\rightarrow$D transition probabilities are uniformly reset to $\log(0.5)$.

**Match State Transition Probabilities** These are updated using **E-step expectations + pseudocounts**. The update is split into two parts:

- **Core States (M[0...N-1]):** The expected log-space transitions for M$\rightarrow$M, M$\rightarrow$I, and M$\rightarrow$D are retrieved. Log-space pseudocounts of -3, -50, and -4 are added, respectively. The three paths are then normalized.
- **Final State (M[N]):** The expected log-space transitions for M$\rightarrow$E (End) and M$\rightarrow$I (final insertion) are retrieved. Log-space pseudocounts of -10 and -10 are added, respectively, followed by normalization.

**Insert State Transition Probabilities** These are updated using **E-step expectations + pseudocounts**. The update is handled in segments (start, middle, and end).

- **Method:** For all segments, a log-space pseudocount of -1 is added to the E-step expectations for both the I$\rightarrow$M and I$\rightarrow$I transitions. The transition to the End state (I$\rightarrow$E) also receives a pseudocount of -1.
- **Normalization:** The probabilities are normalized at each segment.

**Initial State Probabilities (pi_M, pi_I, pi_D)** These are updated using **E-step expectations + pseudocounts** in a two-stage process:

1. The E-step expectations (gamma values) for the start node are retrieved and normalized to get an initial estimate.
2. Log-space pseudocounts of -1, -1, and -1 are added to the pi_M, pi_D, and pi_I estimates, respectively. The probabilities are then re-normalized to produce the final values.

**Degenerate Base Emission Matrices** After the primary emission matrices (`Me_Matrix`, `Ie_Matrix`) are updated, corresponding matrices for degenerate bases are generated for use in downstream steps.

- **Standard Bases (A, T, C, G):** Probabilities are copied directly from the updated matrices.
- **Degenerate Bases (R, Y, N, etc.):** The probability for a degenerate base is calculated as the log-space average of the probabilities of its constituent standard bases (e.g., $P(R) = \log(\frac{\exp(P(A))+\exp(P(G))}{2})$).

**Summary of Numerical Values** The specific values used for regularization are summarized in Table S1 (Fixed Values) and Table S2 (Pseudocounts).

**Table S1: Fixed Values for Parameter Updates.** These parameters ignore E-step expectations and are reset to these log-space values in each M-step.

| Parameter | Log-Space Value | Note |
|---|---|---|
| Insert Emission (`Ie_Matrix`) | $\log(1/4)$ | Uniform probability (0.25) for A, T, C, G. |
| Delete Transition (`D2M_array`) | $\log(1/2)$ | D$\rightarrow$M transition probability set to 0.5. |
| Delete Transition (`D2D_array`) | $\log(1/2)$ | D$\rightarrow$D transition probability set to 0.5. |

**Table S2: Log-Space Pseudocounts Added to E-Step Expectations.** These values are added to the log-space E-step expectations before normalization.

| Parameter | Transition/Emission | Log-Space Pseudocount |
|---|---|---|
| Match Emission | M $\rightarrow$ A/T/C/G | `self.emProbAdds_Match` (variable) |
| Match Transition | M $\rightarrow$ M (Core) | -3 |
| | M $\rightarrow$ I (Core) | -50 |
| | M $\rightarrow$ D (Core) | -4 |
| | M $\rightarrow$ E (Final) | -10 |
| | M $\rightarrow$ I (Final) | -10 |
| Insert Transition | I $\rightarrow$ M (All) | -1 |
| | I $\rightarrow$ I (All) | -1 |
| | I $\rightarrow$ E (Final) | -1 |
| Start Probability | Start $\rightarrow$ M | -1 |
| | Start $\rightarrow$ I | -1 |
| | Start $\rightarrow$ D | -1 |

#### S5 Experimental Environment and Parameter Settings

**Hardware Environment** All benchmark tests were conducted on two different server machines:

- **Machine A:** Used for all tests except for the ultramassive COVID test set.
  - **CPU:** 2 x Intel(R) Xeon(R) Platinum 8375C @ 2.90GHz (Total 64 cores / 128 threads)

  - **Memory (RAM):** 512 GB
  - **Operating System:** Ubuntu 22.04.5 LTS

- **Machine B:** Used specifically for the ultramassive COVID test set (million-scale sequences).
  - **CPU:** 2 x Intel(R) Xeon(R) Gold 6330 CPU @ 2.00GHz (Total 56 cores / 56 threads)
  - **Memory (RAM):** 1.0 TB
  - **Operating System:** Ubuntu 22.04.3 LTS

##### Software Parameter Settings

- **DAG-align:**
  - **Standard Tests:** Run using the automatic mode (`-a`). In this mode, the user only specifies the input and output paths. The program automatically calculates the k-mer diversity ratio ($R_k$), defined as:

$$R_k = \frac{\text{Number of unique k-mers}}{\text{Total number of k-mers}}$$

    When $R_k > 0.9$, the fragment length $L_{frag} = 16$ is automatically selected; otherwise, $L_{frag} = 32$ is used. Additionally, based on the average sequence length, the program automatically estimates and sets the number of sequences per subgraph using a built-in function.
  - **Ultramassive COVID Test Set:** Run using the `-l` mode. This mode defaults to using a fragment length of $L_{frag} = 32$ and a subset size (subgraph size) of $5000$ sequences for parallel graph construction.
- **Other Compared Software:** All other software used for comparison (including MAFFT, Halign4, SR-align, Muscle v3, Muscle v5) were run using their respective default settings or automatic modes, without any additional parameter tuning.

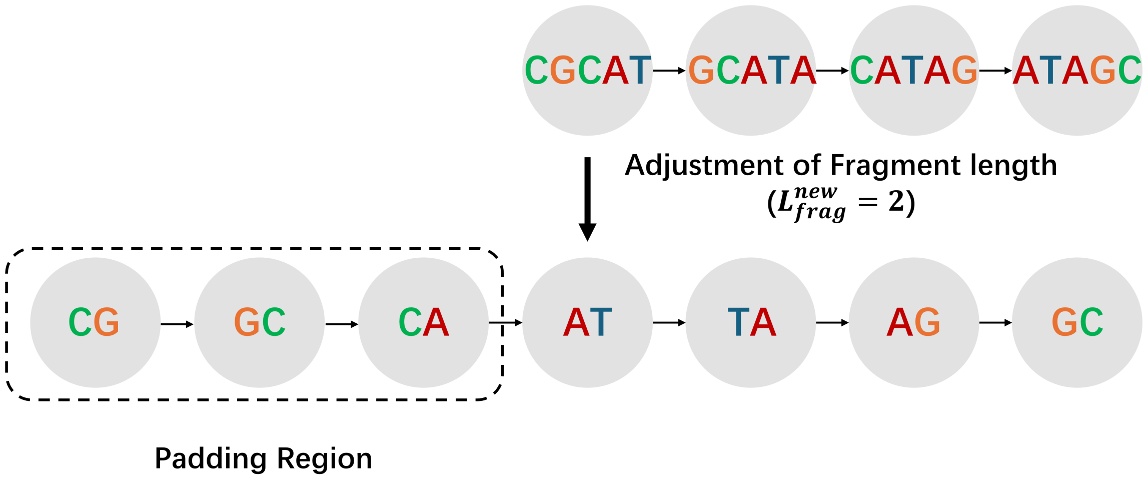


**Figure S4. Schematic illustration of the Graph Resolution Adjustment Process**. The figure demonstrates the procedure for increasing the graph's resolution by reducing the fragment length within nodes, shown for a target length $L_{frag}^{new} = 2$. The process consists of (1) truncating the sequence fragments in all existing nodes to their target length $L_{frag}^{new} = 2$, and (2) prepending new nodes to the start of the graph to restore sequence information lost at the 5' ends.

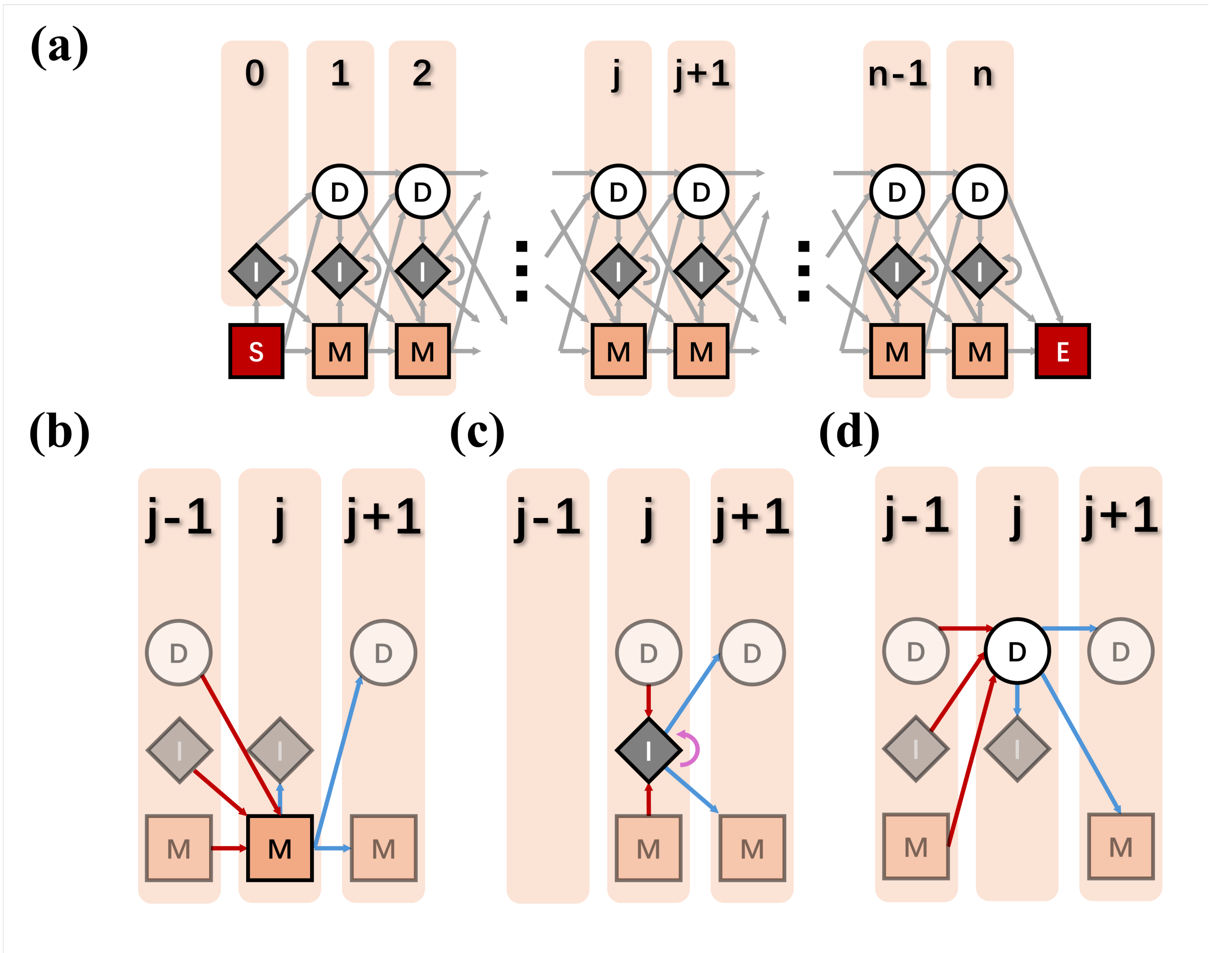


**Figure S5. The Canonical State Architecture of a Profile Hidden Markov Model (PHMM).** The figure illustrates the standard topology of a PHMM used for sequence alignment. The model has a start position ($S$), an end position ($E$), and a fixed number of state positions, each of which for a specific sequence may be in one of three states: match ($M$), insert ($I$), and delete ($D$). (a) An overview of the $S$, $E$, and regular state positions, and the allowed transitions between them. The model progresses from left to right, each state position corresponds to a column of certain conservation in a MSA. Minority sequences have no occupation of a given state position has state $D$ on that position. Letters in a sequence can not be attributed to any state position have a state $I$ based on its nearest preceding letter having a state position. (b-d) Detailed views showing the specific set of preceding and succeeding states for a state position in $M$ state (b), $I$ state (c), and $D$ state (d), respectively. This structure allows the model to score matches, mismatches, insertions, and deletions in a probabilistic framework..

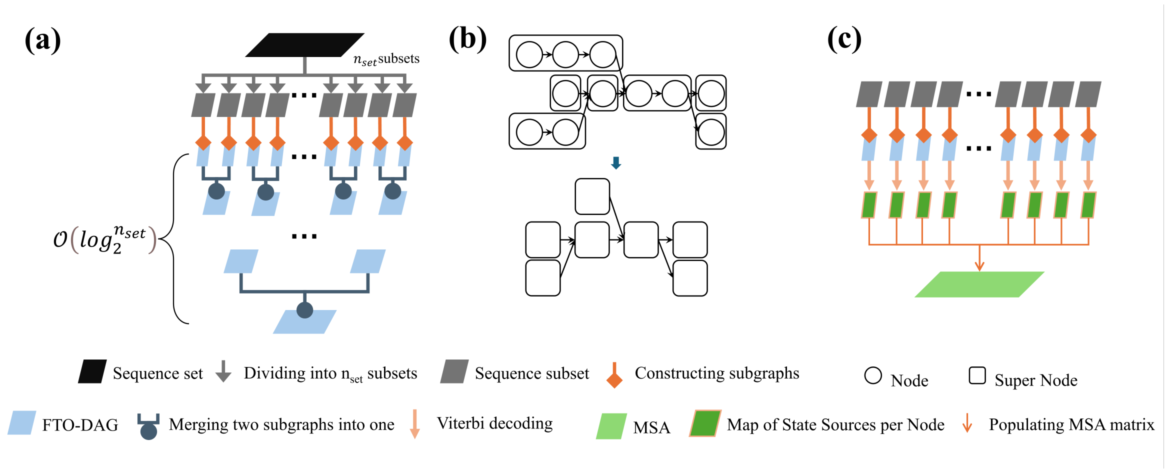


**Figure S6. Parallelization Strategies for various stages of ultra-massive MSA**. (a) Parallel construction of FTO-DAG. The ultra-massive genome set is first divided into $n_{set}$ subsets. After a subgraph is constructed for each subset in parallel, two-to-one merging is performed approximately for $\mathcal{O}log_2^{n_{set}}$ rounds to obtain the whole graph, the number of rounds $\mathcal{O}log_2^{n_{set}}$ is based on the fact that by starting with $n_{set}$, after the first round of merging, we have $\sim \frac{n_{set}}{2}$ graphs. After the second round, we have $\sim \frac{n_{set}}{4}$ graphs, etc. (b)Parallel training. Nodes in each branchless path in the FTO-DAG are organized as a single training task (Super node in the figure). All tasks are listed in a queue with topological dependence, shown as directed edges in the bottom figure. A task may start training only after its preceding tasks are done. Different tasks without topological dependence may be trained in parallel. (c) Parallel decoding. The optimal alignment paths for individual subgraphs are decoded in parallel and then assembled into the final global MSA. In each paragraph, the same TO based parallel queue strategy may be utilized when necessary, except that the reverse TO is utilized for decoding.

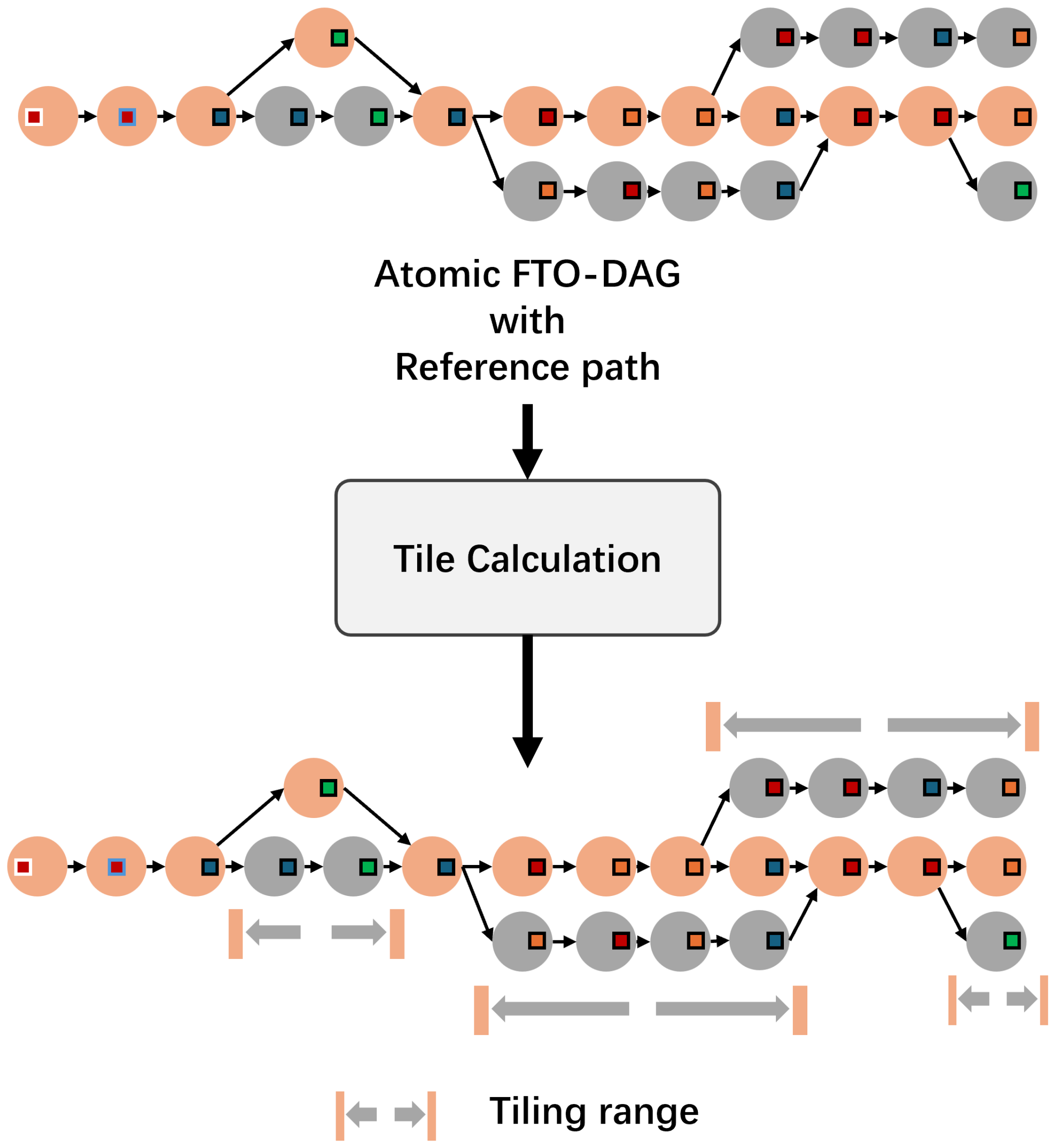


**Figure S7. Illustration of the Tiling Strategy for Reducing Computational Complexity**, related to STAR Methods. Top: Without tiling, probabilities for matching all PHMM state positions must be computed for each node. Bottom: With tiling, computation for each node is restricted to a narrow region of PHMM state positions (the "tile"), drastically reducing complexity. In actual computation, a buffer is added to both directions of a tile for each node to ensure negligible probability leak.

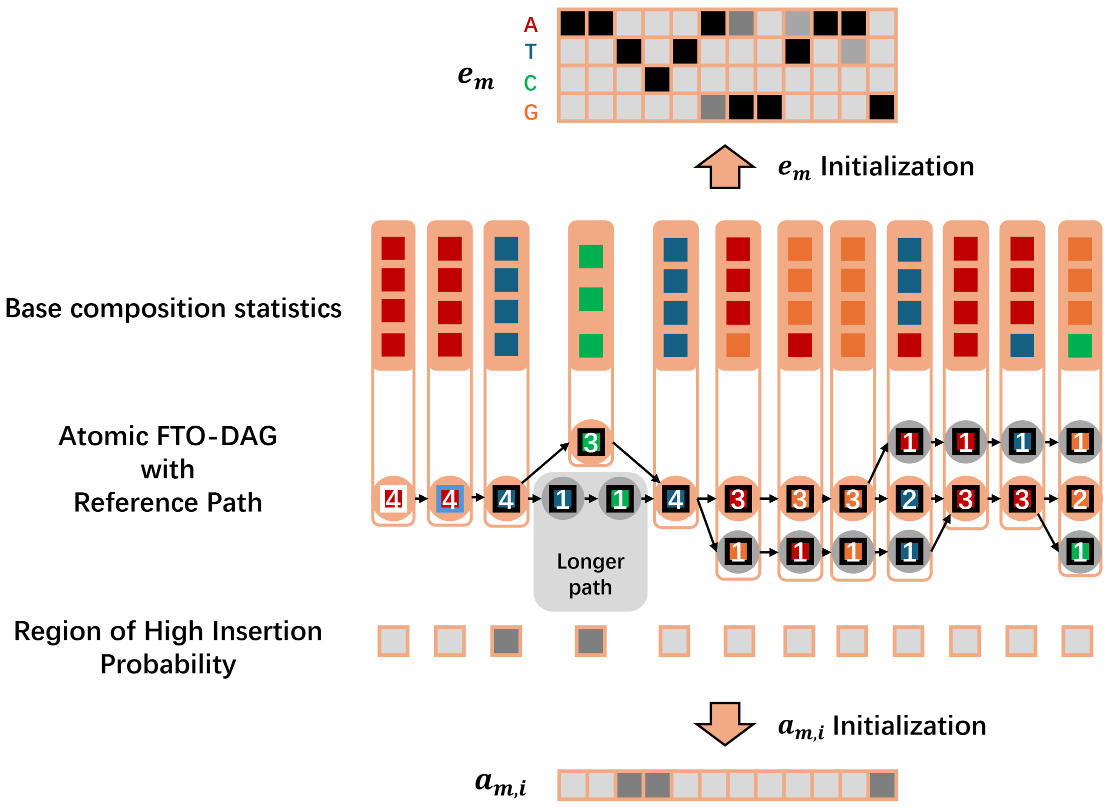


**Figure S8. Graph-guided Initialization of PHMM Parameters**, The figure illustrates the two main stages of parameter initialization based on the graph topology after a reference path has been identified. Nodes on the reference path are shown in orange and other nodes are in grey. (Top) Initialization of match state emission probabilities ($E_M$). The emission profile for a state position ( corresponding to a reference path node ) is calculated from the weighted base frequencies of itself and all topologically corresponding nodes on "parallel paths" (bypass partial graphs of equal length, see STAR Methods, DAG-TPHMM and DAG-Viterbi Algorithm Sets, Graph-guided DAG-TPHMM Parameter Initialization). (Bottom) Initialization of Match-to-Insert transition probabilities ($a_{M \to I}$). When a bypass subgraph is longer than the reference segment it spans, it is identified as a potential insertion hotspot, and the transition probabilities for state positions corresponding to that region of the reference path are correspondingly increased. This entire process ensures a biologically meaningful starting point for model training.

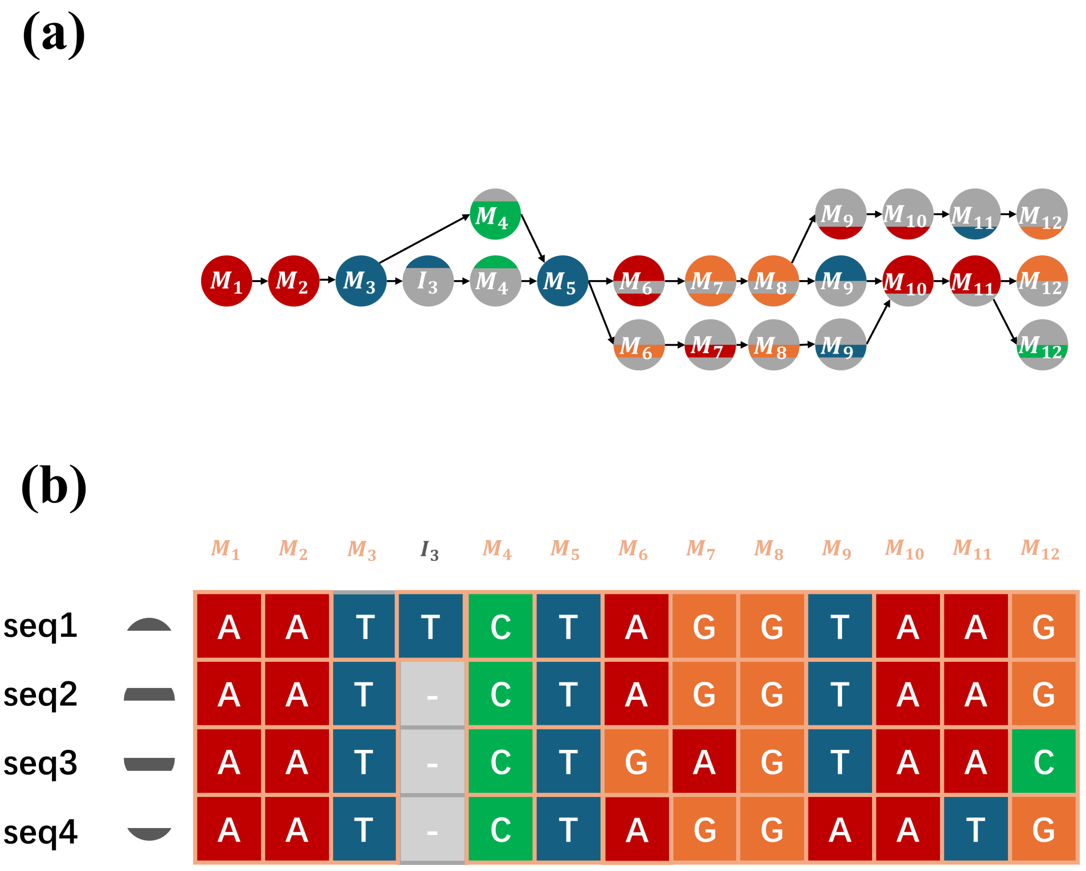


**Figure S9. Viterbi Decoding and Path Reconstruction on the Graph**. The figure illustrates the final stages of the alignment process. (a) Optimal state and state positions as subscript. Other information of the 'FinalOptimalStateMaps' data structure (e.g. sequence IDs) is not shown explicitly. It is important to note that the subscript (e.g. $_4$ in $M_4$) in each node represents state position rather than node index or TC. (b) The final MSA (MSA) matrix, which is populated directly and efficiently from the 'FinalOptimalStateMaps' structure, bypassing the need to reconstruct individual full-length paths.

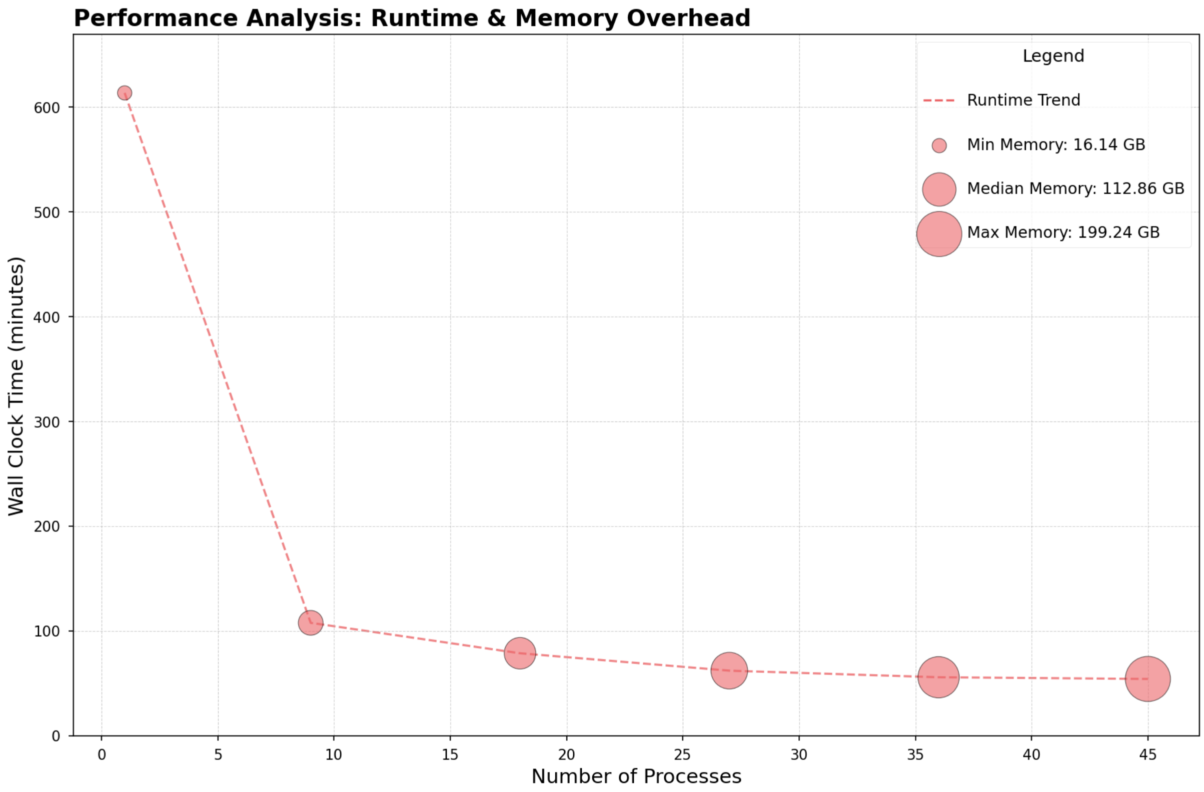


**Figure S10. Performance of Parallelized Graph Construction**. The runtime and peak memory usage for the graph construction phase were measured on a dataset of 500,000 SARS-CoV-2 sequences(the first 500,000 sequences from the largest dataset), using an increasing number of parallel processes (1, 9, 18, 27, and 36). The results demonstrate effective scaling, with runtime decreasing significantly as more processes are added, while memory usage remains stable.

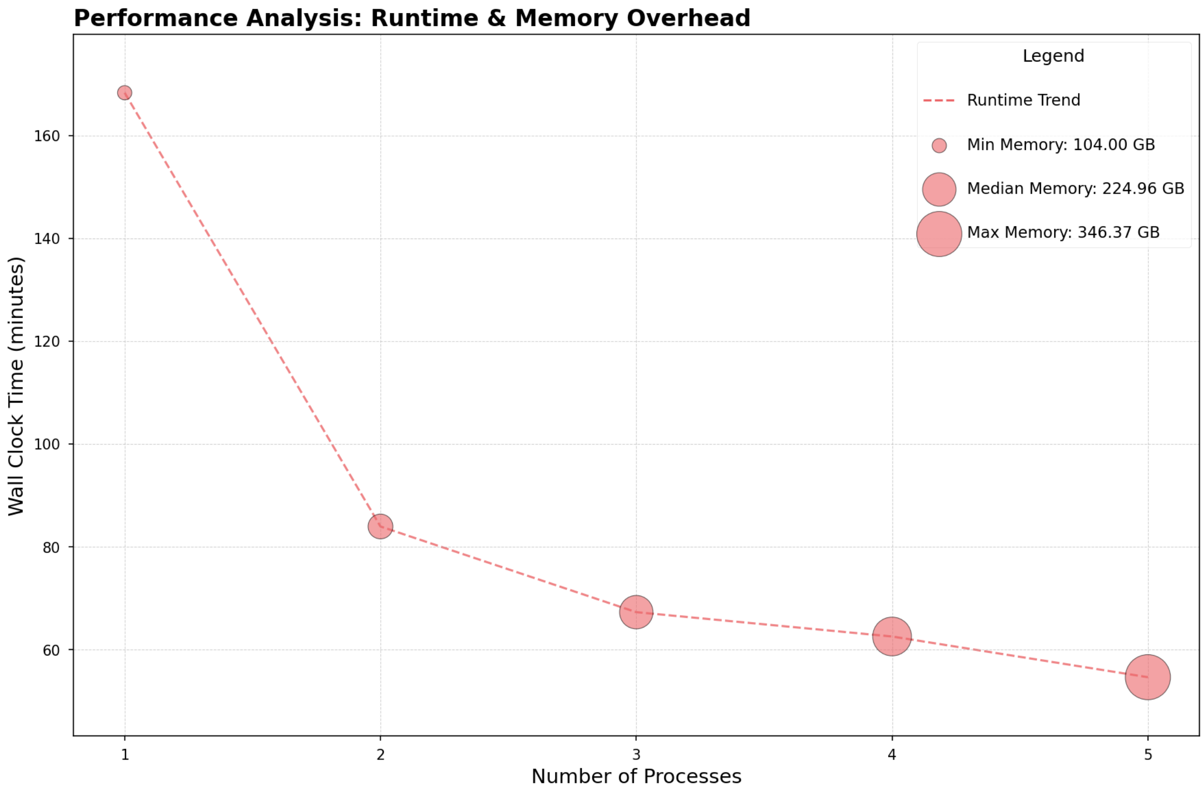


**Figure S11. Performance of Parallelized Viterbi Decoding**. The runtime and peak memory usage for the Viterbi decoding phase were measured on the same 500,000 SARS-CoV-2 sequence dataset(the first 500,000 sequences from the largest dataset), using an increasing number of parallel processes. The decoding phase also exhibits strong scalability, with a substantial reduction in runtime as the number of processes increases.

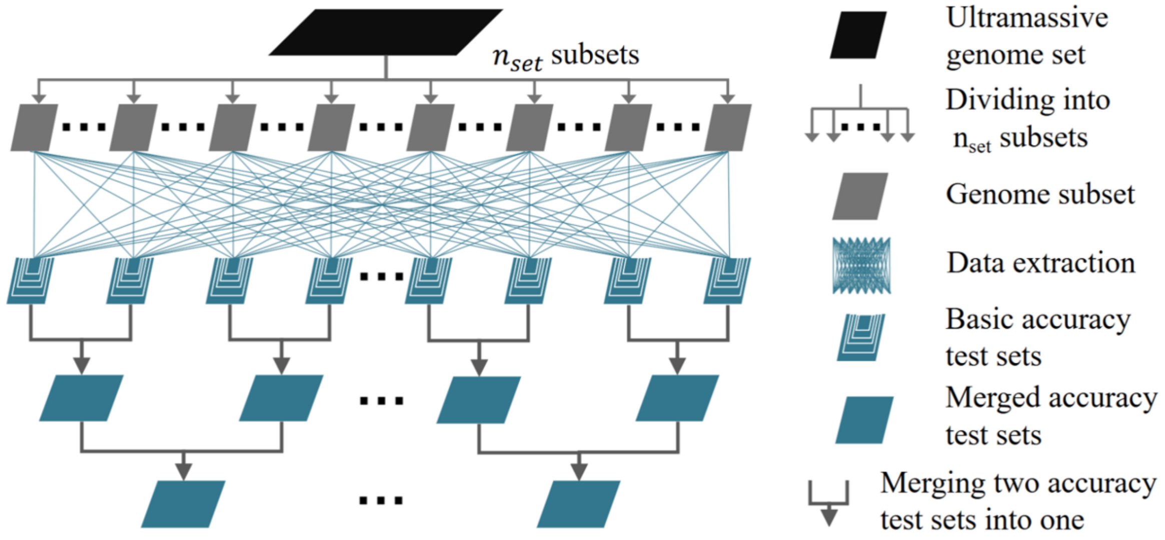


**Figure S12. Schematic for the Construction of the Number-Tiered Benchmark Dataset**, related to STAR Methods. This diagram depicts a hierarchical process for constructing a multi-tiered test set from an initial, very large genome collection. The process unfolds in three main stages: Partitioning, Sampling, and Hierarchical Merging, allowing for the creation of progressively larger datasets for scalability analysis.

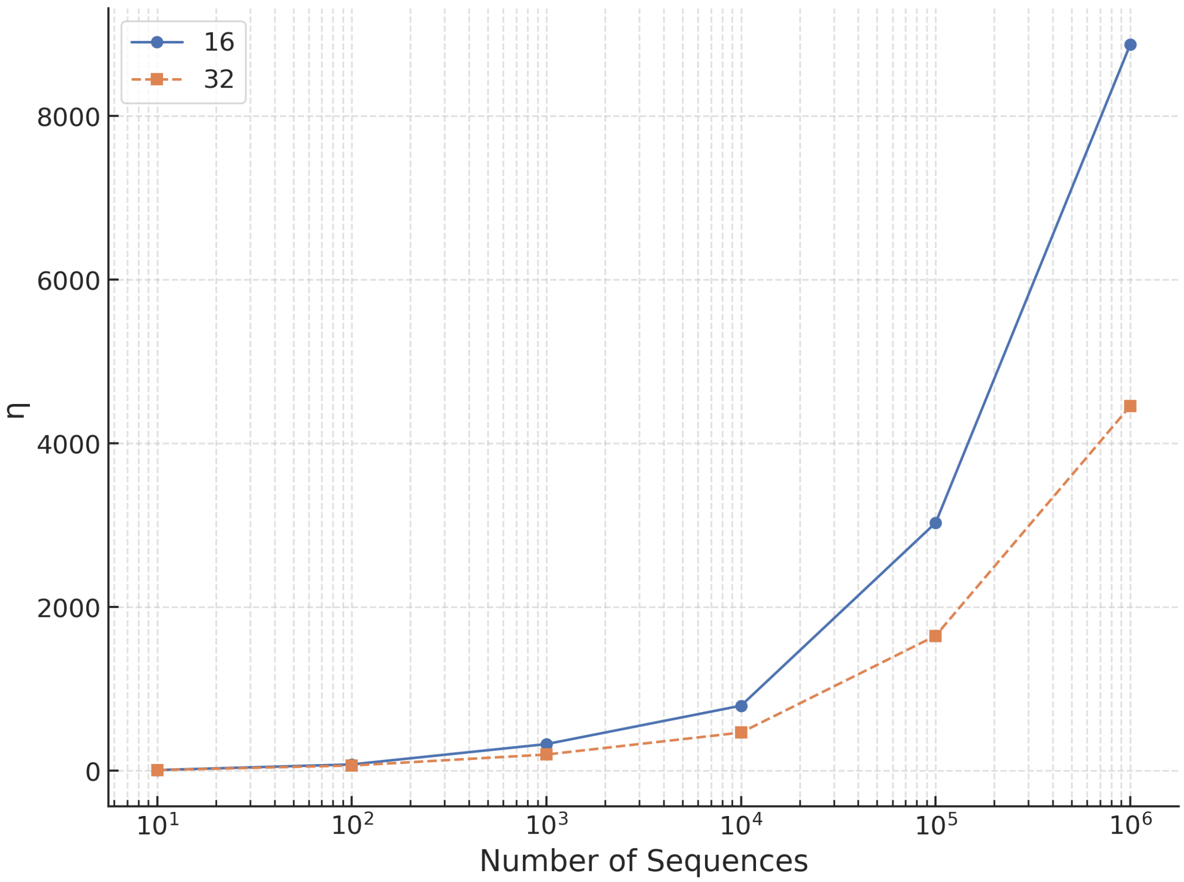


**Figure S13. The compression ratio for two different fragment lengths $L_{frag} = 16$ and $32$.**

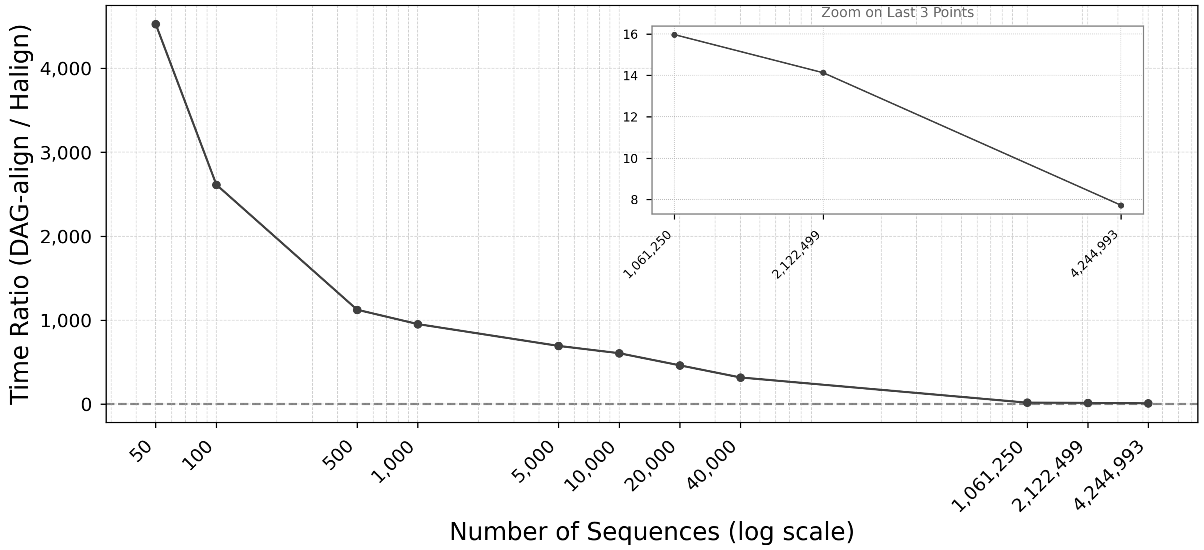


**Figure S14. Ratio between the CPU clock time of DAG-align and Halign4 on various sized SARS-CoV2 genome set.** The observations suggest DAG-align scales better than Halign with increasing data set size, This is consistent with the increase of compression ratio with increasing data set shown in Figure S13. At the level of $\sim 4$ million the ratio is below 8, which is essentially comparable algorithmically considering the programming language difference (Python for DAG-align and C++ for Halign).

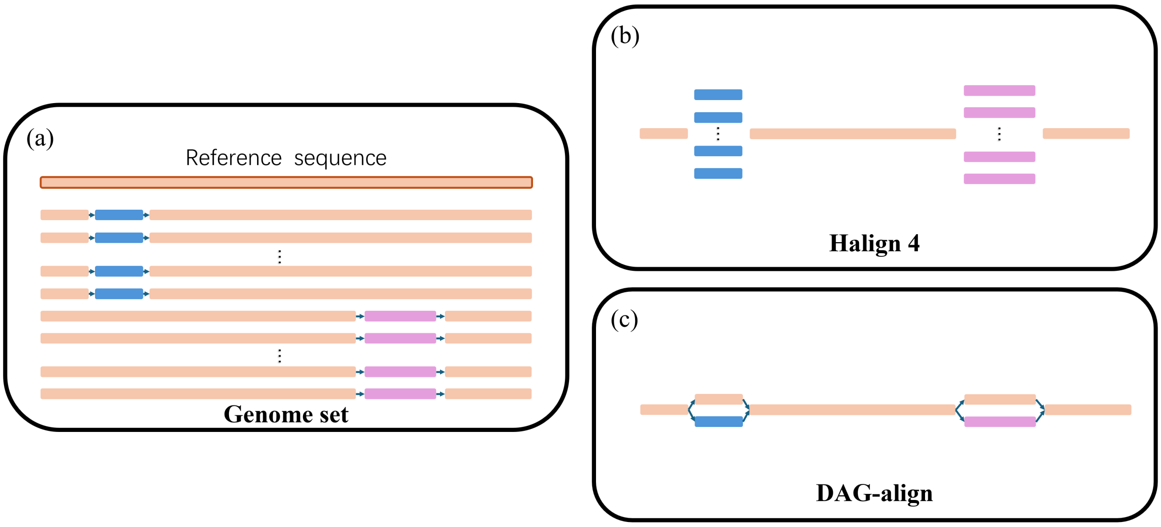


**Figure S15. Schematic illustration of the repetition elimination in Halign and DAG-align.** (a) A reference sequence (the top one) and a large number of genomes in a set. For non-reference sequences, some share a blue segment that is different from the reference, the others share a purple segment that is different from the reference. (b) Halign only aligns different segments to the reference, thus realizes significant repetition elimination. However, repetition of shared identical segments in non-reference sequences remains. (c) DAG-align further eliminates such repetitions. Of course, such repetition elimination comes with cost of expensive graph data structures, which pays off for ultra-massive datasets.
